# Climate-driven mangrove dieback and recovery: a case study in Albert and Leichhardt Rivers, Australia

**DOI:** 10.1101/2025.03.30.646221

**Authors:** Rogerio Victor S. Gonçalves, Emma Asbridge, Jeffrey J. Kelleway, Richard Lucas, Kerrylee Rogers

## Abstract

A severe dieback event occurred in 2015/2016 in the Gulf of Carpentaria, northern Australia, which was later attributed to El Niño-Southern Oscillation (ENSO) and unusually low sea levels associated with the lunar nodal cycle. However, the long-term drivers of mangrove changes, their resilience, and species-specific responses to these environmental events remain insufficiently understood. This study applies an integration of k-means clustering and random forest classification on the Landsat archive to address knowledge gaps regarding mangrove dynamics. Despite episodic dieback in 2002/2004 (9.1 km^2^), 2008 (2.9 km^2^), and 2015/2016 (8.1 km^2^), mangrove zones showed a net increase in area (+260%, from 21.4 km^2^ to 55.7 km^2^) and greenness (NDVI, +18%) over the period 1987-2023. These results indicate cyclical patterns of decline in extent and condition followed by rapid recovery. Landward edges, dominated by *Avicennia marina*, presented the lowest NDVI values during extreme negative ENSO phases, particularly when unusually low sea levels driven by the lunar nodal cycle further reduced tidal inundation. This pattern suggests greater vulnerability to water deficits and salinity stress in these zones. The mature *R. stylosa* dominated areas in the central zone of the mangrove forest remained stable throughout the study period, and significant progradation was observed at the seaward margins, dominated by *A. marina*, likely driven by sediment accretion during increased river discharge years. This highly dynamic mangrove forest demonstrates capacity to extend both landwards and seawards. However, as climate variability intensifies, particularly with more frequent extreme events and shifting tidal regimes projected with sea-level rise, understanding the long-term resilience of mangrove forests and their capacity to recover will be crucial for informing conservation strategies focused on specific zones and maintaining blue carbon stocks of mature mangrove forests.

## 1. Introduction

Changes in the condition and extent of mangrove forests globally are increasingly linked to climate variability, which can pose a considerable risk to their ecosystem services. Ocean-atmosphere interactions, such as the El Niño Southern Oscillation (ENSO) and Indian Ocean Dipole (IOD) modify precipitation, ocean and air temperatures (Luo et al. 2010; Cai et al. 2021). When these conditions exceed the physiological limits of mangroves, dieback and declines in condition can occur (Lovelock et al. 2017; Abhik et al. 2021; Carruthers et al. 2024). Climate variability effects may intensify under overlapping perturbations, such as ENSO and low water stages of the lunar nodal cycle (Saintilan et al. 2022). These impacts are primarily driven by physical disturbances, such as cyclone-induced windthrow (Asbridge et al. 2018), and environmental stressors, including water deficits leading to desiccation (Asbridge et al. 2015) or prolonged waterlogging (Asbridge et al. 2024). In 2015/2016, the Gulf of Carpentaria (hereafter the Gulf), northern Australia, experienced significant and widespread dieback (Duke et al. 2017). At the time, this was largely attributed to a combination of elevated temperatures and reduced precipitation due to a negative ENSO phase (Sippo et al. 2020; Abhik et al. 2021). Associated with this, low tidal stages associated with the lunar nodal cycle may have intensified the effects of ENSO, particularly on mangrove vegetation at higher elevations in the tidal frame (Saintilan et al. 2022). Before the dieback event, mangroves in the Gulf were observed to be highly dynamic with dieback, landward extension and seaward progradation all observed, with an overall net increase in mangrove area from 1987 to 2014 (Asbridge et al. 2015).

The vertical distribution of mangrove species within the intertidal zone, often referred to as mangrove zonation (Snedaker 1982), has been described across the globe (Bunt 1996; Castañeda-Moya et al. 2006; Ximenes et al. 2016; Irawan et al. 2021; Lombard et al. 2023) and is influenced by species-specific physiological tolerances to environmental conditions, including salinity, elevation, substrate, and inundation frequency (Ellison et al. 2000). *Rhizophora stylosa*, for instance, is usually found in lower elevations within the tidal frame due to its tolerance of prolonged inundation with this facilitated by an ability to maintain lower water potentials and regulate water uptake under saline conditions (Reef and Lovelock 2015). *Avicennia marina*, on the other hand, occupies higher intertidal areas, but also exhibits high salinity tolerance through salt exclusion and secretion mechanisms that minimise water loss (Reef and Lovelock 2015) and may support absorption of atmospheric moisture (Coopman et al. 2021). These species-specific adaptations highlight the dynamic nature of mangrove zonation patterns across regions and the importance of long-term monitoring to assess how species respond to climate anomalies over time.

Dense time series analysis from optical satellites sensors (e.g., on board the as Landsat and Sentinel satellites) are widely used to monitor vegetation changes over time, enabling the detection and quantification of shifts in mangrove distribution and condition across large geographic areas (e.g., Pasquarella et al. 2016; Lu and Wang 2021; Bunting et al. 2022, 2023). While Sentinel offers higher spatial resolution (10-20 m), its temporal coverage (2015–present) is more limited than the Landsat archive (1987–present). The extended Landsat record therefor preserves a long-term dataset from which multiple climatic cycles, including ENSO (4–7 years), the IOD (typically 3–5 years), and the lunar nodal cycle (18.6 years), can be discerned. As a consequence, a more comprehensive assessment of how mangrove ecosystems respond to cyclical environmental variability can be observed. While previous research using remote sensing approaches has focused on the short-term impacts of climate cycles on mangrove vegetation (Alongi 2015; Duke et al. 2017), providing limited information on their effects, long-term analyses integrating remote sensing and climate data remain scarce (Asbridge et al. 2016; Abhik et al. 2021; Hickey et al. 2021; Maina et al. 2021). Despite previous research, the long-term environmental and climatic drivers of mangrove dynamics, as well as their species-specific responses, remain insufficiently understood. This is particularly evident when zones that are visibly distinct, largely because of their dominance by mangrove genera, such as *Avicennia* sp. and *Rhizophora* sp., respond to environmental factors, including sea-level fluctuations driven by the ENSO, IOD and lunar nodal cycle. Understanding these species-specific responses is essential for identifying vulnerable zones and developing conservation strategies focused on the most threatened areas.

In this research, key gaps in understanding long-term mangrove dynamics were addressed by examining changes in the extent and condition of mangroves at the Albert and Leichhardt Rivers, Gulf of Carpentaria, Australia, over 36 years (1987 to 2023). The primary aim was to identify where mangrove change have occurred and the environmental factors that might have contributed. Specifically, the objectives were to: (1) describe and quantify changes in the extent and condition of different mangrove zones over time, and (2) determine the likely environmental drivers influencing these changes, including climatic and tidal cycles and geomorphological processes. To achieve this, mangrove zones were classified using Landsat satellite archive (1987 – 2023) and integrated information on mangrove changes in health (NDVI) and extent with climatic indices, including ENSO and IOD, and tidal height data. This study is critical for identifying and monitoring vulnerable mangrove zones and informing targeted management strategies under the uncertainty of long-term environmental changes and climate variability.

## 2. Methods

### 2.1. Study area

The Albert and Leichhardt Rivers drain into the Gulf of Carpentaria, Queensland, Australia (17.57° S, 139.80° E, Figure 1). The region experiences a semi-arid climate, characterised by a dry season from April to November and a wet season from December to March (BOM 2024b). The mean minimum and maximum annual temperatures range from 13°C to 35°C (BOM 2024b). Tides at the mouth of the Albert and Leichhardt Rivers are diurnal mesotidal, with an average range of 2–3 m (Whitfield and Elliott 2011). In the Gulf, *Rhizophora stylosa* is usually found in the central intertidal zone (Hay 2009; Asbridge et al. 2016), *Avicennia marina* dominates the seaward edge but is also present in landward zones, along with *Ceriops* sp. (Hay 2009). Other species such as *Aegialitis annulata* and *Xylocarpus mekongensis* were also identified in the area, but in a limited distribution within mixed species areas.

**Figure 1.**
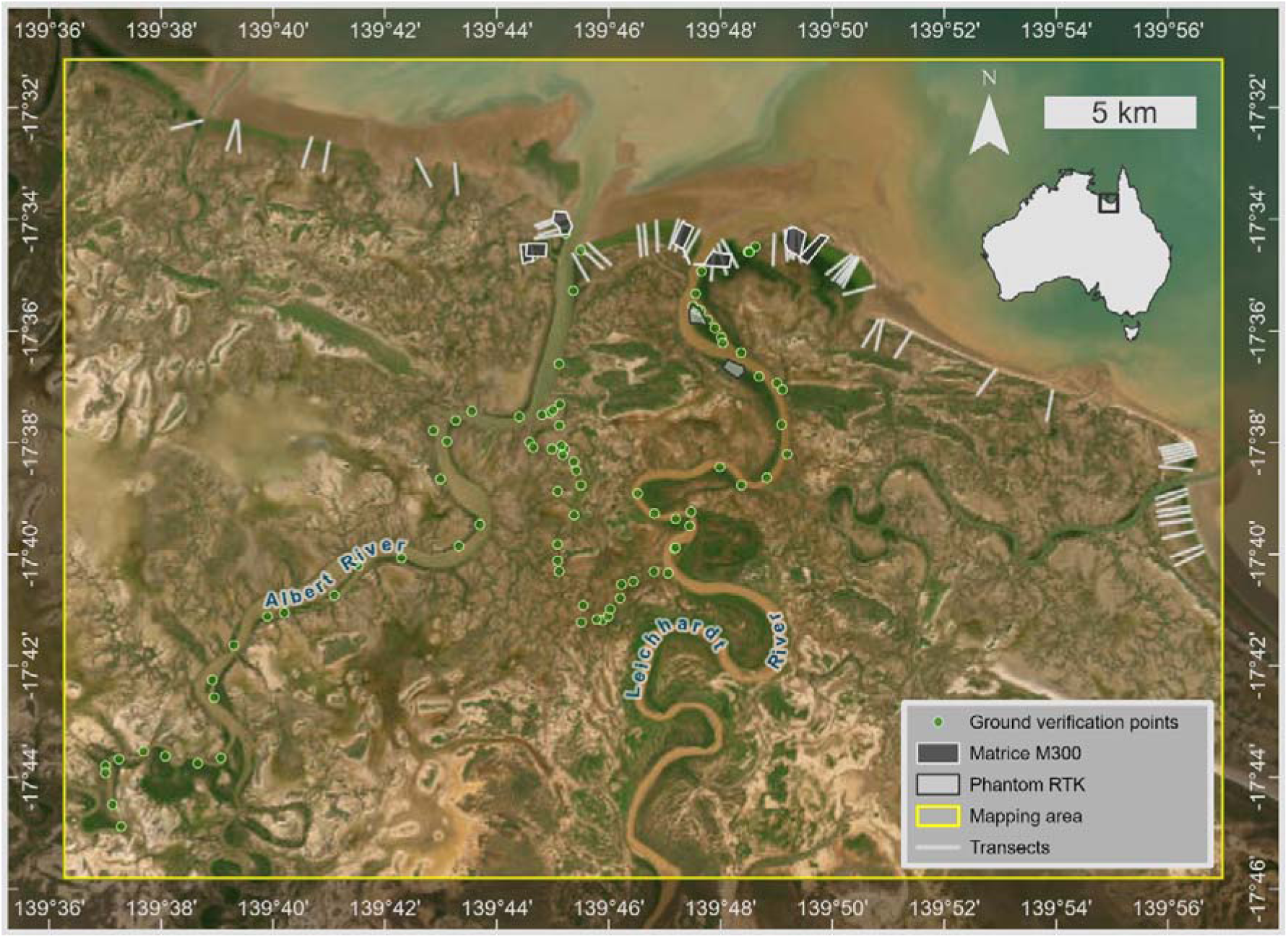
Study area focusing on Albert (west) and Leichhardt (east) River mouths. Ground verification data collected from the boat are represented by green points and remotely piloted aircraft flight plans correspond to dark and light grey polygons for Matrice M300 and Phantom 4 RTK.

### 2.2. Approach

Figure 2 shows the workflow approach for mapping changes in mangrove extent and conditions using Digital Earth Australia sandbox, including the classification model, high-resolution remotely piloted aircraft (RPA) spectral imagery and LiDAR data processing, mangrove condition assessment, and model validation.

**Figure 2.**
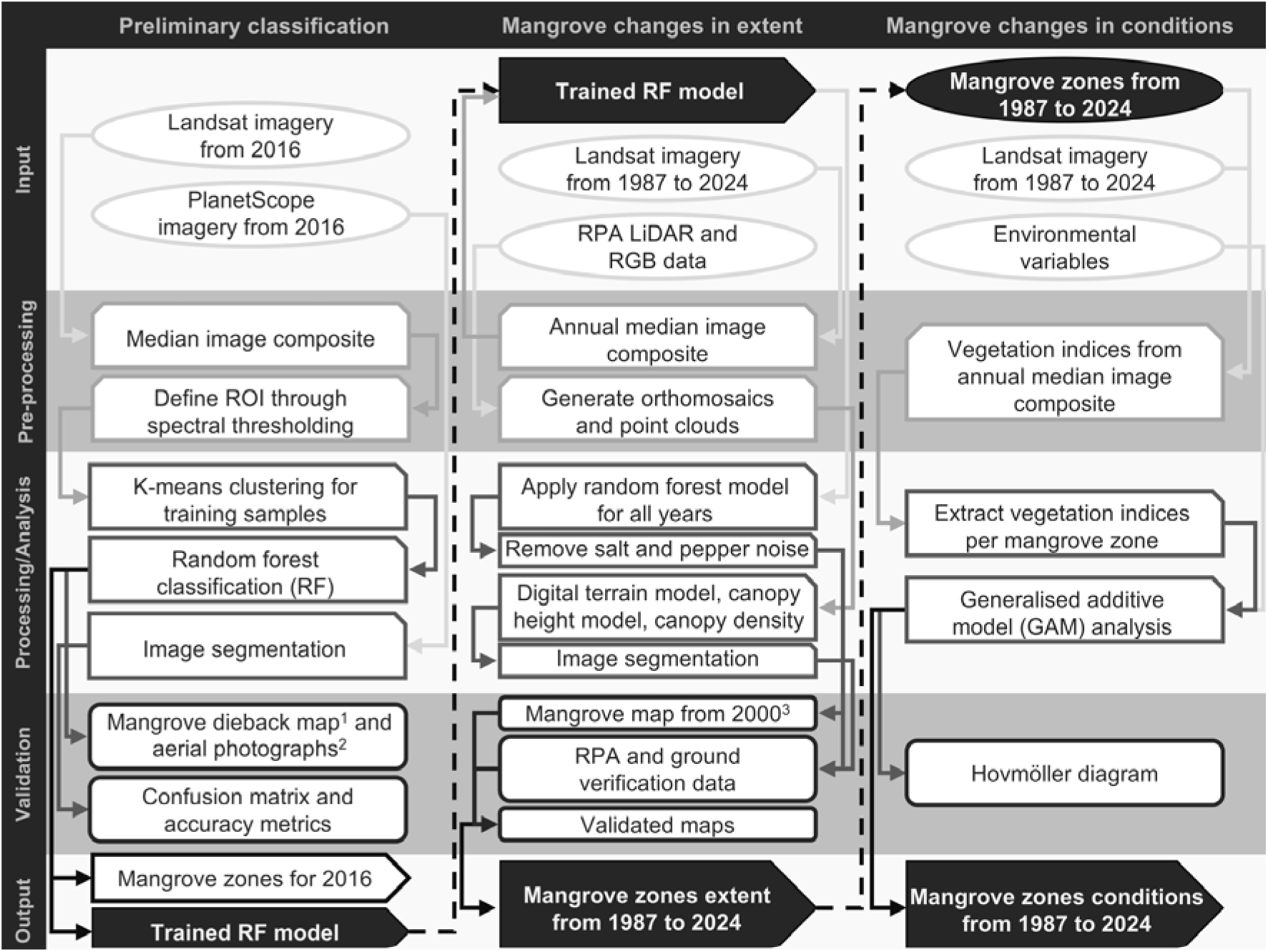
Workflow to map changes in mangrove extent and conditions. Continuous arrows show flow direction and dashed arrows show output being applied as input. ^1^Dieback map from 2016 (Duke et al. 2017), ^2^aerial photographs from 2017 (Duke et al. 2020), and ^3^mangrove classification map from 2000 (Hay 2009).

### 2.3. Mangrove changes in extent

#### 2.3.1. Preparation for classifying mangrove zones

Annual median composites were created for all available, analysis-ready, cloud-free Landsat 5 TM, 7 ETM+, and 8 OLI imagery (path 99, row 72, n = 1167) and used to classify mangrove zones yearly from 1987 to 2023. A mangrove mask was developed for each year by excluding ocean and inland water bodies using the Normalised Difference Water Index, and the Modified Normalised Difference Water Index. Non-mangrove vegetation was excluded using the Modular Mangrove Recognition Index (Diniz et al. 2019), which is a vegetation index used for identifying mangrove vegetation, and dieback areas were included based on the difference between 2016 and 2015 Modular Mangrove Recognition Index. These masks were extended to within 12 km of the coastline considering the Digital Earth Australia dynamic coastline dataset (Bishop-Taylor et al. 2021). Thresholds were defined based on visual interpretation of annual median composite from 2016 PlanetScope satellite data (Planet Labs PBC 2024), captured using the PS2 sensor with a spatial resolution of approximately 3.7 meters, and RPA-derived data (see *2.3.2. Ground verification and previous classification data*). Thresholds for mask development are presented in SI 1.

#### 2.3.2. Ground verification and previous classification data

RPA-derived data were collected at nine locations across the Albert and Leichhardt Rivers using DJI Phantom 4 RTK and DJI Matrice 300 RTK (Figure 1, SI 2). Flights using the DJI Matrice 300 RTK collected 3D point cloud data using Light Detection and Ranging sensor (LiDAR), and optical (red, green, blue; RGB) imagery data limited to areas that could be accessed on shore due to the size and weight of the RPA. For sites not accessible onshore, the Phantom 4 RTK RPA (smaller at <2kg) was deployed from a boat, collecting only RGB data. LiDAR point clouds from Matrice 300 RTK data were generated using DJI Terra for seven onshore accessible areas (Figure 1). Digital Terrain Models, Canopy Height Models and derived slope and aspect were generated for each site using an R script adapted from Australia’s Terrestrial Ecosystem Research Network (Sivanandam et al. 2023). Orthomosaics from both RPA were created for all areas (n = 9) using a Python-based script for Agisoft Metashape v 2.1.

To identify dominant species zones in RPA-derived data, object-based image analysis was undertaken through multiresolution segmentation in eCognition applied to the combination of digital terrain and canopy height models for the LiDAR-derived data and the orthomosaics for the DJI Phantom 4-derived data. The segmentation of each flight was initially performed using the default scale parameters of 100, shape of 0.1, and compactness of 0.5. To improve the segmentation, the scale parameter was decreased in increments of 10 until the object boundaries visually corresponded to the target mangrove dominant species zones. The RPA mangrove mask was generated to include all mangrove zones and was used to determine the thresholds for the Landsat mangrove mask.

RPA-derived data were further used for verification, with this supplemented with ground verification points indicating the presence or absence of mangroves and identifying the dominant mangrove species. These observations were undertaken by boat, with photographs takes as reference for their settings and structure (Figure 3), and geographical coordinates of each record using Google Maps.

**Figure 3.**
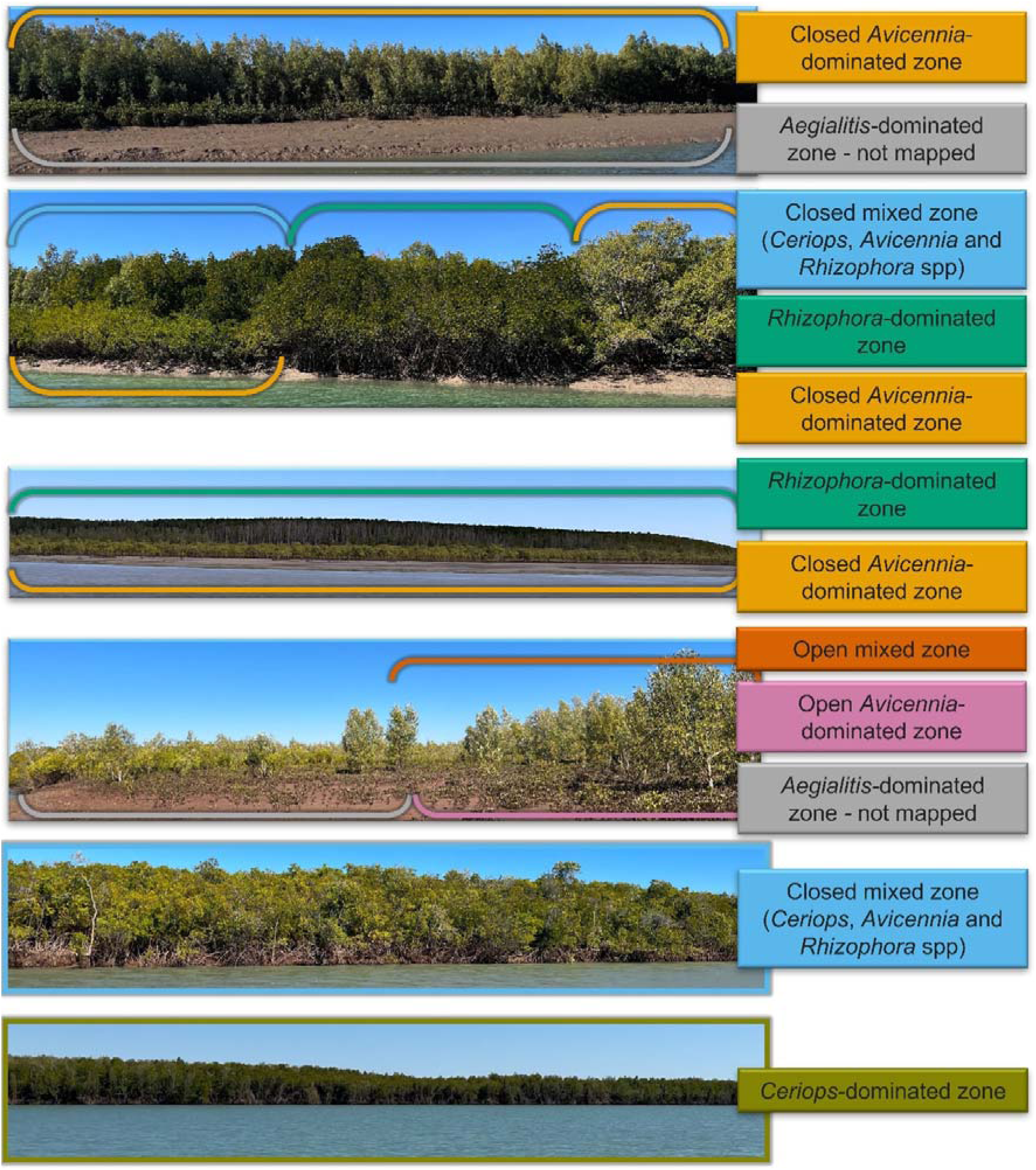
Main vegetation types and settings identified during field inspection. Photos are labeled according to how each class was characterised in the field.

Confusion matrices with overall, user and producer accuracies (Congalton 1991) were generated between the mapping and two products generated previously for the same areas. The first was produced by Hay et al. (2009), who mapped *Rhizophora*-dominated zone, closed *Avicennia*-dominated zone, closed mixed zone, closed *Ceriops*-dominated zone, and open *Avicennia*-dominated zone using Landsat imagery from 2000, providing a comparable mapping product for comparison with mangrove zone maps generated in this study. The second was Duke et al. (2017) which mapped mangrove dieback in 2016.

#### 2.3.3. Preliminary classification of mangrove zones

All available, analysis-ready, cloud-free Landsat 8 OLI imagery from 2016 (path 99, row 72, n = 24) were used to create an annual median composite and develop the mangrove mask for a baseline year. This baseline year was considered to represent the lowest extent and highest variability in condition over the Landsat record and provide the capacity to calculate changes relative to this year. A preliminary classification was performed on the 2016 masked annual composite using k-means to cluster the natural breaks of the data. The NDVI from 2015 and changes in NDVI between 2016 and 2015 were also included in the features for the preliminary classification to capture the historical context of previous mangrove extents. The number of classes was based on the number of dominant zones, including *Rhizophora-* dominated, *Avicennia*-dominated, *Ceriops*-dominated, open mixed, closed mixed, open forest dieback, and closed forest dieback. Visual assessment of the accuracy of preliminary mangrove classification was conducted using RGB and NDVI 2016 PlanetScope imagery and random sampling of 30 points within each mangrove zone. The accuracy of the classifications was again assessed through confusion matrices (SI 3). Unlike previous time series maps of mangrove extent (Lymburner et al. 2020), annual mangrove masks were prepared to ensure extent dynamics were effectively characterised for each mangrove zone.

#### 2.3.4. Time series classification of mangrove dominant species zones

Classification of dominant species within the mapped extent of mangroves for each year from 1987 to 2023 was conducted using Random Forest. This algorithm is widely used in remote sensing applications for classifying land cover because of its ability to handle multi-dimensional features with high accuracy and low computing demand. The main parameters defining the structure of the random forest in this model include the number of decision trees, the number of features considered for splitting at each node, the maximum depth of the trees, the minimum number of samples required for a split, and the minimum number of samples required at each leaf node. Optimal parameters for the model were estimated using the *gridsearchcv* function from the *sklearn.model_selection* library (Van et al. 2024). A total of 250 pixels for each class were randomly sampled from the 2016 preliminary classification, and 70% of these pixels were used for training the Random Forest classifier. The remaining 30% were used to evaluate model performance through a confusion matrix. ChatGPT 4o was used for optimising and debugging the classification models.

#### 2.3.5. Accuracy assessment

To validate the mangrove zones classification, 30 m radius circles were created at the centre of each ground verification point and the accuracy was assessed using confusion matrices of these circles for the 2023 classification (SI 8). RPA-derived data, including orthomosaics and the canopy height model, were used to visually compare the classified 2023 mangrove zones with 30 random points selected for each of the nine RPA surveys (n = 270) considering the orthomosaics and the canopy height model. Shoreline oblique photography from helicopter in 2017 and 2019 (Duke et al. 2020) was also used for visual verification for these years.

### 2.4. Mangrove changes in condition

#### 2.4.1. Data preparation

All available, analysis-ready, cloud-free Landsat 5 TM, 7 ETM+, and 8 OLI imagery (path 99, row 72, n = 1167) were used for evaluating the condition of mangrove zones from 1987 to 2023. NDVI was calculated for each image to serve as a proxy for vegetation health (Mafi-Gholami et al. 2019). An annual median composite was generated from these NDVI values to derive a consistent time series of mangrove condition.

Monthly data of the ENSO, IOD, and lunar nodal cycle were accessed from the Niño 3.4 Index (NINO3.4). (NOAA/NCEP 2024a), Dipole Mode Index – DMI (NOAA/NCEP 2024b), and mean sea level (MSL) from the Karumba tidal gauge (BOM 2024a), respectively.

#### 2.4.2. Evaluation of changes in mangrove conditions

Mangrove condition in this study is defined as the spatial and temporal variation in NDVI, reflecting changes in vegetation greenness as a proxy for health. Likewise, dieback refers primarily to a reduction in canopy greenness, indicated by lower NDVI values. Although dieback may sometimes indicate plant mortality, it can also represent partial canopy loss or reduced vegetation health, which may be reversible if environmental conditions improve. To identify stable or dynamic conditions, the number of times each pixel changed between mangrove classes over the study period was calculated. Additionally, visual interpretation of the relative proportion of changes was used in combination with array plots, Hovmöller diagrams, created for 53 transects, each 1 km in length, along the shoreline.

#### 2.4.3. Environmental influences on mangrove vegetation

To identify the temporal trends in mangrove conditions and extent related to climate variables, a series of Generalised Additive Models were fitted. A separate model was developed for each mangrove zone with the mangrove condition and extent metrics modelled as response variables, including the NDVI, estimated area, and their respective rates of change, with this calculated from the differences between two consecutive years. The models incorporated temporal effects and smooth interactions between the climate variables, which included the annual median of DMI, NINO3.4, and MSL, and their pairwise interaction. Model summaries are presented in effective degrees of freedom, adjusted R² values (proportion of variance explained for Gaussian models), deviance explained (likelihood-based measure of discrepancy), and p-values (SI 6 and 7). A Principal Component Analysis was also used to investigate the combined influence of climate variables on mangrove condition and extent for each classified zone. All statistical analyses were conducted in the R environment using *mgcv* (version 1.9-1) for Generalised Additive Models.

## 3. Results

### 3.1. Mangrove changes in extent

#### 3.1.1. Classification model evaluation

Accuracy metrics applied to the 2016 preliminary classification based on k-means clustering of the training samples achieved an overall accuracy of 68%. The training data performed best in the closed *Avicennia*-dominated zone (86%), *Rhizophora*-dominated zone (75%) and in both open (90%) and closed dieback (81%) zones, while the *Ceriops*-dominated (37%) and the open (45%) and closed (48%) mixed zones showed the lowest accuracies (SI 3). The subsequent Random Forest model (trained using these samples and applied to the 1987 – 2023 time series), was well-balanced and presented high performance across all classes with an overall accuracy of 97% (F1-score = 0.97, AUC = 0.99, SI 4). Additionally, the 2023 classification data, validated using the ground verification points collected from the boat, was 71% accurate overall, with the highest accuracies in the *Ceriops*-(88%) and *Avicennia*-dominated (85%) zones. These results provided confidence in applying the classification to time series data within Digital Earth Australia while also considering some limitations in the model for the mixed zones.

#### 3.1.2. Changes in mangrove extent

All mangrove zones increased in extent from 1987 to 2023, with notable gains seaward and along creeks inland, while losses were concentrated in landward areas (Figure 4A, H, Table 1). Most of the mangrove gain occurred where *Avicennia*-dominated zones prograded seaward (Figure 4C). Inland creeks experienced substantial landward extension, primarily by *Ceriops* sp. and mixed zones (Figure 4A-C). Mangrove loss occurred mostly in landward regions (Figure 4A), especially in areas previously dominated by *Rhizophora stylosa* or *Ceriops* sp. (Figure 4C, D). The most stable zones were in the centre of the mangrove forest (Figure 4A), which remained classified primarily as *Rhizophora*-dominated zone throughout the entire study period (Figure 4A, D).

**Figure 4.**
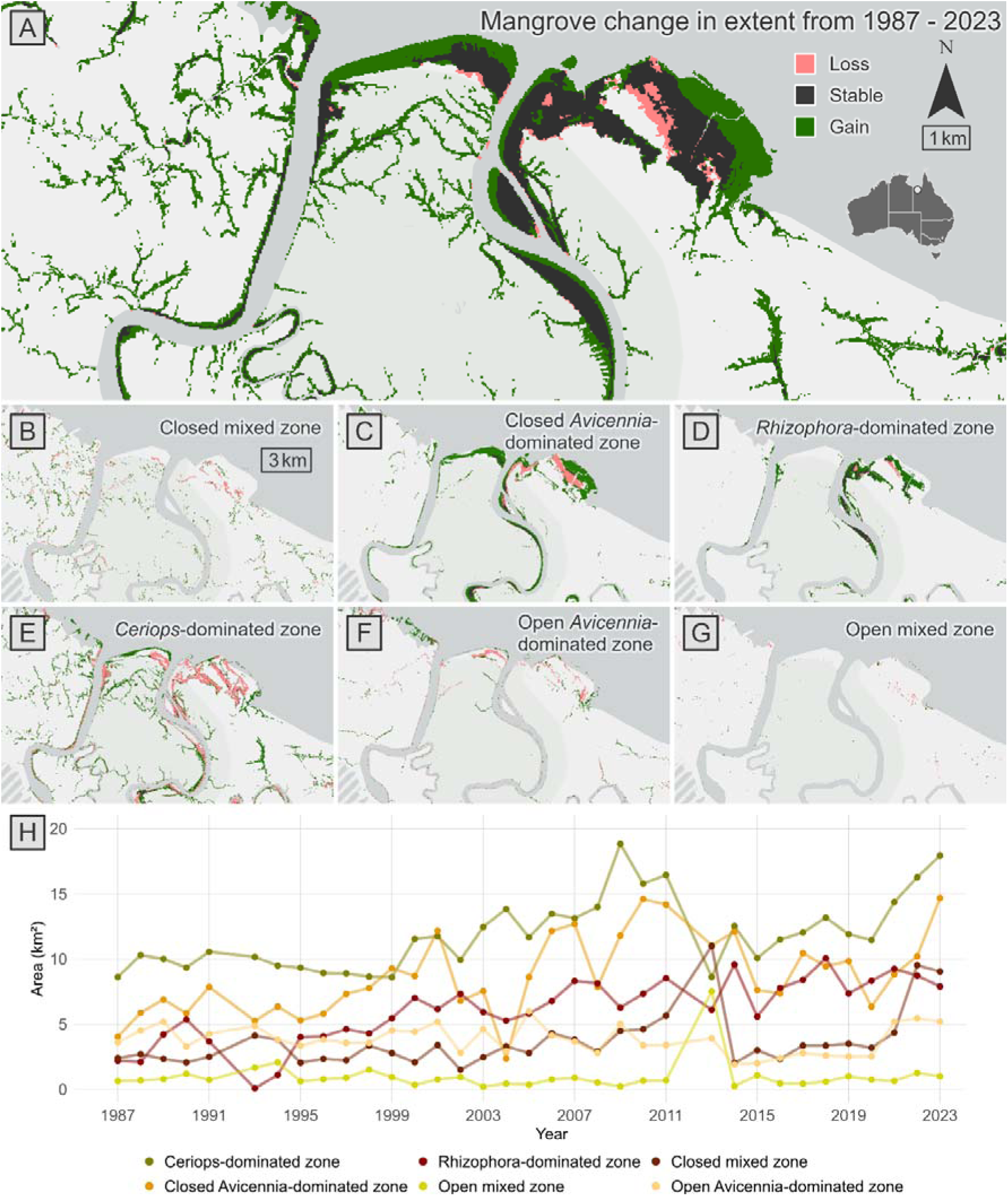
Mangrove loss (pink), gain (green) and stable areas (dark grey) between 1987 and 2023 for (A) all mangrove zones, (B) closed mixed zone (3km bar shows the scale for plots B – G), (C) closed *Avicennia*-dominated zone, (D) *Rhizophora*-dominated zone, (E) *Ceriops*-dominated zone, (F) open *Avicennia*-dominated zone and (G) open mixed zone. Change in the extent of (H) area of mangrove zones.

**Table 1.**
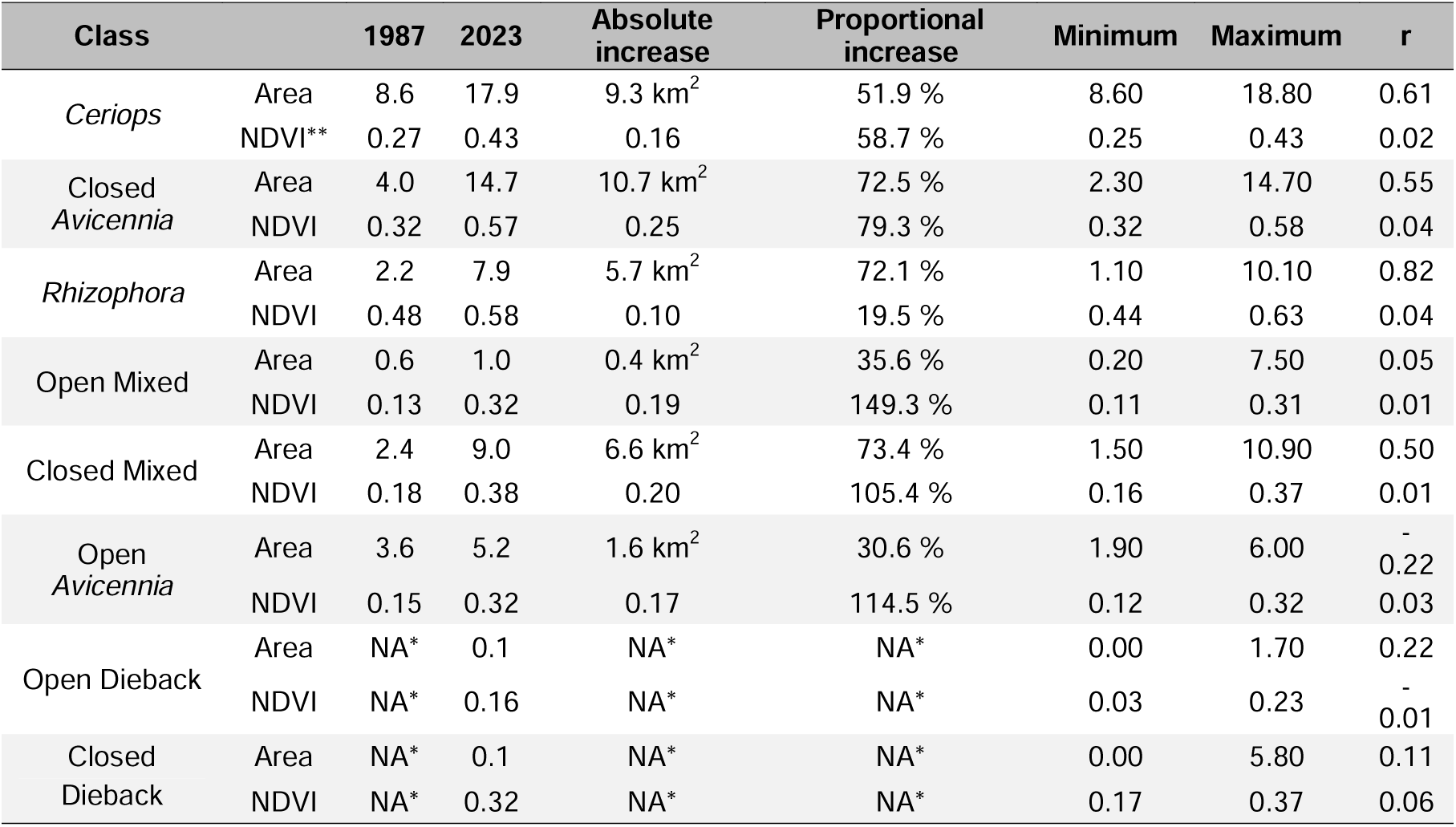
Proportional and absolute increase in extent (area, km^2^) and NDVI of each mangrove zone from 1987 to 2023, minimum and maximum extent and condition and slope of the fitted area and NDVI trends over the years. R is used as an indicator of the strength of the linear trend of each variable (NDVI and area) over time. *No values for calculating changes since the first classification map due to the absence of these classes. ** Mean values.

Closed *Avicennia*-dominated zone showed the greatest total area increase seaward from 4.0 km^2^ in 1987 to 14.7 km^2^ in 2023 (Figure 4C, H, Table 3). Expansion was also evident along tidal creek networks, where *Ceriops*-dominated zones occupied these areas (Figure 4E, Table 1). Meanwhile, *Rhizophora*-dominated zones comprised two categories: (1) mature *Rhizophora*-dominated areas where high NDVI remained relatively stable through the time series, hereafter the stable forest; and (2) areas prograding seaward, resulting from the conversion of other mangrove classes into *Rhizophora*-dominated zones (Figure 4D, H). By contrast, retreat of mangroves on the landward margin was observed at the higher elevation areas (Figure 5). Periods of dieback were recorded across several intervals, with the most pronounced events occurring between 2002-2004 (9.1 km^2^), 2008 (2.9 km^2^), and 2015-2016 (8.1 km^2^) (Figure 4I). These events were more pronounced in closed compared to open forest areas (Figure 4I).

**Figure 5.**
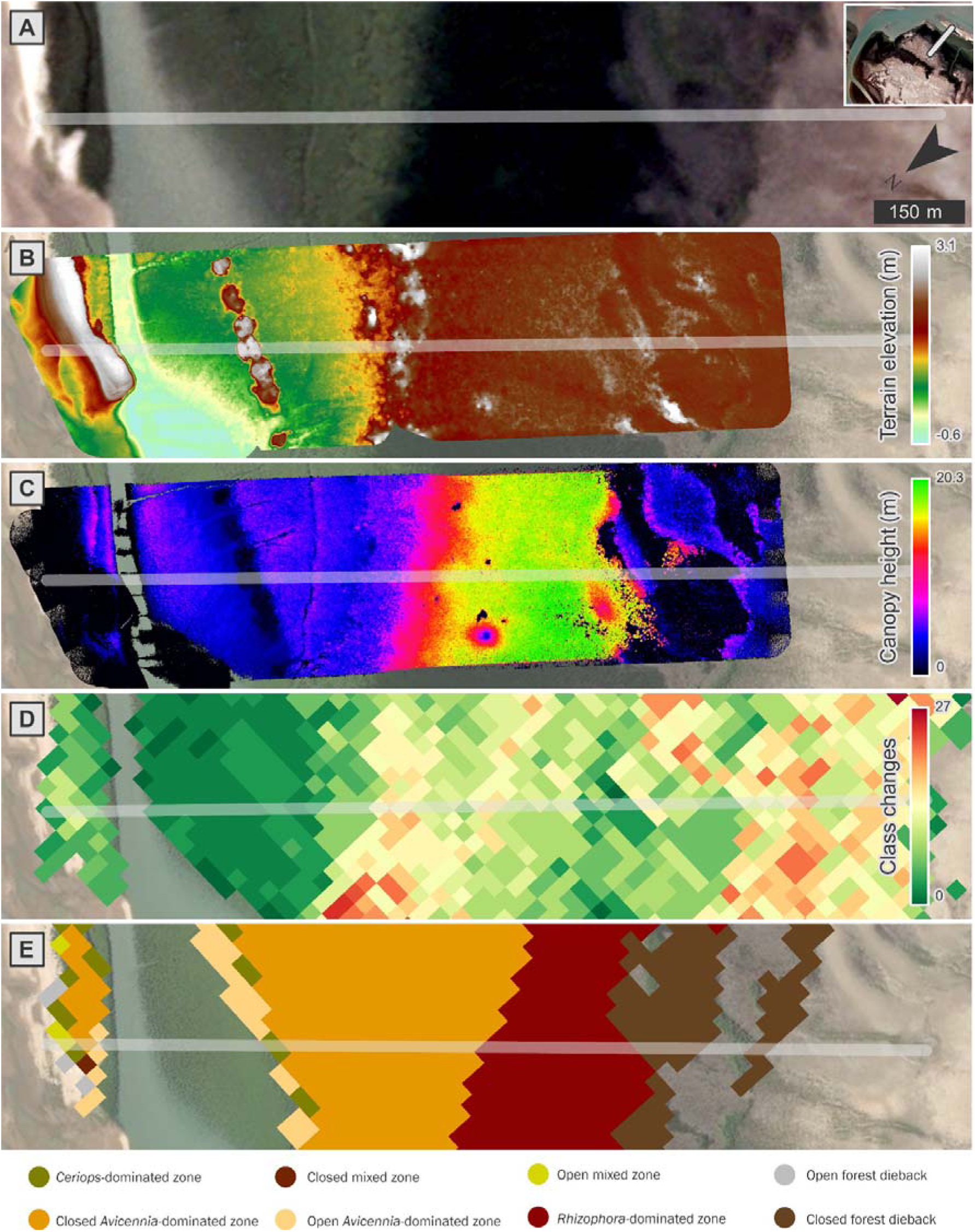
High resolution satellite imagery from 2017 (Queensland Globe), post dieback (A), drone-derived digital terrain elevation (B) and canopy cover (C) from 2024, class changes over the 1987 – 2023 period (D), and classified mangrove zones (E) for transect 57 (transparent line).

#### 3.1.3. Changes in mangrove condition

The NDVI progressively increased across all classes, indicating the mangroves were increasing in extent and becoming greener (Table 1). The greatest NDVI increases occurred in the newly established seaward fringe areas, where closed *Avicennia*-dominated zone prograded (Figure 6, Table 1). By contrast, the stable forests, dominated by mature *Rhizophora stylosa*, had the least change in condition (+0.09), reflecting stable and healthy vegetation conditions throughout the study period. This class also showed the highest NDVI among all zones at 0.63 (2018). However, not all locations experienced NDVI gains. Transects 26 and 57, for example, showed periods of declining NDVI due to the 2002/2004 and 2015/2016 dieback events, with these coinciding with low sea-level periods (Figures 4H and 6B–G).

**Figure 6.**
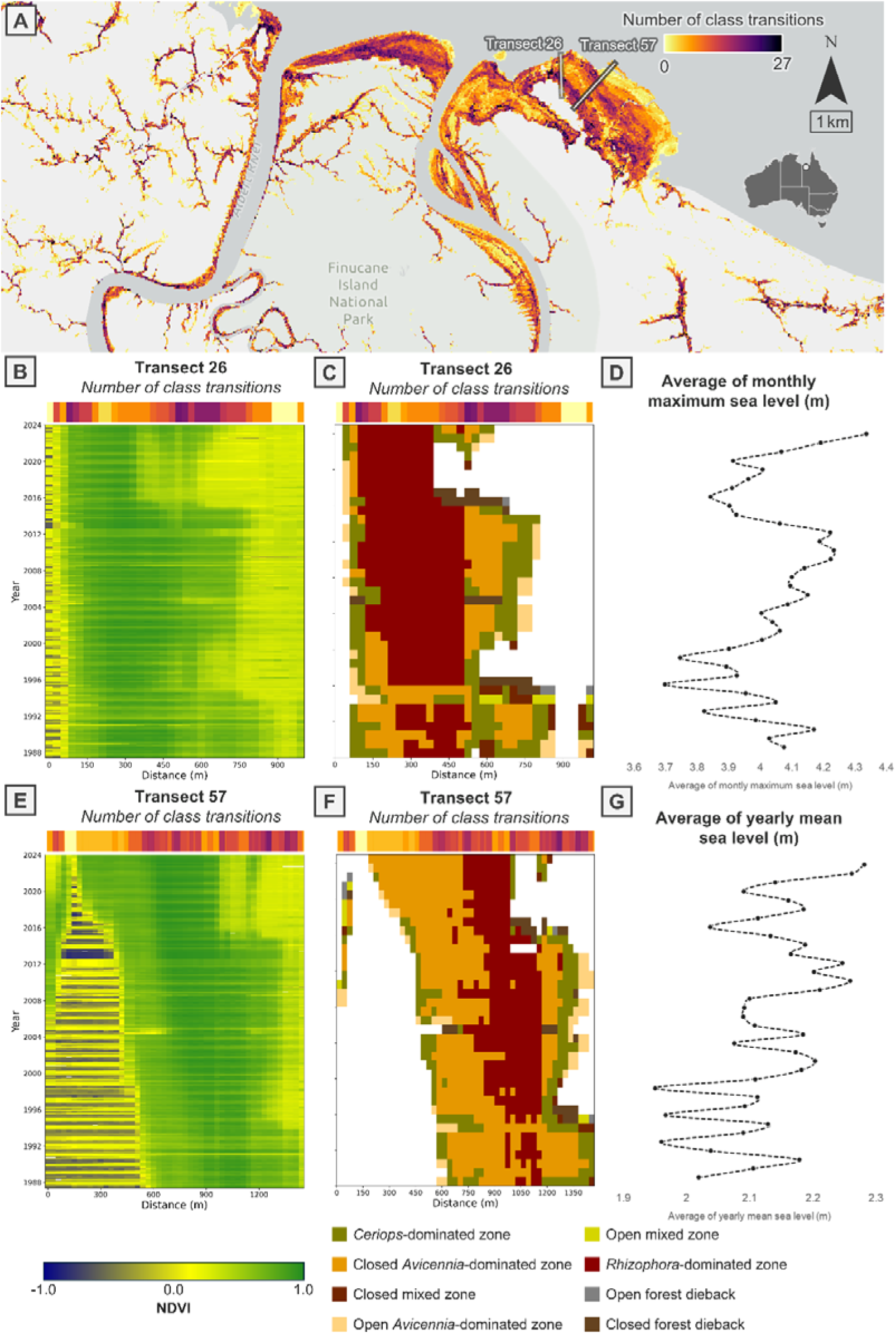
Cumulative number of class transitions, yellow showing no changes and purple representing the highest changes (26) identified throughout the study period and transects 26 and 57 as reference (A), array plot of NDVI (B, E) and classification zones (C, F) over time on transect 26 and 57, where white areas show no mangrove zones, and variation in the lunar nodal cycle expressed in sea level in the closest tidal gauge (Karumba) with average monthly maximum sea level (D) and yearly mean sea level (G). Array plot in B, C, E and F show changes in mangrove conditions over space and time for NDVI and mangrove zones. The x-axis represents the length of the transect, showing spatial variation, while the y-axis represents time, showing temporal changes. Colours indicate the values of NDVI or dominant species, illustrating how mangrove conditions vary along the transect and change over time at these specific locations.

Regions with the least class transitions were observed towards the seaward margin and established forests, particularly in *Rhizophora*-dominated zones (Figures 6A, 5C–E). The seaward margin, where *Avicennia marina* colonised areas previously occupied by water for an extended period presented minimal changes over time. Progradation became more pronounced after 2012, see Transect 57 (Figure 6E). Similarly, the *Rhizophora*-dominated zone, remained relatively stable in extent throughout the entire study period, and retained high NDVI values with minimal fluctuations (Figure 6A, B, D). By contrast, the landward margin presented the highest number of class transitions (Figure 6A, C, E) and greater NDVI variability (Figure 6B, D). These were often aligned with environmental fluctuations, such as changes in mean sea level (Figure 6D). These shifts in the landward zones occasionally led to dieback events, which were also associated with lower tidal heights (Figure 6B-E) and higher elevations in the tidal frame, see Transect 57 (Figure 5B, D, E). However, mangrove dieback was usually followed by recovery after 3 to 4 years, with *Avicennia-* or *Ceriops*-dominated zones being the first species to re-establish post-dieback areas (Figure 6C).

### 3.2. Environmental variables related to changes in mangrove vegetation

Overall, the Generalised Additive Models for temporal trends in mangrove conditions showed that ENSO, IOD and MSL are frequently associated with changes in NDVI and area across multiple zones (Sup Tables 6 and 7). Changes in NDVI and area for the closed *Avicennia*-dominated zone were not explained by ENSO, IOD, or MSL, even though the model for NDVI explained 85.2% of the deviance (R^2^ = 0.74). This suggests that there are other factors likely driving these changes. However, PCA (PC1 = 47.3% variance) loadings were positive for NDVI (+0.48) and MSL (+0.54) and negative for ENSO (-0.47), indicating that while higher MSL typically correlates with better canopy condition, strong El Niño events could lead to a decrease in NDVI (Figure 7A). On the other hand, changes in area of *Rhizophora*-dominated zones were significantly influenced by interactions between IOD and MSL (p < 0.01) and ENSO and MSL (p = 0.03). Specifically, higher IOD or ENSO values combined with lower MSL were associated with a reduced extent for this zone, with the model explaining 87.6% of the deviance (R^2^ = 0.78). The PCA (PC1 = 49.7% variance) supports this, showing positive loadings on area (+0.48) and MSL (+0.54) but negative loadings on ENSO (-0.49) and IOD (-0.15) (Figure 7B). A trend was not observed for NDVI in the model for *Rhizophora*-dominated zone. In the closed forest dieback areas, changes in NDVI were associated with ENSO and MSL, with lower NDVI occurring under higher ENSO and lower MSL (p < 0.01) (Figure 7 D – F). The model explained 90.5% of the deviance (R^2^ = 0.76). The PCA corroborates these results with PC1 accounting for 49.6% of the total variance, showing negative loadings on NDVI (-0.47) and MSL (-0.56), but positive loading on ENSO (+0.59) (Figure 7C). This zone exhibited the strongest relationships with environmental variables (Figure 7 G – I), with time (p = 0.01), DMI (p < 0.01), and NINO3.4 (p < 0.01) significantly related to area, with the model explaining 57% of the deviance (R-sq adj = 0.43).

**Figure 7.**
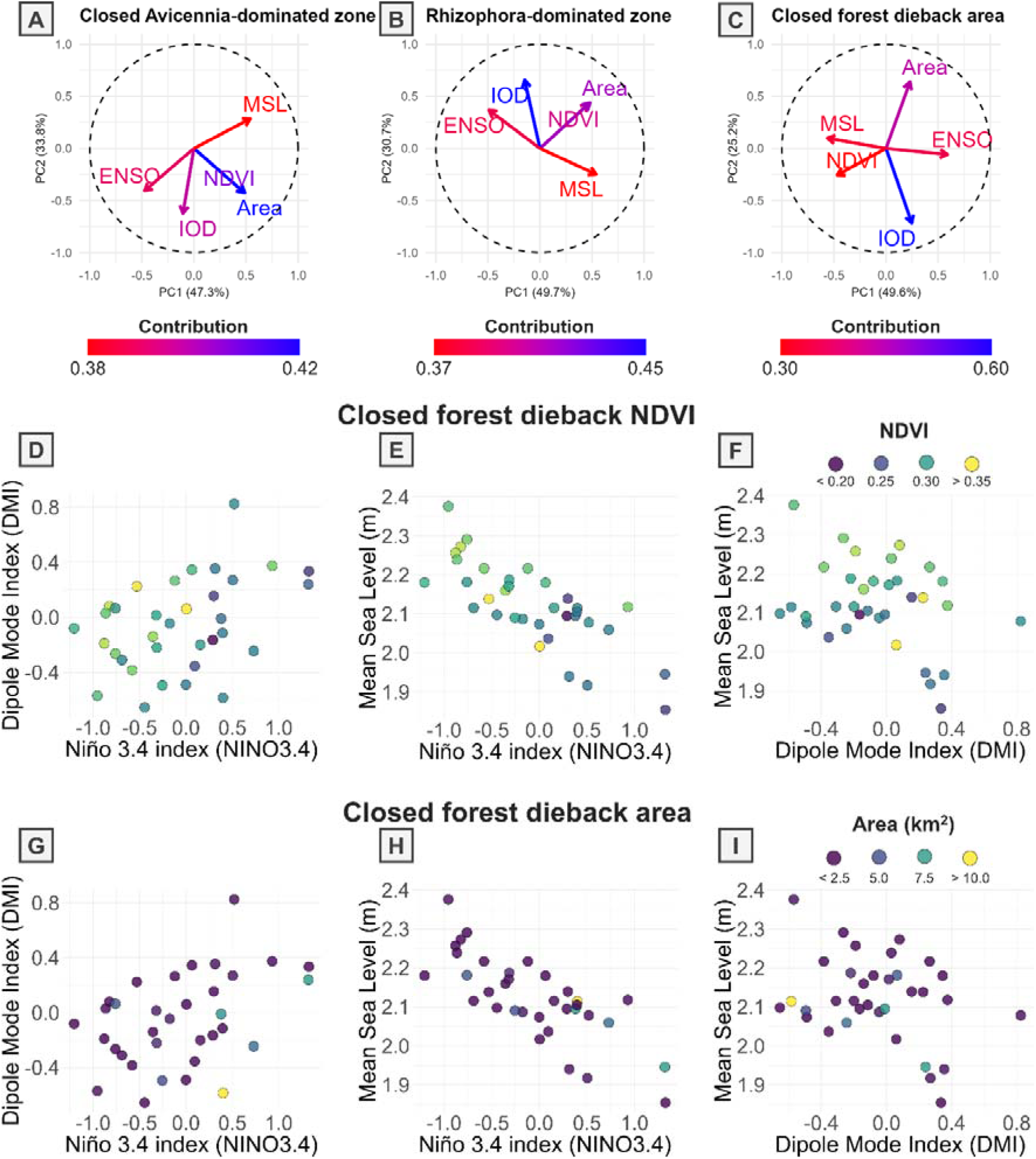
(A – C) PCA for closed *Avicennia*, *Rhizophora*, and closed dieback zones, with variable contributions shown in a red-blue gradient. (D – F) Observed IOD (Dipole Mode Index), ENSO (Niño 3.4 Index), and MSL for each annual observation (n = 36) in the closed forest dieback zone coloured by NDVI on a blue-yellow gradient (blue = lower NDVI, yellow = higher NDVI). (G – I) The same annual IOD, ENSO, and MSL distributions for the closed forest dieback zone, but coloured by area (blue = lower area, yellow = higher area). Panels D – I illustrate raw data points averaged for each year.

### 3.3. Model performance and limitations

The classification model used in this study, combined with ground verification points, is robust, with high performance on the Random Forest classifier (overall accuracy = 97%; F1-score = 0.97, AUC = 0.99). These results indicate a reliable capacity to differentiate mangrove zones, especially in closed *Avicennia*-, *Rhizophora*-dominated, and dieback zones. However, the preliminary classification using k-means clustering showed lower accuracy (68%), highlighting challenges in identifying the mixed and *Ceriops*-dominated zones where overlaps in spectral signatures or transitional areas may contribute to misclassification. The 2023 ground verification points improved the model performance (71% accuracy), improving the model’s limitations in distinguishing the challenging zones.

The field verification data (SI 8) showed that terrestrial vegetation, *Rhizophora*-, and *Ceriops*-dominated zones had the lowest user accuracies (36%, 50% and 58%, respectively, SI 8). Most of the points classified as *Rhizophora*-dominated were actually closed mixed zones in the field verification data, while most *Ceriops*-dominated areas in the model were either closed mixed or *Rhizophora*-dominated zones in the field data. Several closed mixed zones were classified as not mangrove in the 2023 map, indicating that the model has limitations in identifying this class as mangrove areas. This might occur due to the complexity of different species in the same pixel or the size of the areas, limited by the spatial resolution of the satellite. *Avicennia*-dominated and closed mixed zones had the highest user accuracies (85% and 90%, respectively).

When compared to the previous classification (Hay 2009, SI 9) and dieback (Duke et al. 2017, SI 10) maps, overall accuracy was 43% and 76%, respectively, suggesting only a small agreement between previous and current classifications. The closed *Avicennia*-dominated zone had the highest user accuracy (49%, SI 9). In contrast, the *Rhizophora*-dominated zone had a low user accuracy (47%, SI 8). The closed mixed zone was the class with the lowest accuracy (16%, SI 9), followed by the *Ceriops*-dominated zone (0.449, SI 9), and open *Avicennia*-dominated zone (0.47, SI 9). *Ceriops*-dominated observations on the previous classification (Hay 2009) were mostly classified now as closed *Avicennia*-or *Rhizophora*-dominated zones. The producer accuracy for no dieback was 0.68 (SI 10), indicating that approximately 32% of no dieback areas identified previously were predicted as dieback now. The user accuracy comparing the previous dieback classification map with the current was 0.53 (SI 10), suggesting that nearly half of the areas predicted as dieback do not match the previous classification. The use of high-resolution RPA data, combined with advanced classification algorithms and field verification data in this study improved the previous classification maps for the mangroves from Albert and Leichhardt Rivers.

While variations in the NDVI were well-explained across most zones and indicated that mangrove health might be linked to climatic variability, models for changes in the area showed poorer fits overall. For example, the change in area model for the *Ceriops*-dominated zone had a low adjusted R-squared value (R-sq adj = 0.04), suggesting that unmeasured factors, such as sediment deposition, might drive changes in mangrove extent for this zone or this zone might have a longer lag time to respond to environmental changes. In contrast, the open forest dieback zone demonstrated a good fit for NDVI (R-sq adj = 0.961, 98% deviance explained) with time as the only significant predictor (p < 0.001). This indicated a consistent temporal trend in vegetation health despite minimal influence from ENSO or IOD.

## 4. Discussion

### 4.1. Mangrove zones are greening and expanding despite dieback

Despite the two episodic dieback events shown in the region in 2002-2004 and 2015-2016, net increases in area and greenness of all mangrove zones over 36 years were identified. Closed *Avicennia*-dominated zones presented the highest proportional increase in extent and greenness, showing consistent expansion over the years, particularly in the seaward direction and along some tidal creeks. This highlights their capacity to colonise new substrates, with this attributed to their tolerance to varying salinity and inundation levels, particularly where sediment deposition is high, and accommodation space is available (Woodroffe et al. 2016). By contrast, the open mixed and open *Avicennia*-dominated zones showed the least expansion, with most remaining stable between 1987 and 2023. This may reflect the high dynamism of these areas in extent and condition, where both open zones experienced significant increases in condition, even converting to closed forest in some areas, while also transitioning from non-mangrove vegetation to open mangrove areas. This relatively rapid change in zone classification aligns with the fast-growing characteristics of the *Avicennia marina* and the capacity of this species to establish rapidly on suitable substrates where sedimentation is high (Almahasheer et al. 2016). These patterns of increased condition in all mangrove zones over time corroborate findings from previous studies documenting mangrove greening in northern Australia (Lymburner et al. 2020) and during post-dieback recovery periods (Lovelock et al. 2017; Hickey et al. 2021). Significant increases in mangrove greenness are usually attributed to favourable climatic conditions, including rainfall and sediment availability (Asbridge et al. 2016).

### 4.2. Mangrove zones are dynamic and characterised by dieback, recovery, and progradation

Mangrove forests in the Albert and Leichhardt Rivers are highly dynamic, fluctuating in extent and condition over time. Two dieback episodes were identified in the area, each followed by 3 to 6 years of canopy recovery. The most severe dieback events, in 2002/2003 and 2015/2016, coincided with extreme climate conditions and large-scale mangrove mortality in other parts of northern Australia (Lovelock et al. 2017; Hickey et al. 2021).

However, the recovery time was shorter, estimated at 4 to 7 years, compared to the ∼10 years observed elsewhere, suggesting greater resilience in the area. While episodic dieback events are often considered detrimental to mangrove persistence (e.g., Duke et al. 2017), evidence from this study suggests these events are part of a natural cycle of loss and recovery. Over the past three decades, this cyclic process has contributed to an overall increase in mangrove extent and condition across all mangrove zones.

Cyclic patterns of declining condition and recovery are reflected differently across the study area, with dieback occurring in the landward zone and progradation in the seaward zone. Landward zones, dominated by *Avicennia marina* and *Ceriops* sp., experience cycles of extension and contraction at the landward fringe. These fluctuations likely contributed to frequent transitions between vegetation classes, with NDVI trends predominantly indicating increases. Similarly, *Rhizophora*-dominated landward zones also present variability in extent and condition, but their sensitivity to declining NDVI is likely related to their occurrence at elevations in the tidal frame where inundation is less frequent, making them particularly vulnerable to reduced inundation (Harris et al. 2018; Asbridge et al. 2019; Saintilan et al. 2022). In contrast, seaward zones, dominated by *Avicennia marina*, present a general trend of progradation, likely driven by sediment accretion, as river-derived sediments are deposited in these areas, such as the first 300 m of transect 57, where seaward expansion aligns with areas expected to experience high sediment deposition from the Leichhardt River. Over the past 36 years, NDVI trends in seaward zones indicate an overall increase, suggesting enhanced productivity and lateral expansion of mangroves. These patterns align with broader trends in the Gulf, where widespread mangrove progradation has been associated with sediment deposition, propagule establishment, and periods of high rainfall and river discharge (Asbridge et al. 2016; Rogers et al. 2024). The ability of mangroves to prograde, despite rising sea levels, reflects their capacity to adapt to changes in environmental conditions.

Sediment availability is a critical factor in mangrove expansion, and in regions where tidal deposition and riverine sediment influx are sufficient, mangroves continue to prograde (Woodroffe et al. 2016).

### 4.3. Despite dynamism, stable mangrove forests have persisted

While mangrove forests in the Albert and Leichhardt Rivers present dynamic shifts in response to environmental changes, stable forests dominated by *Rhizophora stylosa* little to no variation in condition over time. These stable zones are primarily located at the centre of the forest, where mature *R. stylosa* persist with minimal variation in NDVI. Their persistence suggests that these areas experience favourable hydrological conditions and tidal connectivity, which may contribute to their structural resilience against environmental variability (Rogers et al. 2024). Additionally, old-growth *Avicennia marina* forests have also contributed to this stability, particularly in the seaward transitional zones before *R. stylosa* expands and establishes dominance. The persistence of these mature forests at the Albert and Leichhardt Rivers indicates that while *A. marina* initially colonised certain areas, *R. stylosa*-dominated zones eventually replace *A. marina*-dominated zones as progradation advances seaward. This process highlights the longer-term resilience of these stable forests. Further, the resilience of these stable forests may provide improved confidence in mangrove biomass as a blue carbon store that contributes to climate mitigation. Similar stability has also been documented in other areas of the Gulf (Asbridge et al. 2016) and along the Australian coastline (Lymburner et al. 2020). These findings demonstrate the ecological importance of stable mangrove forests in carbon sequestration and habitat maintenance.

### 4.4. Combined effects of climate cycles drive mangrove changes in extent and condition

ENSO-driven fluctuations in sea level and water availability were identified as primary drivers of changes in mangrove extent and condition in the Albert and Leichhardt Rivers, with tidal inundation frequency and sediment supply likely being the dominant controlling factors. Periods characterised by high sea levels, associated with ENSO events and intensified by the lunar nodal cycle, resulted in seaward expansion and improved mangrove condition. The accretion of sediment, combined with higher sea levels during positive phases of ENSO, may have enabled the seaward expansion of mangroves, providing a substrate for seedlings to establish and thrive indicating the importance of sediment supply in sustaining mangrove ecosystems under changing climatic conditions. This pattern of progradation associated with sedimentation processes is also reported to drive seaward expansion and greening elsewhere (Alongi et al. 2012; Swales et al. 2015). Conversely, mangrove dieback coincided with reduced tidal inundation during prolonged low sea-level periods, primarily affecting landward zones at higher elevations where environmental stresses were compounded. This vulnerability was reflected in greater NDVI variability and more frequent class transitions in these zones. Landward mangrove retreat along with declines in NDVI observed during the 2015/2016 mangrove dieback in the Gulf were similarly documented in Kakadu National Park, Northern Territory (Asbridge et al. 2019) and in Mangrove Bay, Western Australia (Lovelock et al. 2017; Hickey et al. 2021). Dieback is reported to occur when tidal inundation is reduced due to prolonged low sea levels (e.g. Lovelock et al. (2017), Sippo et al. (2020)), exacerbated by extreme positive ENSO and the influence of the lunar nodal cycle (Saintilan et al. 2022). The effects of the 18.6-year lunar nodal cycle and the 4.4-year perigean subharmonic on tidal amplitude modulation in the Gulf is amongst the strongest in the world, causing variations between 0.3 and 0.6 m (Haigh et al. 2011). High tidal amplitude phases enhance inundation frequency and duration, promoting mangrove growth and resilience, while the low amplitude phases reduce tidal inundation, intensifies drought impacts and vulnerability to dieback (Saintilan et al. 2022). While the IOD has been linked to mangrove changes elsewhere (Asbridge et al. 2019; Carruthers et al. 2024), its influence in this region appears to be limited. The absence of extreme cyclones (BOM 2025) and intensive human activities (Lucas et al. 2019) in the study area over the past four decades further supports that mangrove changes are driven by the effects of these environmental variables.

Most dieback events occurred in the landward margins of the forest at higher elevations within the tidal frame, where the compounded environmental stresses were more pronounced. This was evidenced by greater NDVI variability and a higher frequency of class changes in these zones. These findings demonstrate that mangrove landward zones are particularly vulnerable to declines in condition and potential dieback when exposed to compound climatic effects. The unusually low sea level and extremely negative IOD and positive ENSO phases have been linked to mangrove changes in extent and condition globally (Ward et al. 2016; Ximenes et al. 2016; Carruthers et al. 2024). In the Maldives, the extremely negative IOD phase in 2020 caused seawater ponding in low-lying mangrove basins, which intensified salinity stress and tree mortality (Carruthers et al. 2024). During combined climate cycle events, such as the 2015/2016 dieback in northern Australia, the prolonged drops in sea level change tidal cycles, reducing inundation and, consequently, limiting water availability to mangrove roots (Lovelock et al. 2017; Saintilan et al. 2022). Alongside this, elevated temperatures and reduced rainfall during these periods tend to increase soil desiccation, creating hypersaline conditions that compromise mangrove water uptake (Lovelock et al. 2017). However, climate events alone are insufficient to fully explain the extensive spatial variation in mangrove dieback severity observed in the Gulf of Carpentaria during 2015/2016. The coincidence of an extreme positive ENSO phase with the minimum amplitude phase of the 18.61-year lunar nodal cycle intensified the drought-induced stress by reducing tidal inundation (Saintilan et al. 2022). Neighbouring regions that experienced the same extreme ENSO phase, such as the Arnhem coast, Northern Territory, but high tidal amplitude phase of the lunar nodal cycle, presented less mangrove dieback due to maintenance of tidal inundation, creating a buffer against drought stress (Saintilan et al. 2022).

Sea-level fluctuations, associated with ENSO and IOD, indicated a cyclical pattern of mangrove loss and recovery, with *Avicennia*-and *Ceriops*-dominated zones recovering and expanding faster than *Rhizophora*-dominated zones. Mangrove recovery and expansion typically occurred during positive phases of IOD and negative phases of ENSO. Similar cycles of dieback and recovery associated to climate cycles are reported globally (Maxwell et al. 2018; Masud-Ul-Alam et al. 2021). However, projections of stronger ENSO and IOD events, accelerated sea-level rise, and more frequent droughts suggest that if these factors exceed mangrove tolerance, dieback may become more common on mangrove, and landward retreat may accelerate due to insufficient sediment accretion (Saintilan et al. 2020, 2023).

## 5. Conclusion

Despite experiencing episodic dieback events, mangrove forests in the Albert and Leichhardt Rivers have recovered and increased in extent and condition over the past 36 years. These changes were strongly associated with ENSO-driven fluctuations in sea level, primarily modulated by the lunar nodal cycle. This suggests that while extreme events may temporarily reduce canopy cover, they do not necessarily lead to long-term ecosystem collapse. However, if the frequency or intensity of extreme climatic events increases, as future climate projections suggest, mangrove recovery may be compromised, particularly in vulnerable landward zones. Additionally, despite episodic sea-level fluctuations associated with climatic cycles, the continued progradation of mangroves in the seaward zone implies sediment supply is sufficient to support seaward expansion. At the same time, even in this highly dynamic mangrove system, the conditions of stable mangrove forests have persisted relatively unchanged for at least 36 years, serving as long-term blue carbon reservoirs. This stability supports aboveground carbon storage through continuous tree growth and belowground carbon accumulation in sediments. However, while seaward progradation and stable forest resilience are evident, landward zones remain highly vulnerable to environmental stressors. Landward retreat dynamics should be closely examined, particularly how hydrological constraints, salinity stress, and elevation influence mangrove zonation shifts. Given that landward zones are the most vulnerable to dieback during extreme climate events, understanding their capacity for recovery will provide critical insights into the long-term stability of these ecosystems. Additionally, decreases in NDVI often precede and can be used to anticipate dieback events. Therefore, understanding how vegetation indices might serve as early indicators of vegetation stress is crucial for predicting and acting promptly against increasing extreme climatic events.

## Authors contributions

RG and KR conceived the ideas, RG designed methodology; RG and KR collected the data; RG analysed the data; RG led the writing of the manuscript. All authors contributed critically to the drafts and gave final approval for publication.

## Data accessibility

Datasets for LiDAR and RGB flights for the area will be provided upon request to the corresponding author. Open-source code developed for this study is available in GitHub (https://github.com/rogervsg/supplementary_material_leichhardt.git).

## Acknowledgements

We thank Professor Colin Woodroffe for his valuable comments on the manuscript. We also thank Lurick Sowden and Maxine Sowden from the Gangalidda and Garawa Land and Sea Ranger Unit who provided valuable assistance during the field component of the study, the Gangalidda and Garawa People, Carpentaria Land Council Aboriginal Corporation and Yagurli Tours. We also thank Queensland Parks and Wildlife Services for providing the permit for data collection (P-PTUKI-100674037). This research was supported by the Australian Research Council Discovery Project (ARC DP21O1WO739), and UOW University Post Graduation Award.

---

SI 1. Vegetation indices used for defining the mangrove mask. SI 2. Flight details and RPA pre-processing information

SI 3. Confusion matrix for the 2016 preliminary classification data. SI 4. Classification report for test data and predictions.

SI 5. Confusion matrix for test data and predictions.

SI 6. Model summaries for Generalised Additive Models, including adjusted R² values, deviance explained, and p-values.

SI 7. Model summaries for Generalised Additive Models, including effective degrees of freedom.

SI 8. Confusion matrix for mangrove classification from 2023 using field verification data as a reference.

SI 9. Confusion matrix for mangrove zonation from 2000 using reference data (Hay 2009). SI 10. Confusion matrix for mangrove dieback from 2016 using Duke et al. (2017) as reference data.

SI 11. High resolution imagery from 1988 and 2017, Hovmöller for NDVI and mangrove classification zones for transects 1 to 57.

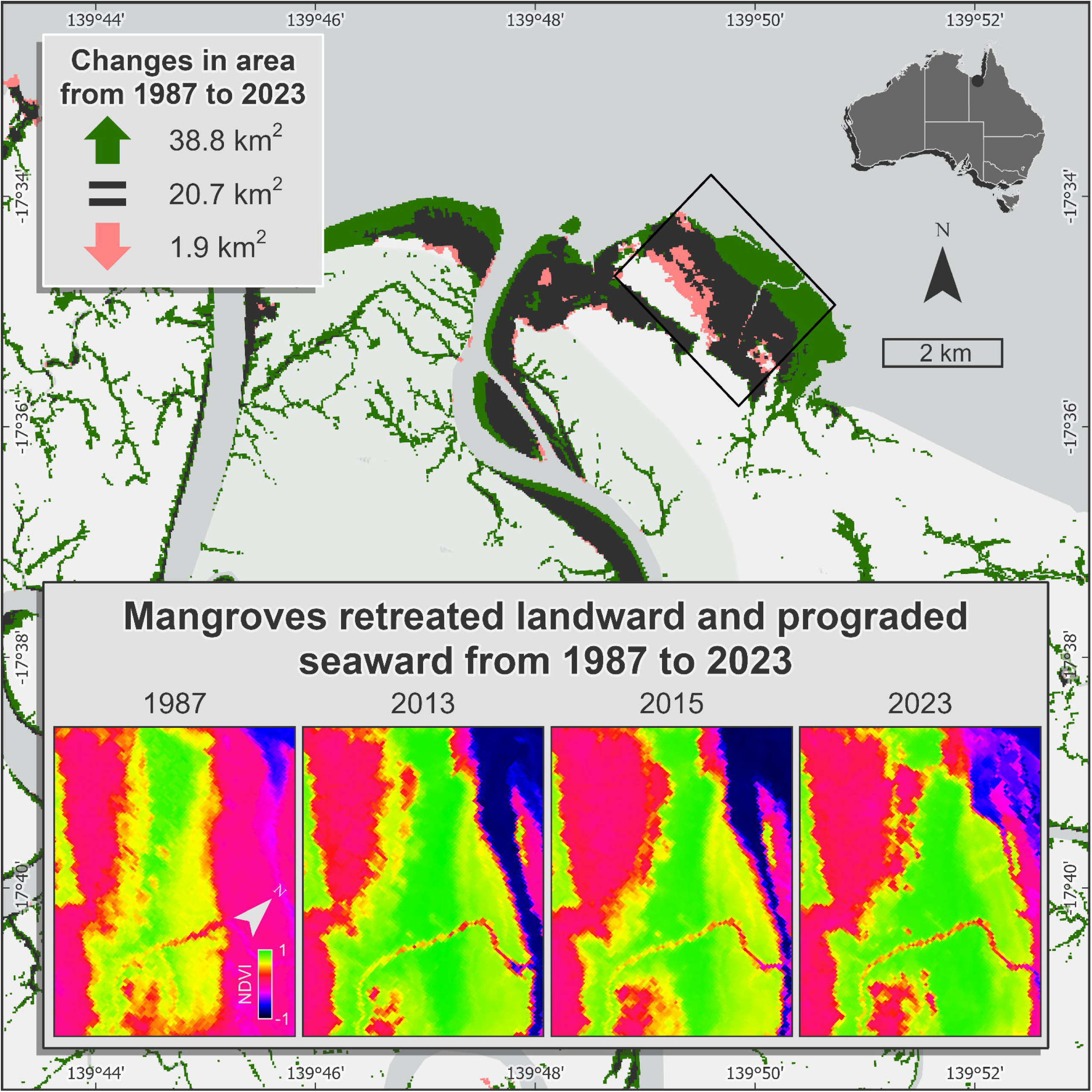

## Notes

https://github.com/rogervsg/supplementary_material_leichhardt

### Competing Interest Statement

The authors have declared no competing interest.

