## Supporting Information for "Climate-driven mangrove dieback and recovery: a case study in Albert and Leichhardt Rivers, Australia"

Supporting Information 1. Vegetation indices used for defining mangrove mask.

| **Index** | **Equation** | **Threshold value** | **Index reference** |
| --- | --- | --- | --- |
| Modular Mangrove Recognition Index (MMRI) | $\frac{(\left\vert MNDWI \right\vert-\left\vert NDVI \right\vert)}{(\left\vert MNDWI \right\vert+\vert NDVI\vert)}$ | < 0 | Diniz et al. 2019 |
| Delta Modular Mangrove Recognition Index (dMMRI) | ${MMRI}_{2016}-{MMRI}_{2015}$ | < 0.15 | NA |
| Modified Normalised Difference Water Index (MNDWI) | $\frac{(Green-SWIR2)}{(Green+SWIR2)}$ | > – 0.5 | Xu 2006 |
| Normalised Difference Water Index (NDWI) | $\frac{(Green-NIR)}{(Green+NIR)}$ | < – 0.25 | McFeeters 1996 |

Supporting Information 2. Flight details and RPA pre-processing information

| **Survey name:** LEIC1 | | | |
| --- | --- | --- | --- |
| **RPA and sensor:** Phantom / RGB | | | |
| **dRTK height (mm):** 1360 | | | |
| **Date:** 19/07/2024 | | **Time:** 15:00 – 15:20 | |
| **Flight details** | | **LiDAR pre-processing details** | |
| Average altitude (m) | 73.6 | DJI Terra software version | NA |
| Area flown (km^2^) | 0.12 | Cloud point density | NA |
| Independent GCPs | 4 | Optimize Point Cloud Accuracy | NA |
| Side and forward overlap | 80% | Smooth Point Cloud | NA |
| Return mode | NA | Projected Coordinate System (EPSG) | NA |
| LiDAR sampling rate | NA | Returns | NA |
|  |  | Outputs |  |
| **Survey name:** LEIC2 | | | |
| **RPA and sensor:** Phantom / RGB | | | |
| **dRTK height (mm):** 1505 | | | |
| **Date:** 20/07/2024 | | **Time:** 14:20 – 15:00 | |
| **Flight details** | | **LiDAR pre-processing details** | |
| Average altitude (m) | 73.6 | DJI Terra software version | NA |
| Area flown (km^2^) | 0.28 | Cloud point density | NA |
| Independent GCPs | 4 | Optimize Point Cloud Accuracy | NA |
| Side and forward overlap | 80% | Smooth Point Cloud | NA |
| Return mode | NA | Projected Coordinate System (EPSG) | NA |
| LiDAR sampling rate | NA | Returns | NA |
|  |  | Outputs | NA |
| **Survey name:** LEIC10 | | | |
| **RPA and sensor:** M300 / RGB and LiDAR | | | |
| **dRTK height (mm):** 1610 | | | |
| **Date:** 20/07/2024 | | **Time:** 11:15 – 12:00 | |
| **Flight details** | | **LiDAR pre-processing details** | |
| Average altitude (m) | 61 | DJI Terra software version | 3.7.6 |
| Area flown (km^2^) | 0.508 | Cloud point density | High |
| Independent GCPs | 4 | Optimize Point Cloud Accuracy | Yes |
| Side and forward overlap | 60% | Smooth Point Cloud | Yes |
| Return mode | Triple | Projected Coordinate System (EPSG) | WGS 84 / UTM zone 54S |
| LiDAR sampling rate | 160 KHz | Returns | 3 |
|  |  | Outputs | LAS file |
| **Survey name:** LEIC11 | | | |
| **RPA and sensor:** M300 / RGB and LiDAR | | | |
| **dRTK height (mm):** 1610 | | | |
| **Date:** 20/07/2024 | | **Time:** 12:30 – 1310 | |
| **Flight details** | | **LiDAR pre-processing details** | |
| Average altitude (m) | 62.7 | DJI Terra software version | 3.7.6 |
| Area flown (km^2^) | 0.288 | Cloud point density | High |
| Independent GCPs | 4 | Optimize Point Cloud Accuracy | Yes |
| Side and forward overlap | 60% | Smooth Point Cloud | Yes |
| Return mode | Triple | Projected Coordinate System (EPSG) | WGS 84 / UTM zone 54S |
| LiDAR sampling rate | 160 KHz | Returns | 3 |
|  |  | Outputs | LAS file |
| **Survey name:** LEIC13 | | | |
| **RPA and sensor:** M300 / RGB and LiDAR | | | |
| **dRTK height (mm):** 1625 | | | |
| **Date:** 18/07/2024 | | **Time:** 13:15 – 14:15 | |
| **Flight details** | | **LiDAR pre-processing details** | |
| Average altitude (m) | 62.2 | DJI Terra software version | 3.7.6 |
| Area flown (km^2^) | 0.406 | Cloud point density | High |
| Independent GCPs | 4 | Optimize Point Cloud Accuracy | Yes |
| Side and forward overlap | 60% | Smooth Point Cloud | Yes |
| Return mode | Triple | Projected Coordinate System (EPSG) | WGS 84 / UTM zone 54S |
| LiDAR sampling rate | 160 KHz | Returns | 3 |
|  |  | Outputs | LAS file |
| **Survey name:** LEIC14 | | | |
| **RPA and sensor:** M300 / RGB and LiDAR | | | |
| **dRTK height (mm):** 1625 | | | |
| **Date:** 18/07/2024 | | **Time:** 14:25 – 14:45 | |
| **Flight details** | | **LiDAR pre-processing details** | |
| Average altitude (m) | 62 | DJI Terra software version | 3.7.6 |
| Area flown (km^2^) | 0.21 | Cloud point density | High |
| Independent GCPs | 4 | Optimize Point Cloud Accuracy | Yes |
| Side and forward overlap | 60% | Smooth Point Cloud | Yes |
| Return mode | Triple | Projected Coordinate System (EPSG) | WGS 84 / UTM zone 54S |
| LiDAR sampling rate | 160 KHz | Returns | 3 |
|  |  | Outputs | LAS file |
| **Survey name:** LEIC15 | | | |
| **RPA and sensor:** M300 / RGB and LiDAR | | | |
| **dRTK height (mm):** 1625 | | | |
| **Date:** 18/07/2024 | | **Time:** 14:55 – 15:50 | |
| **Flight details** | | **LiDAR pre-processing details** | |
| Average altitude (m) | 62.4 | DJI Terra software version | 3.7.6 |
| Area flown (km^2^) | 0.353 | Cloud point density | High |
| Independent GCPs | 4 | Optimize Point Cloud Accuracy | Yes |
| Side and forward overlap | 60% | Smooth Point Cloud | Yes |
| Return mode | Triple | Projected Coordinate System (EPSG) | WGS 84 / UTM zone 54S |
| LiDAR sampling rate | 160 KHz | Returns | 3 |
|  |  | Outputs | LAS file |
| **Survey name:** LEIC16 | | | |
| **RPA and sensor:** M300 / RGB and LiDAR | | | |
| **dRTK height (mm):** 1640 | | | |
| **Date:** 19/07/2024 | | **Time:** 11:00 – 11:55 | |
| **Flight details** | | **LiDAR pre-processing details** | |
| Average altitude (m) | 61.8 | DJI Terra software version | 3.7.6 |
| Area flown (km^2^) | 0.284 | Cloud point density | High |
| Independent GCPs | 4 | Optimize Point Cloud Accuracy | Yes |
| Side and forward overlap | 60% | Smooth Point Cloud | Yes |
| Return mode | Triple | Projected Coordinate System (EPSG) | WGS 84 / UTM zone 54S |
| LiDAR sampling rate | 160 KHz | Returns | 3 |
|  |  | Outputs | LAS file |
| **Survey name:** LEIC17 | | | |
| **RPA and sensor:** M300 / RGB and LiDAR | | | |
| **dRTK height (mm):** 1640 | | | |
| **Date:** 19/07/2024 | | **Time:** 12:30 – 13:50 | |
| **Flight details** | | **LiDAR pre-processing details** | |
| Average altitude (m) | 63.5 | DJI Terra software version | 3.7.6 |
| Area flown (km^2^) | 0.719 | Cloud point density | High |
| Independent GCPs | 4 | Optimize Point Cloud Accuracy | Yes |
| Side and forward overlap | 60% | Smooth Point Cloud | Yes |
| Return mode | Triple | Projected Coordinate System (EPSG) | WGS 84 / UTM zone 54S |
| LiDAR sampling rate | 160 KHz | Returns | 3 |
|  |  | Outputs | LAS file |

Supporting Information 3. Confusion matrix for the 2016 preliminary classification data.

|  |  | **Reference data** | | | | | | | |  | Total | User Accuracy |
| --- | --- | --- | --- | --- | --- | --- | --- | --- | --- | --- | --- | --- |
|  | **Class** | *Ceriops* | Closed *Avicennia* | *Rhizophora* | Open Mixed | Closed Mixed | Open *Avicennia* | Open Dieback | Closed Dieback | Not mangrove |  |  |
| **Classification data** | *Ceriops* | 10 | 1 | 0 | 3 | 11 | 0 | 0 | 0 | 2 | 27 | 0.37 |
|  | Closed *Avicennia* | 0 | 24 | 3 | 0 | 1 | 0 | 0 | 0 | 0 | 28 | 0.86 |
|  | *Rhizophora* | 0 | 1 | 18 | 1 | 4 | 0 | 0 | 0 | 0 | 24 | 0.75 |
|  | Open Mixed | 1 | 0 | 0 | 13 | 8 | 0 | 0 | 0 | 7 | 29 | 0.45 |
|  | Closed Mixed | 0 | 0 | 0 | 8 | 13 | 1 | 0 | 0 | 5 | 27 | 0.48 |
|  | Open *Avicennia* | 0 | 5 | 0 | 0 | 0 | 16 | 0 | 0 | 6 | 27 | 0.59 |
|  | Open Dieback | 0 | 0 | 0 | 0 | 0 | 0 | 18 | 1 | 1 | 20 | 0.90 |
|  | Closed Dieback | 0 | 0 | 0 | 0 | 0 | 0 | 5 | 21 | 0 | 26 | 0.81 |
|  | Not mangrove | 0 | 0 | 0 | 0 | 0 | 0 | 0 | 0 | 30 | 30 | 1.00 |
|  | Total | 11 | 31 | 21 | 25 | 37 | 17 | 23 | 22 | 51 | 238 |  |
|  | Producer Accuracy | 0.91 | 0.77 | 0.86 | 0.52 | 0.35 | 0.94 | 0.78 | 0.95 | 0.59 |  | **0.68** |

Supporting Information 4. Classification report for test data and predictions.

| **Class** | **Precision** | **Recall** | **F1-score** | **Support** |
| --- | --- | --- | --- | --- |
| *Ceriops* | 0.944 | 0.95 | 0.94 | 291 |
| Closed *Avicennia* | 0.98 | 0.99 | 0.99 | 269 |
| *Rhizophora* | 0.98 | 0.96 | 0.97 | 333 |
| Open Mixed | 0.97 | 0.94 | 0.96 | 286 |
| Closed Mixed | 0.96 | 0.95 | 0.95 | 313 |
| Open *Avicennia* | 0.98 | 1.00 | 0.99 | 308 |
| Open Dieback | 0.98 | 1.00 | 0.99 | 300 |
| Closed Dieback | 0.97 | 0.99 | 0.98 | 300 |
| Micro avg |  |  | 0.97 | 2400 |
| Macro avg | 0.97 | 0.97 | 0.97 | 2400 |
| Weighted avg | 0.97 | 0.97 | 0.97 | 2400 |

Supporting Information 5. Confusion matrix for test data and predictions.

|  |  | **Predicted label** | | | | | | | |  |
| --- | --- | --- | --- | --- | --- | --- | --- | --- | --- | --- |
|  | **Class** | *Ceriops* | Closed *Avicennia* | *Rhizophora* | Open Mixed | Closed Mixed | Open *Avicennia* | Open Dieback | Closed Dieback | Total |
| **True label** | *Ceriops* | 275 | 3 | 4 | 0 | 5 | 1 | 0 | 3 | 291 |
|  | Closed *Avicennia* | 1 | 267 | 1 | 0 | 0 | 0 | 0 | 0 | 269 |
|  | *Rhizophora* | 10 | 3 | 320 | 0 | 0 | 0 | 0 | 0 | 333 |
|  | Open Mixed | 0 | 0 | 0 | 269 | 5 | 3 | 5 | 4 | 286 |
|  | Closed Mixed | 7 | 0 | 0 | 6 | 296 | 3 | 0 | 1 | 313 |
|  | Open *Avicennia* | 0 | 0 | 0 | 0 | 0 | 308 | 0 | 0 | 308 |
|  | Open Dieback | 0 | 0 | 0 | 1 | 0 | 0 | 299 | 0 | 300 |
|  | Closed Dieback | 0 | 0 | 0 | 1 | 1 | 0 | 0 | 298 | 300 |
| Total | | 293 | 273 | 325 | 277 | 307 | 315 | 304 | 306 | 2400 |

Supporting Information 6. Model summaries for Generalised Additive Models (GAM), including adjusted R² values, deviance explained, and p-values.

| **Mangrove zone** | **Variable** | **R-sq (adj)** | **Deviance explained (%)** | **p-value** | | | | | | |
| --- | --- | --- | --- | --- | --- | --- | --- | --- | --- | --- |
|  |  |  |  | **Time (year)** | **DMI** | **NINO3.4** | **MSL** | **DMI:NINO3.4** | **DMI:MSL** | **NINO3.4:MSL** |
| *Ceriops*-dominated zone | NDVI | 0.777 | 90.7 | **0.030*** | 0.245 | **0.019*** | 0.147 | 0.107 | 0.328 | 0.212 |
|  | Area | 0.774 | 88.5 | **0.001*** | **0.019*** | 0.147 | 0.243 | **0.028*** | **0.030*** | 0.401 |
| Closed *Avicennia*-dominated zone | NDVI | 0.77 | 85.2 | 0.71 | 0.988 | 0.323 | 0.054 | 0.713 | 0.951 | 0.542 |
|  | Area | 0.462 | 66.9 | 0.256 | 1 | 0.08 | 0.34 | 0.81 | 0.433 | 0.916 |
| Closed forest dieback | NDVI | 0.765 | 90.5 | 0.203 | 0.277 | 0.156 | 0.342 | **0.004*** | 0.131 | 0.117 |
|  | Area | 0.426 | 57.0 | **0.010*** | **0.005*** | **0.005*** | **0.033*** | **0.001*** | **0.017*** | 0.22 |
| Closed mixed zone | NDVI | 0.751 | 85.2 | 0.321 | 0.888 | 0.125 | 0.39 | 0.241 | 0.304 | 0.955 |
|  | Area | 0.435 | 62.8 | 0.159 | 0.271 | 0.095 | 0.089 | 0.596 | **0.042*** | 0.199 |
| Open *Avicennia*-dominated zone | NDVI | 0.851 | 91.7 | **0.003*** | 0.619 | 0.083 | 0.157 | 0.056 | 0.118 | 0.304 |
|  | Area | 0.396 | 64.5 | 0.178 | 0.225 | 0.693 | 0.163 | 0.153 | 0.705 | 0.379 |
| Open forest dieback | NDVI | 0.961 | 98.0 | **0.000*** | 0.961 | 0.664 | 0.216 | 0.348 | 0.628 | 0.254 |
|  | Area | -0.084 | 30.8 | 0.643 | 0.436 | 0.936 | 0.644 | 0.928 | 0.644 | 0.921 |
| Open mixed zone | NDVI | 0.855 | 91.5 | 0.174 | 0.693 | **0.027*** | 0.103 | 0.051 | 0.158 | 0.529 |
|  | Area | -0.0868 | 27.9 | 0.723 | 0.355 | 0.777 | 0.925 | 0.165 | **0.045*** | 0.403 |
| *Rhizophora*-dominated zone | NDVI | 0.483 | 69.3 | 0.148 | 0.746 | 0.068 | 0.667 | 0.737 | 0.888 | 0.506 |
|  | Area | 0.782 | 87.6 | 0.218 | 0.769 | 0.269 | 0.275 | 0.178 | **0.007*** | **0.034*** |

Supporting Information 7. Model summaries for Generalised Additive Models (GAM), including effective degrees of freedom (EDF).

| **Mangrove zone** | **Variable** | **EDF** | | | | | | |
| --- | --- | --- | --- | --- | --- | --- | --- | --- |
|  |  | **Time (year)** | **DMI** | **NINO3.4** | **MSL** | **DMI:NINO3.4** | **DMI:MSL** | **NINO3.4:MSL** |
| *Ceriops*-dominated zone | NDVI | 3.383 | 2 | 2 | 2 | 1 | 1.174 | 8.247 |
|  | Area | 6.826 | 2 | 2 | 2 | 1 | 1 | 1.836 |
| Closed *Avicennia*-dominated zone | NDVI | 1 | 2 | 2.538 | 3.52 | 1 | 1 | 1 |
|  | Area | 2.674 | 2 | 3.43 | 2 | 1 | 1 | 1 |
| Closed forest dieback | NDVI | 1 | 2 | 2 | 2 | 1 | 3.703 | 7.398 |
|  | Area | 0.255 | 1.998 | 1.994 | 1.662 | 0.906 | 0.955 | 0.984 |
| Closed mixed zone | NDVI | 3.496 | 2 | 3.243 | 2 | 1 | 1 | 1 |
|  | Area | 1 | 2 | 3.601 | 2 | 1 | 1 | 1 |
| Open *Avicennia*-dominated zone | NDVI | 4.989 | 2 | 3.049 | 2 | 1 | 1 | 1 |
|  | Area | 4.3 | 2.489 | 2 | 2 | 1.201 | 1 | 1 |
| Open forest dieback | NDVI | 7.028 | 2 | 2.325 | 2 | 1 | 1 | 1 |
|  | Area | 1 | 2 | 2 | 3.86 | 1 | 1 | 1.069 |
| Open mixed zone | NDVI | 3.744 | 2 | 3.378 | 2 | 1 | 1 | 1 |
|  | Area | 1 | 2 | 2.656 | 2 | 1 | 1.037 | 1.741 |
| *Rhizophora*-dominated zone | NDVI | 2.768 | 2 | 2 | 2 | 1 | 1 | 3.02 |
|  | Area | 1 | 2 | 2 | 2 | 2.814 | 3.921 | 1 |

Supporting Information 8. Confusion matrix for mangrove classification from 2023 using field verification data as a reference.

|  |  | **Field verification data** | | | | | Total | User Accuracy |
| --- | --- | --- | --- | --- | --- | --- | --- | --- |
|  | **Class** | *Ceriops* | *Avicennia* | *Rhizophora* | Closed mixed | Not mangrove |  |  |
| **Classification** **data** | *Ceriops* | 7 | 1 | 2 | 2 | 0 | 12 | 0.88 |
|  | *Avicennia* | 0 | 22 | 0 | 4 | 0 | 26 | 0.85 |
|  | *Rhizophora* | 0 | 1 | 5 | 4 | 0 | 7 | 0.71 |
|  | Closed mixed | 1 | 1 | 0 | 17 | 0 | 33 | 0.52 |
|  | Not mangrove | 0 | 1 | 0 | 6 | 4 | 4 | 1 |
|  | Total | 12 | 26 | 10 | 19 | 11 | 78 |  |
|  | Producer Accuracy | 0.58 | 0.85 | 0.5 | 0.9 | 0.36 |  | **0.71** |

Supporting Information 9. Confusion matrix for mangrove zonation from 2000 using reference data (Hay 2009).

|  |  | **Reference data** | | | | | Total | User Accuracy |
| --- | --- | --- | --- | --- | --- | --- | --- | --- |
|  |  | *Ceriops* | Closed *Avicennia* | *Rhizophora* | Closed mixed | Open *Avicennia* |  |  |
| **Classification data** | *Ceriops* | 49 | 1 | 58 | 0 | 1 | 109 | 0.45 |
|  | Closed *Avicennia* | 60 | 103 | 28 | 4 | 16 | 211 | 0.49 |
|  | *Rhizophora* | 3 | 5 | 7 | 0 | 0 | 15 | 0.47 |
|  | Closed mixed | 40 | 2 | 3 | 10 | 7 | 62 | 0.16 |
|  | Open *Avicennia* | 16 | 0 | 0 | 2 | 16 | 34 | 0.47 |
|  | Total | 168 | 111 | 96 | 16 | 40 | 431 |  |
|  | Producer Accuracy | 0.29 | 0.93 | 0.07 | 0.63 | 0.40 |  | **0.43** |

Supporting Information 10. Confusion matrix for mangrove dieback from 2016 using Duke et al. (2017) as reference data.

|  |  | **Reference data** | | Total | User Accuracy |
| --- | --- | --- | --- | --- | --- |
|  | **Class** | No dieback | Dieback |  |  |
| Classification data | No dieback | 250 | 118 | 368 | 0.68 |
|  | Dieback | 0 | 132 | 132 | 1 |
|  | Total | 250 | 250 | 500 |  |
|  | Producer Accuracy | 1 | 0.53 |  | **0.76** |

Supporting Information 14. High resolution imagery from 1988 and 2017, Hovmöller for NDVI and mangrove classification zones for transects 1 to 57.

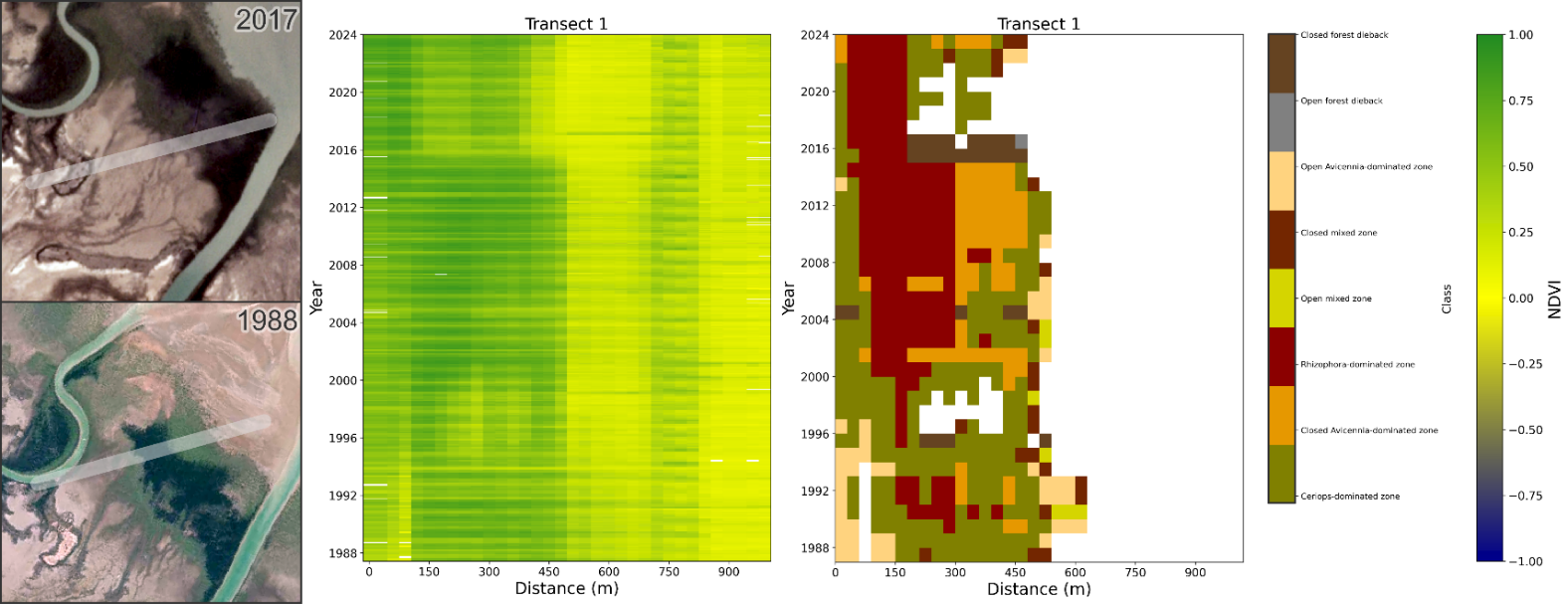

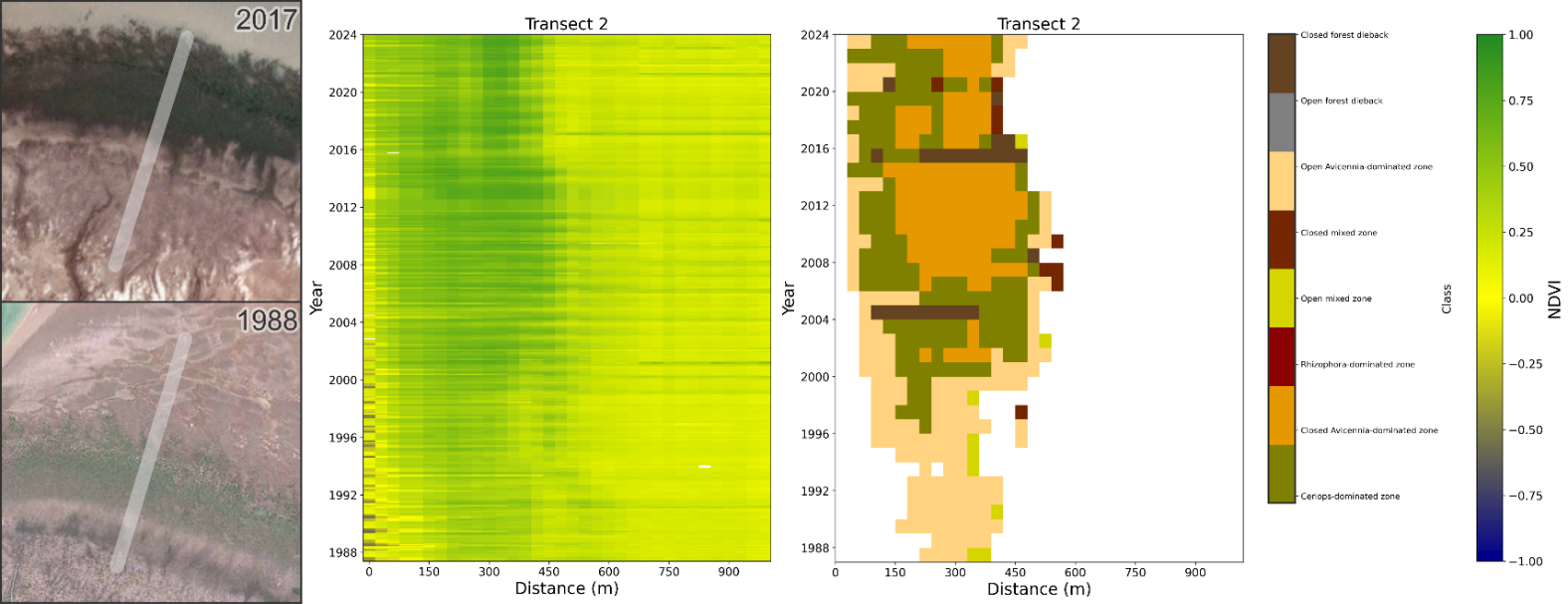

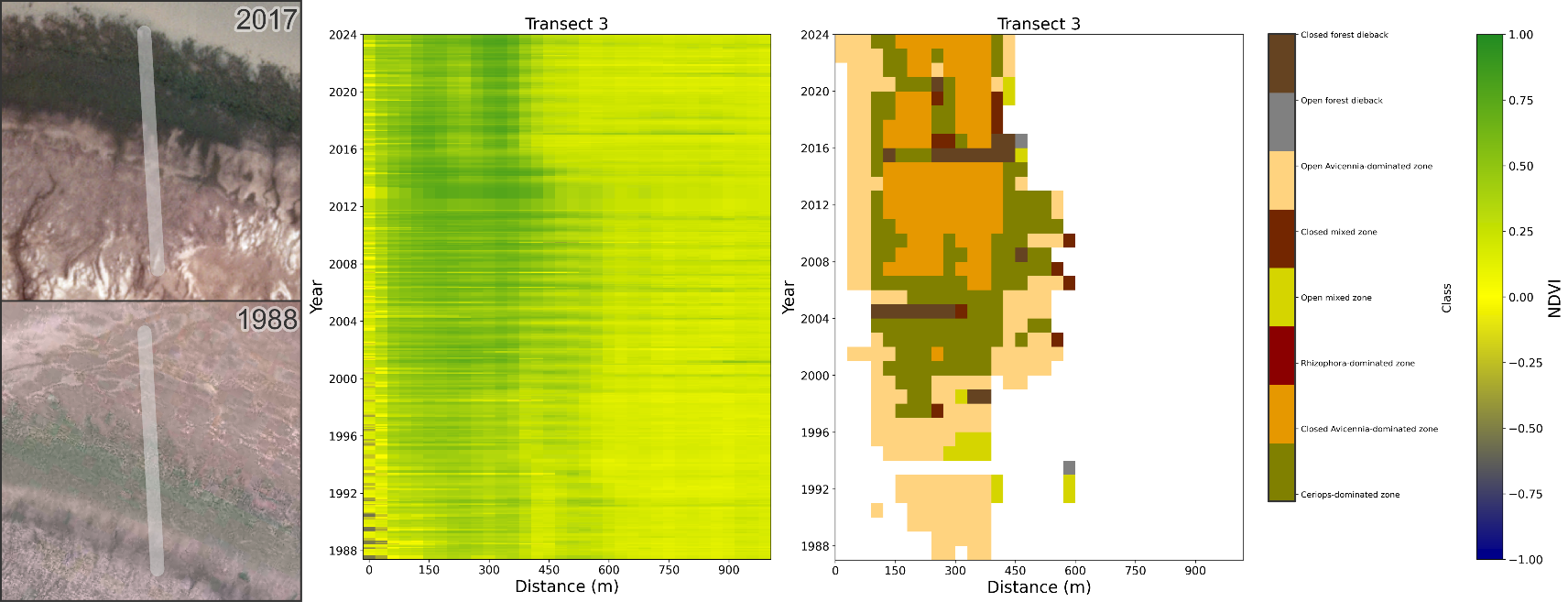

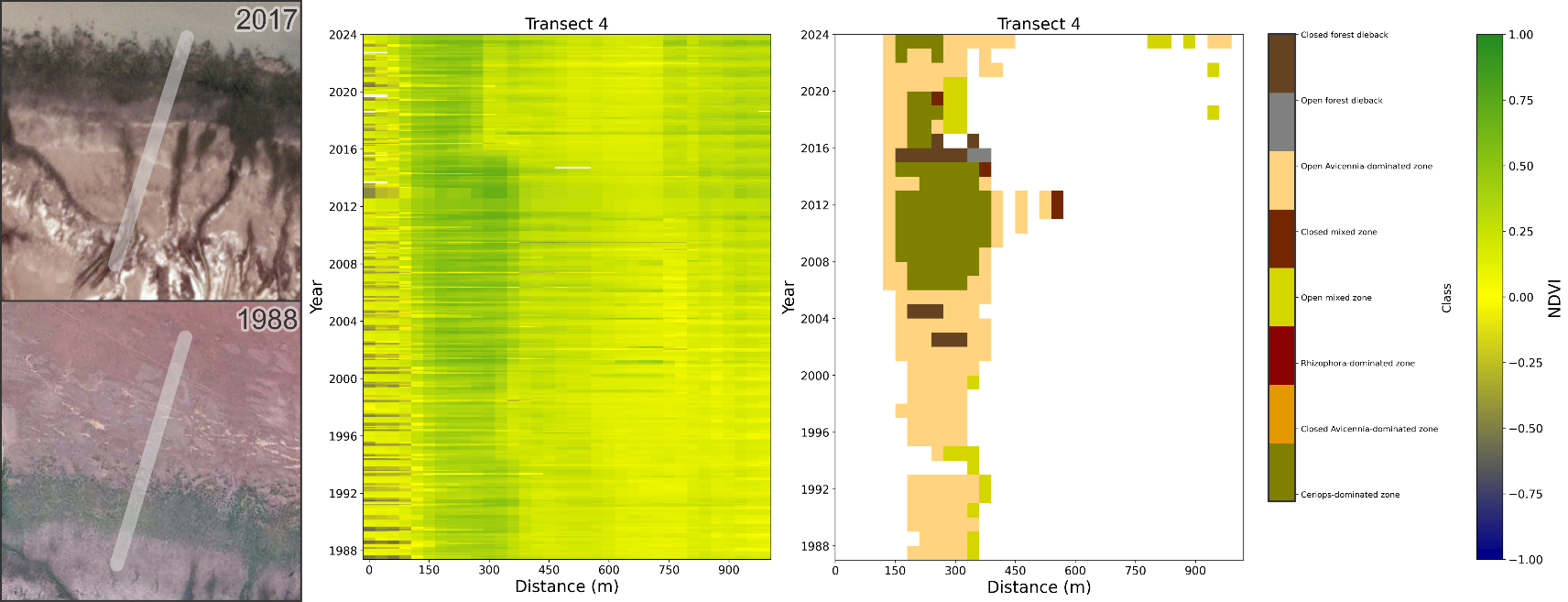

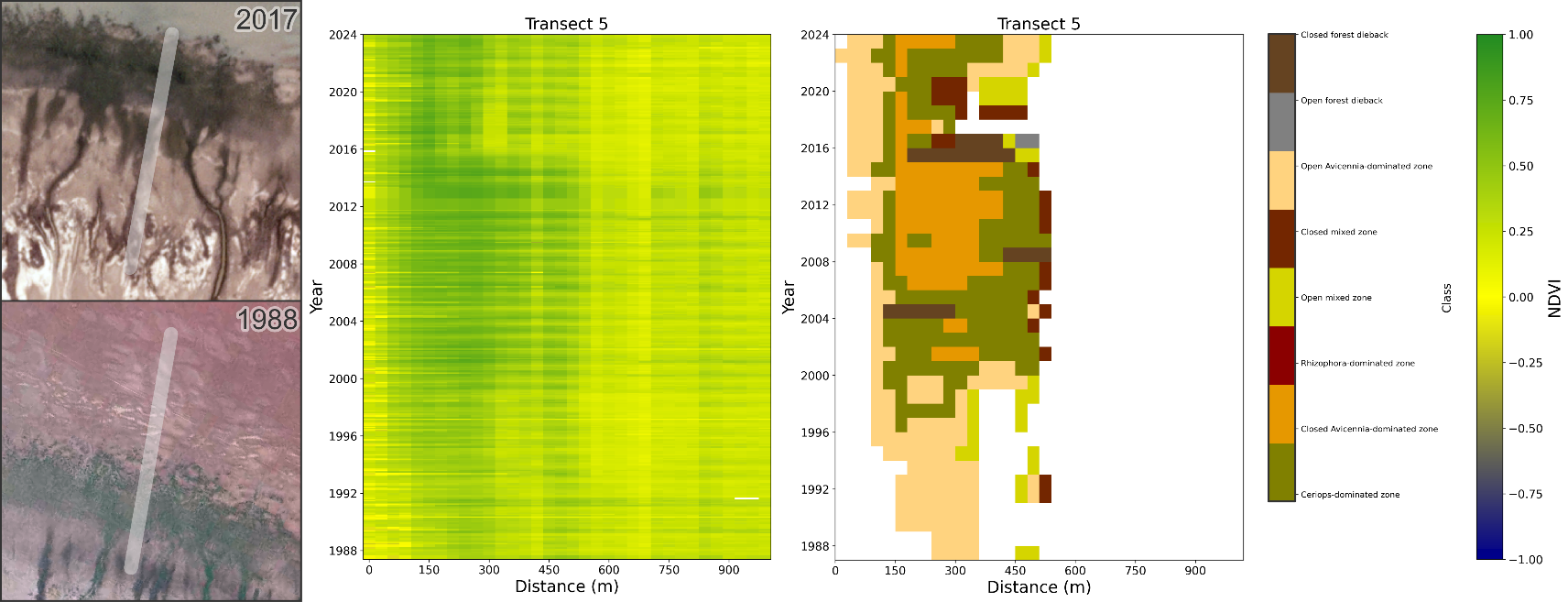

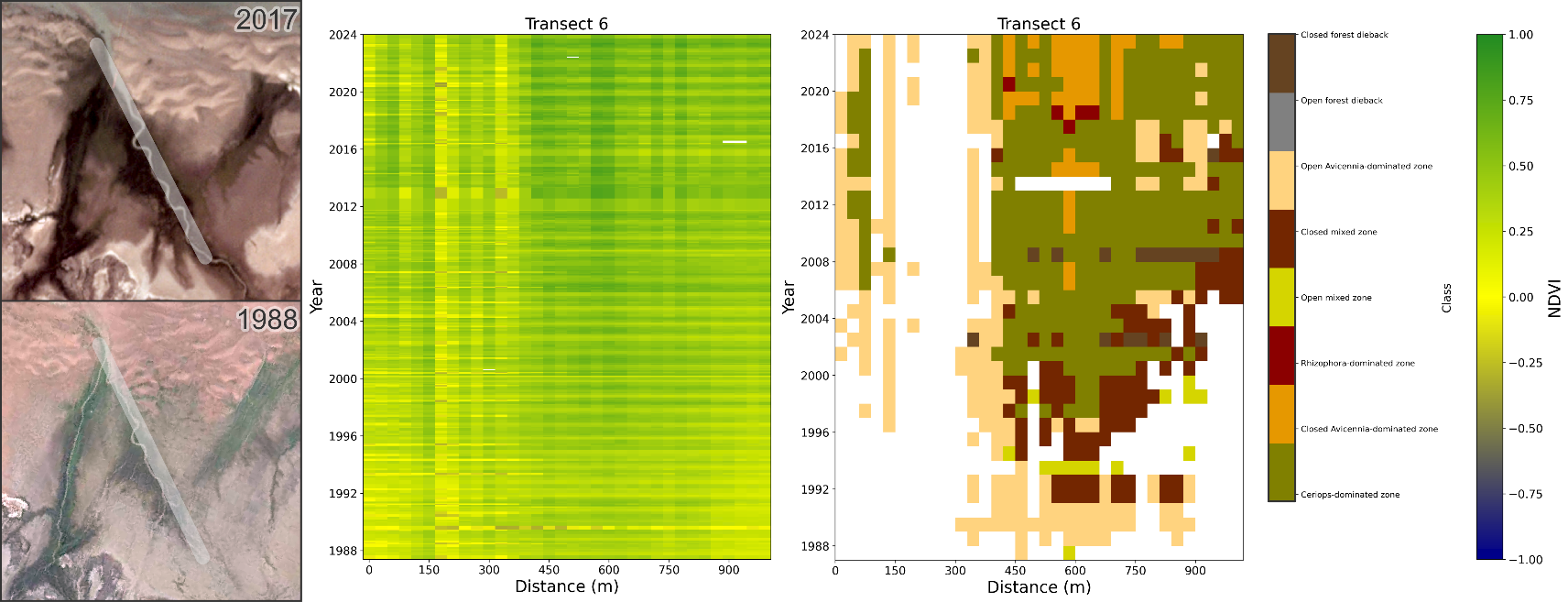

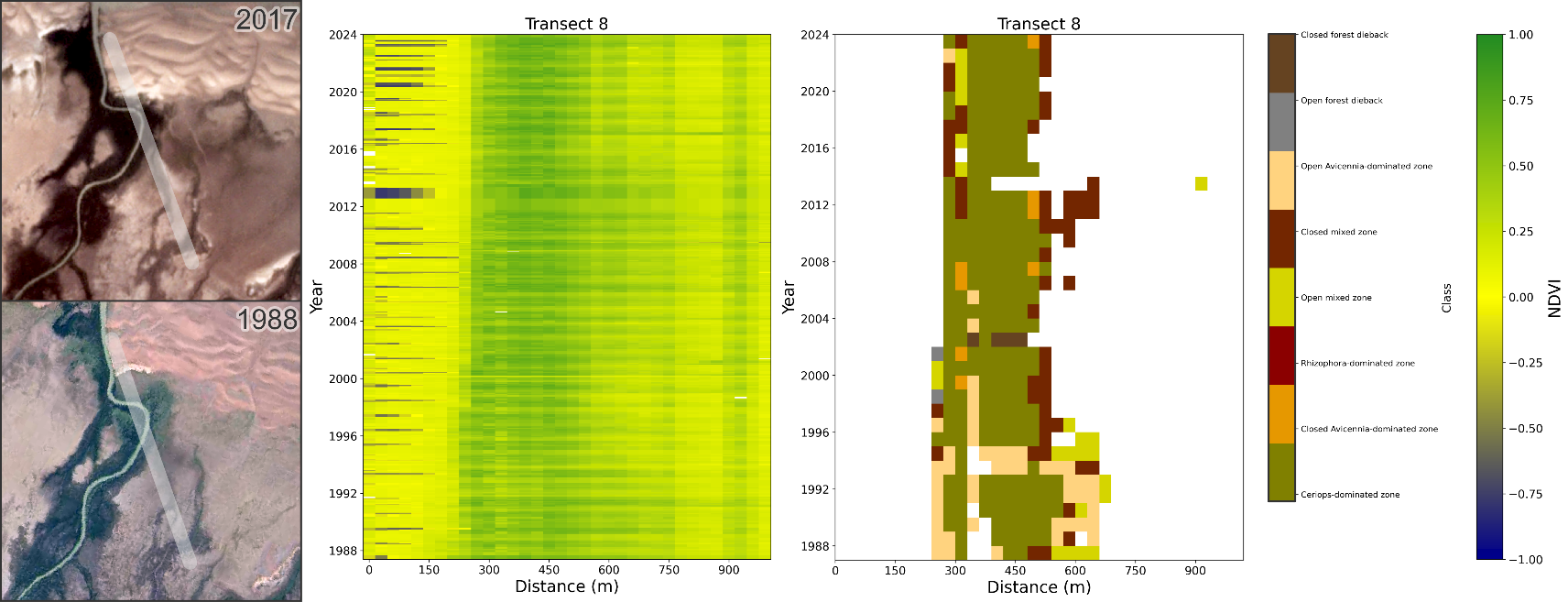

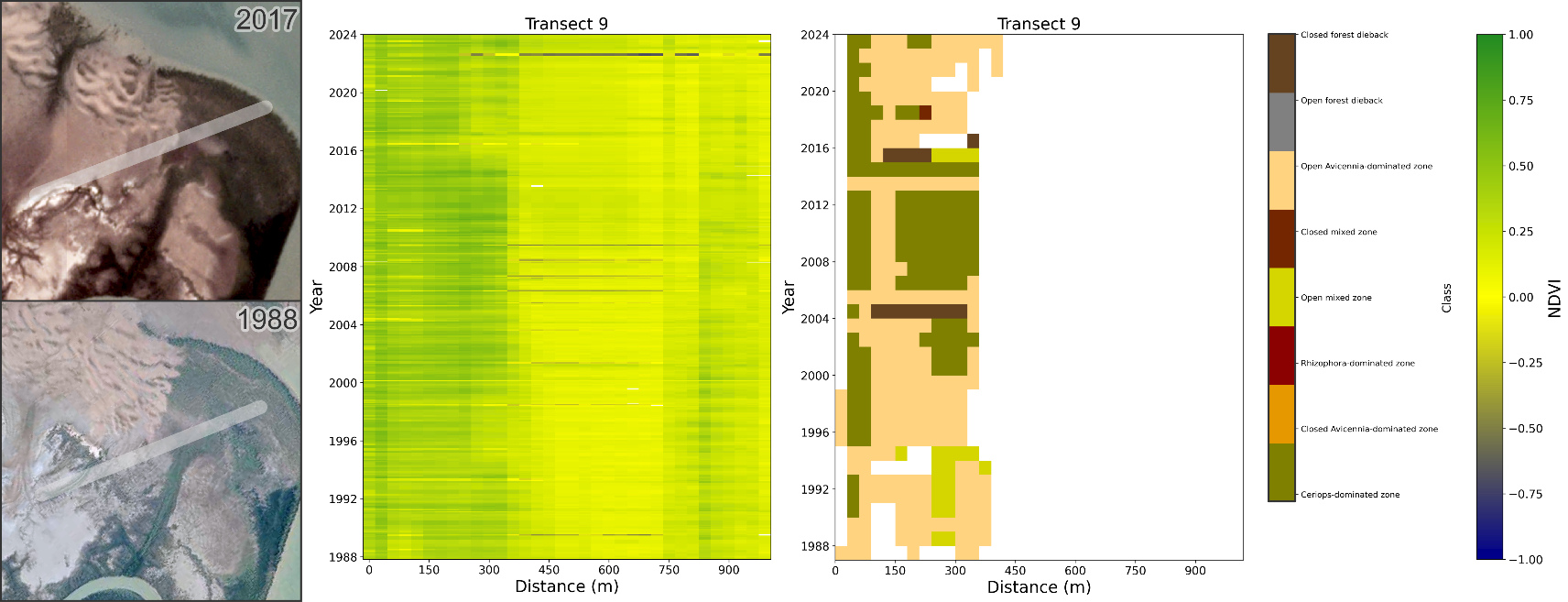

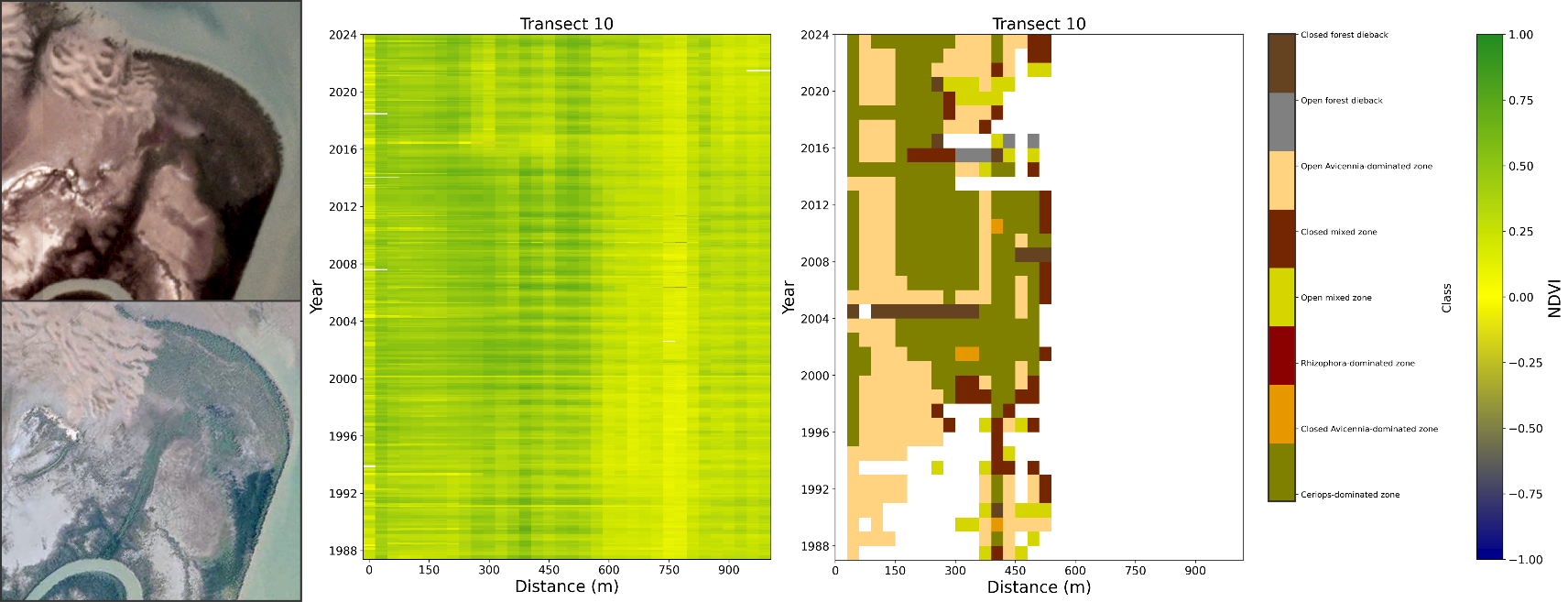

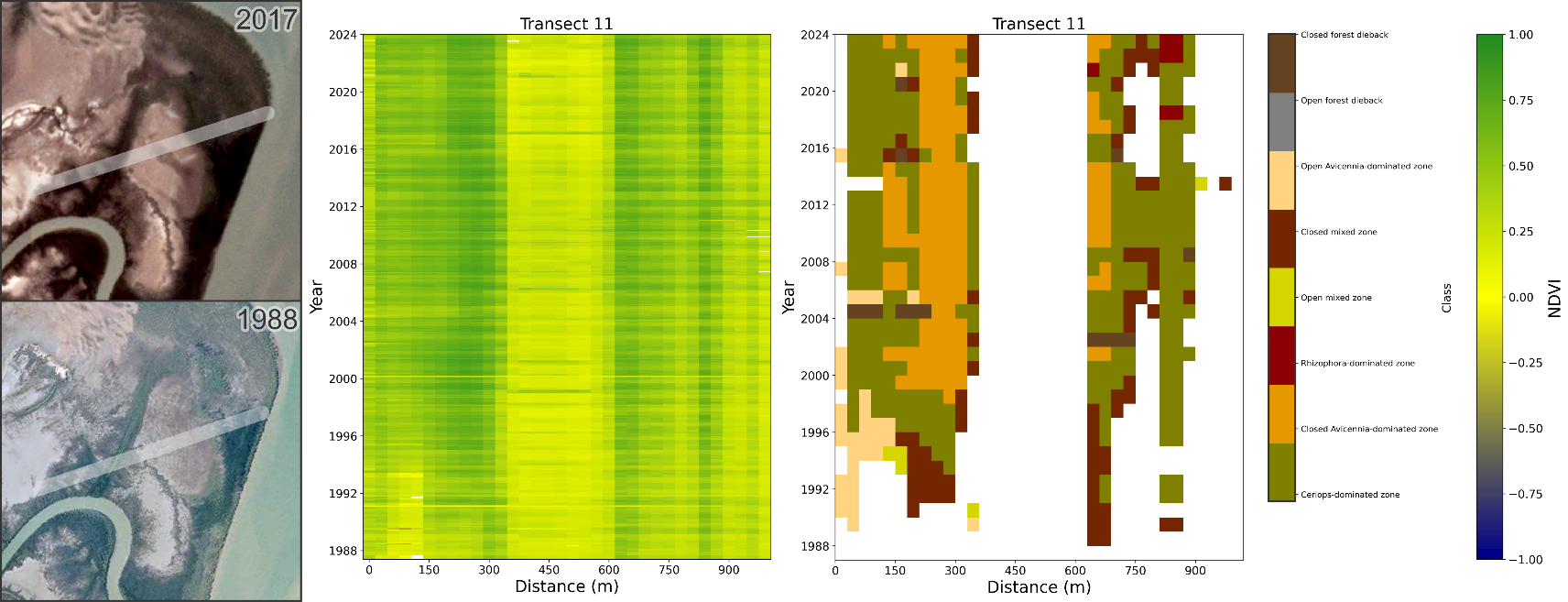

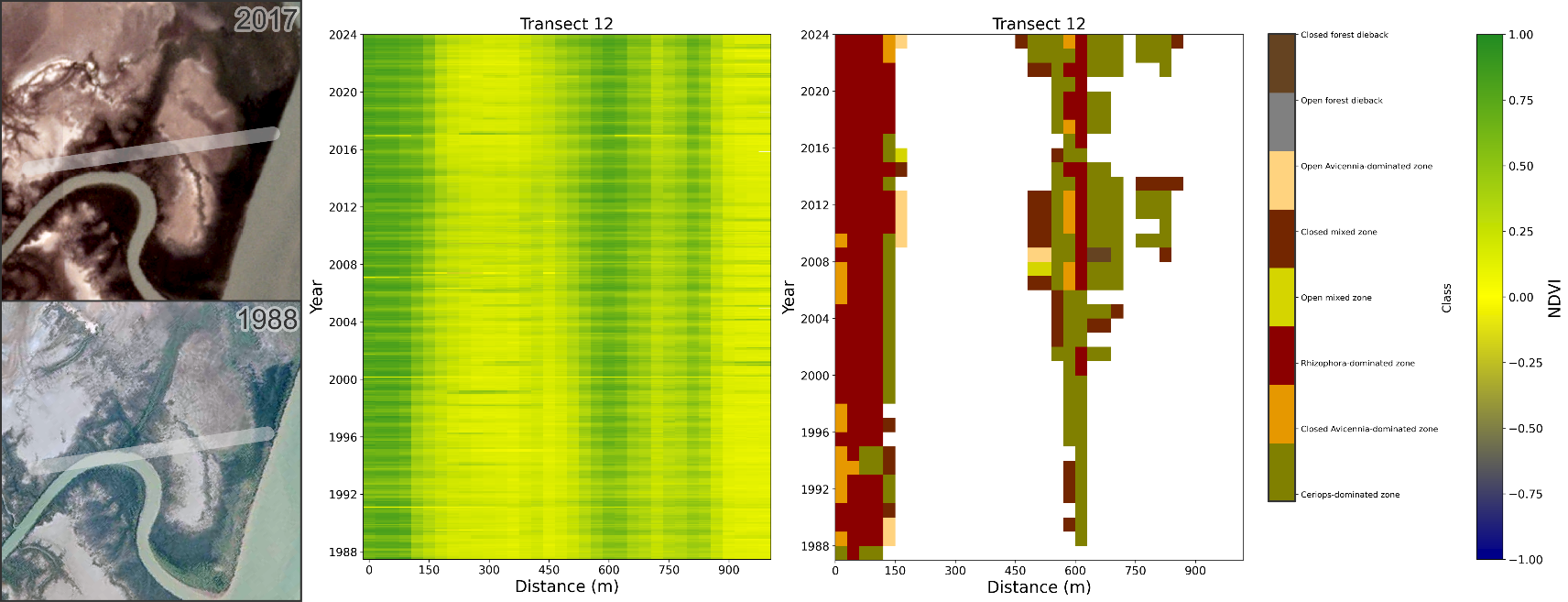

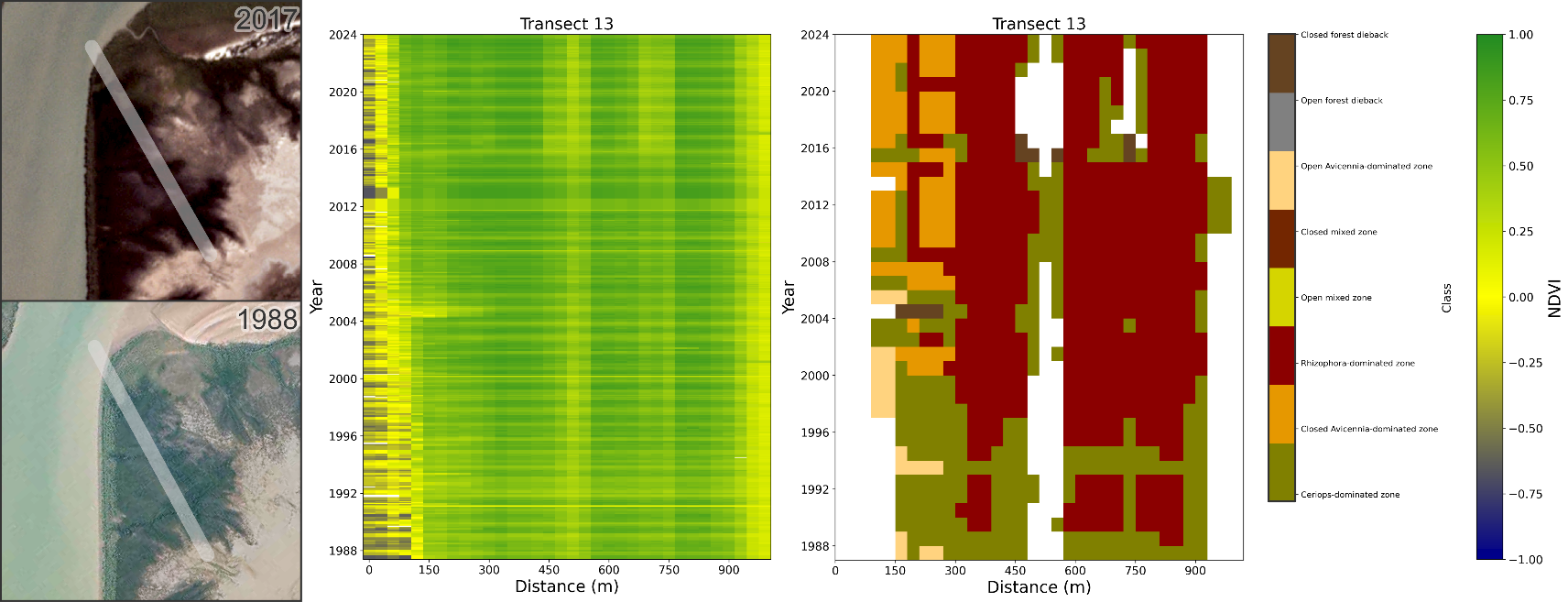

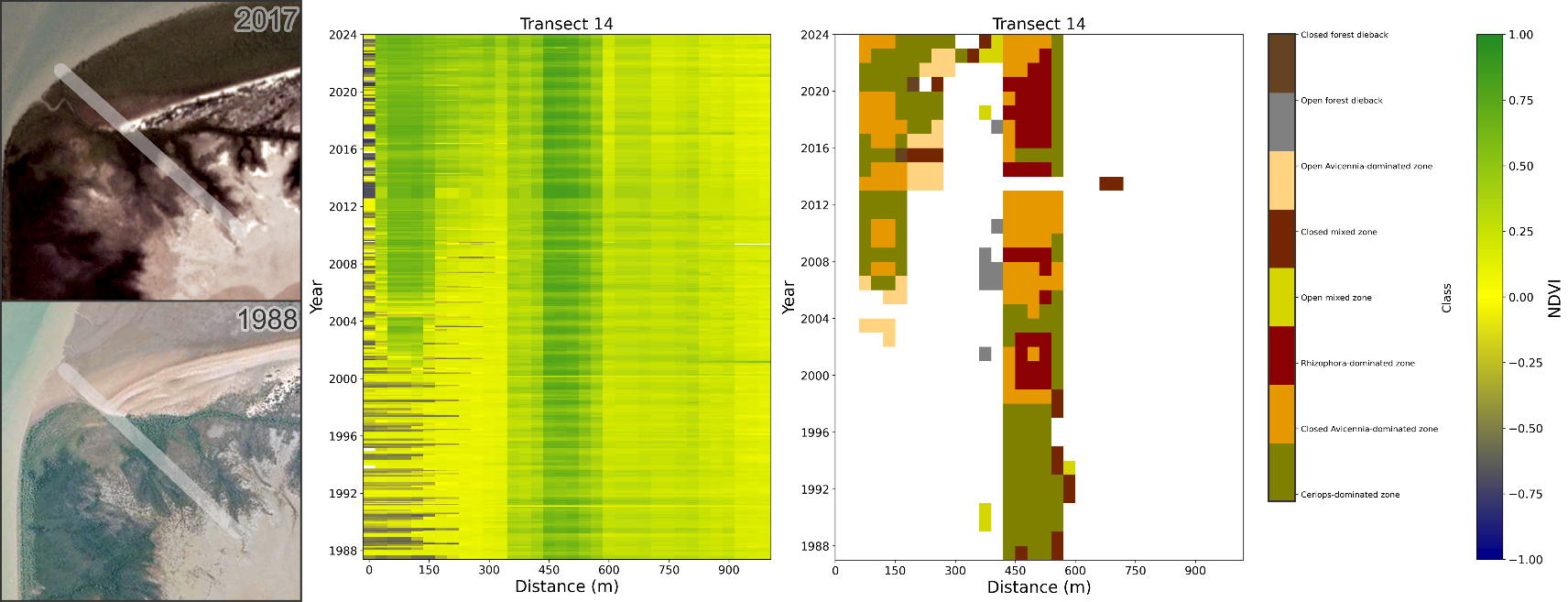

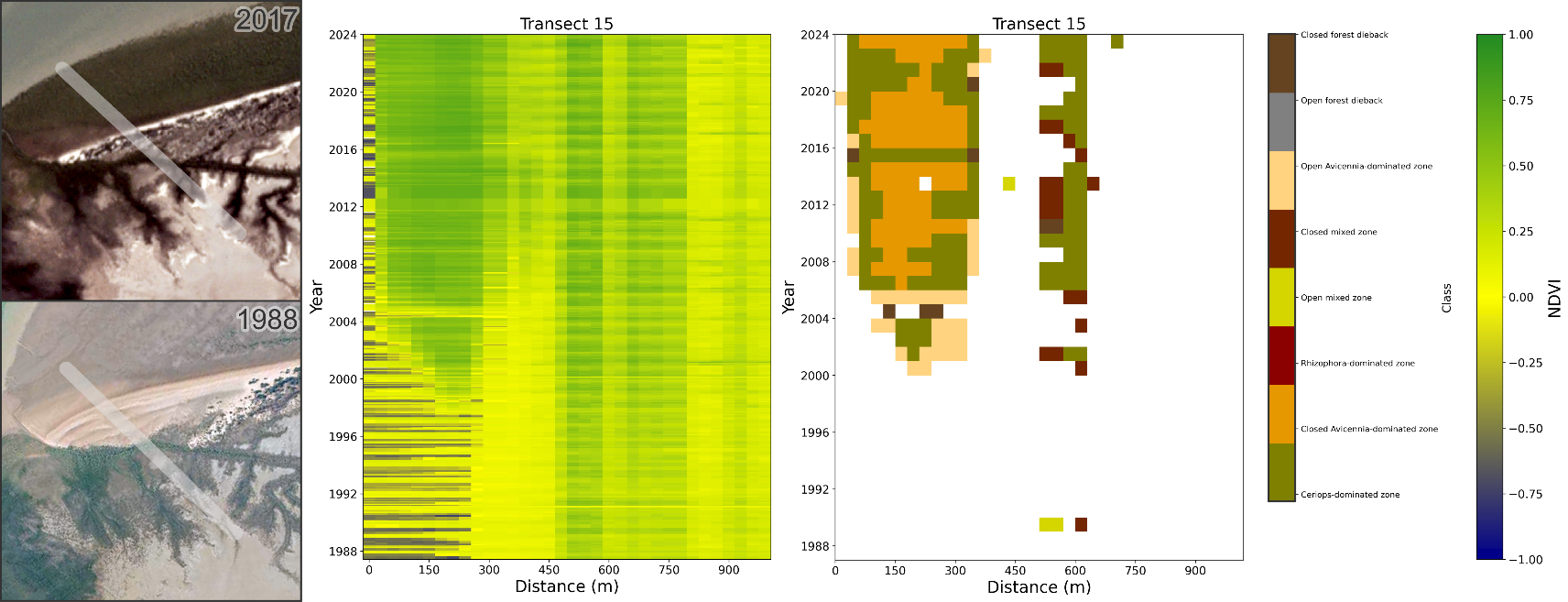

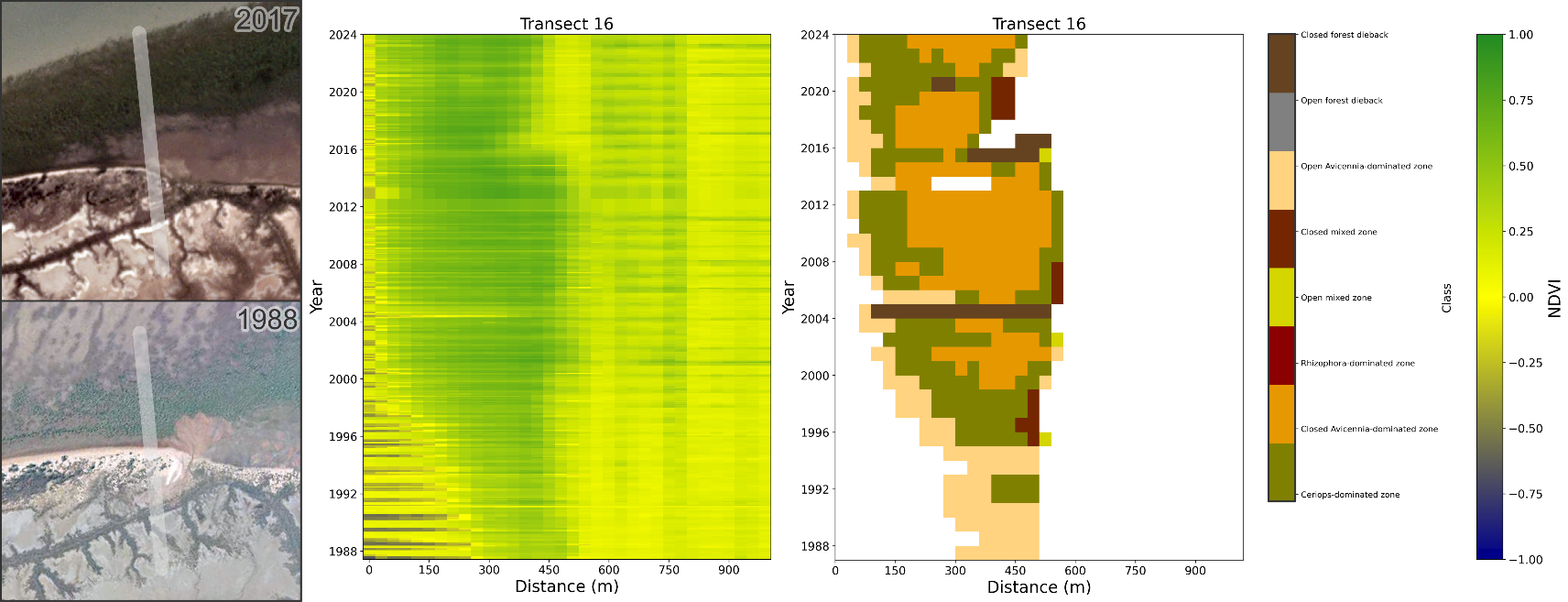

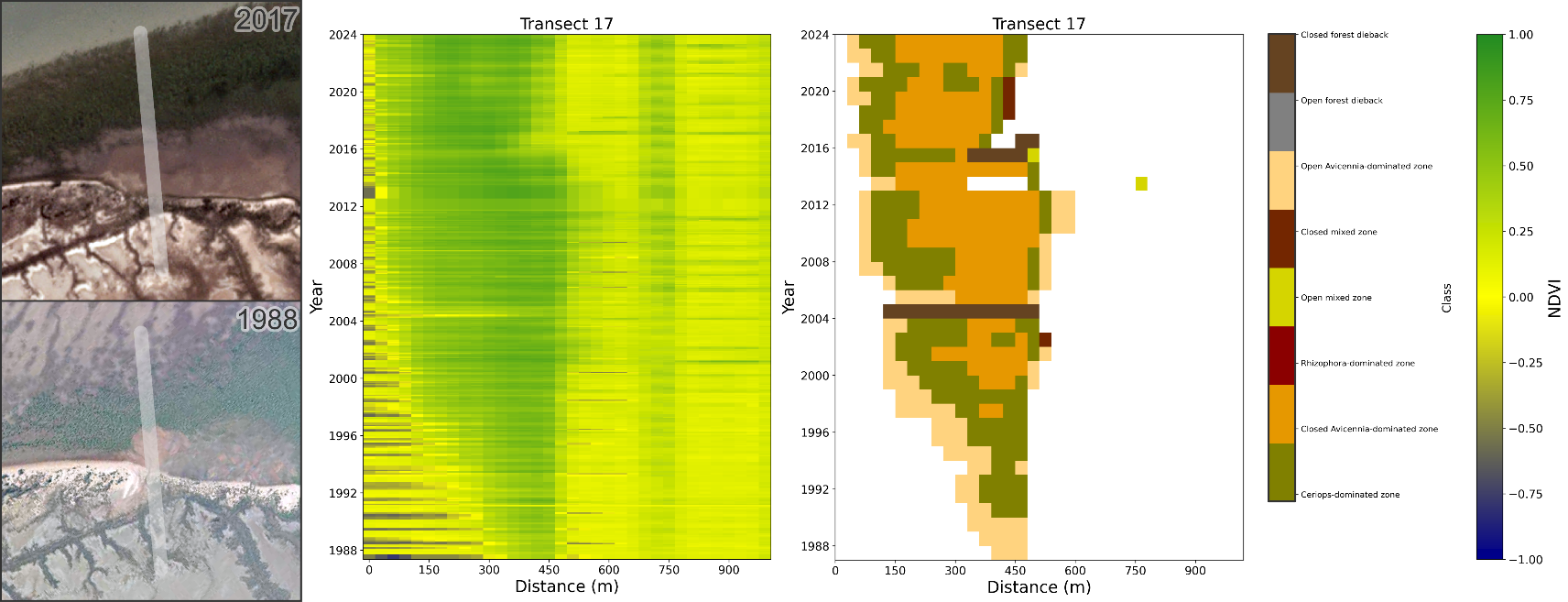

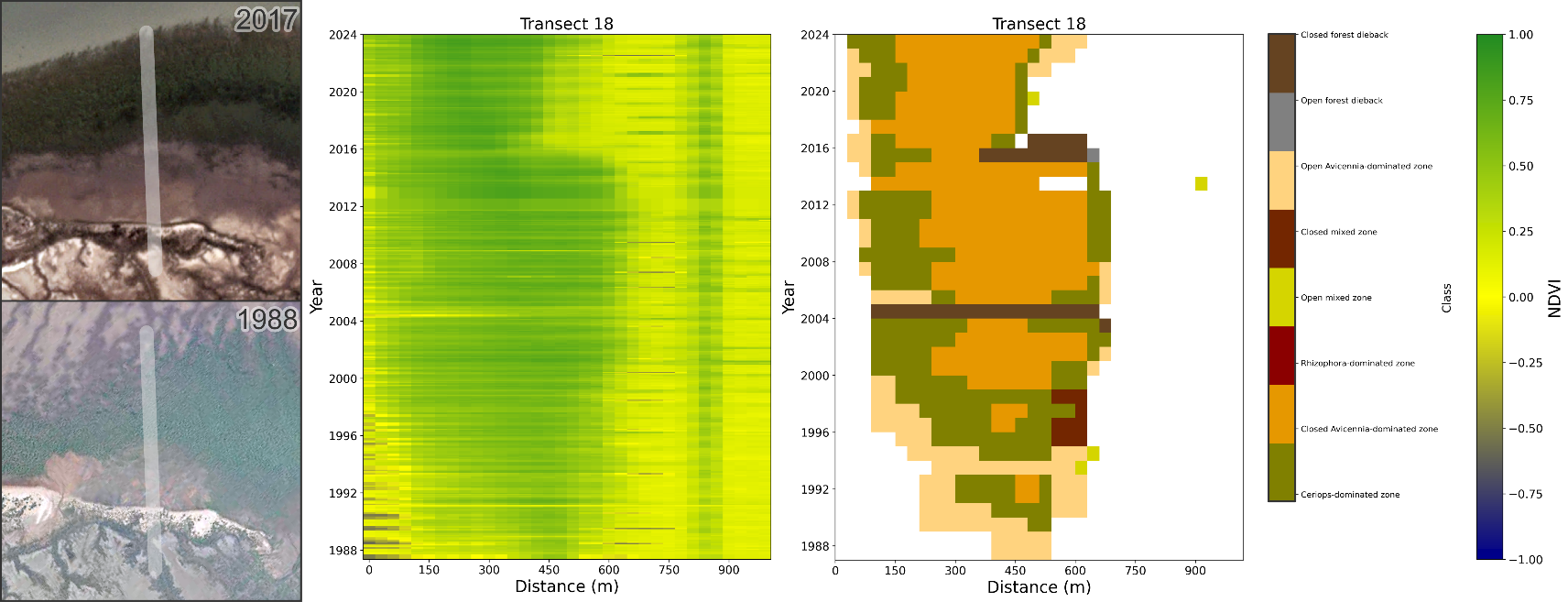

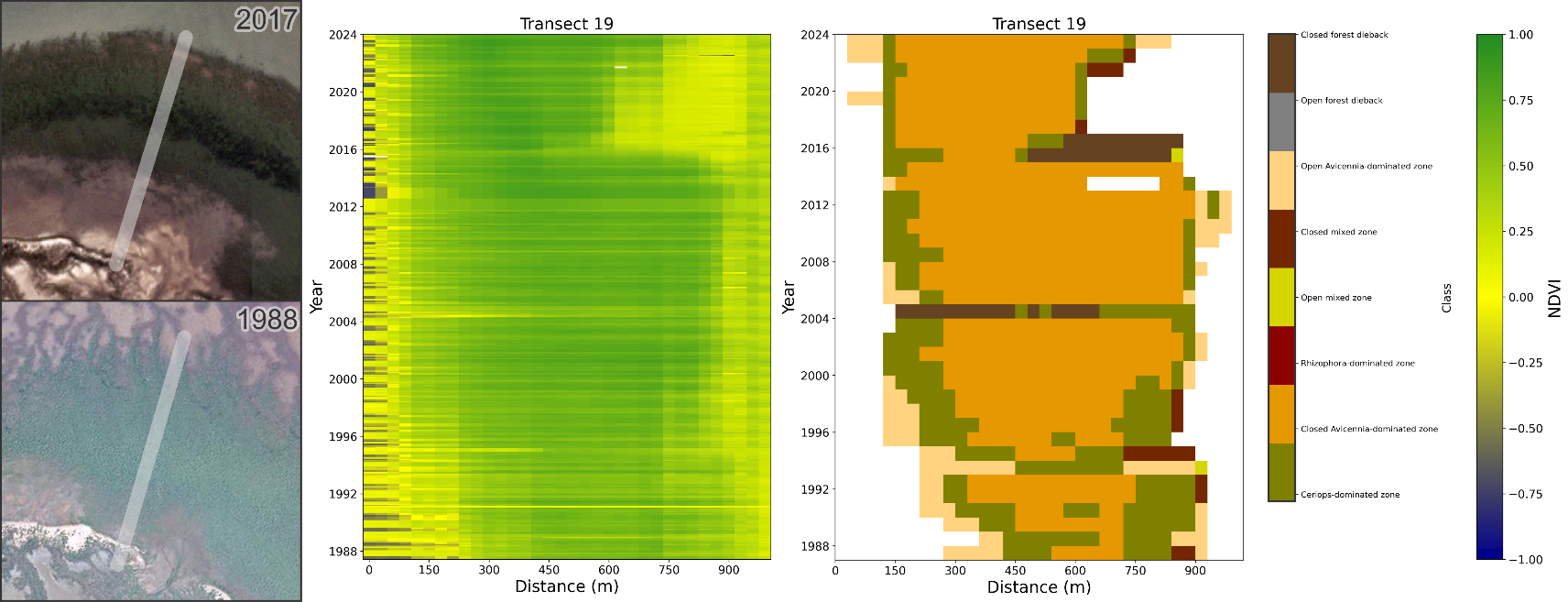

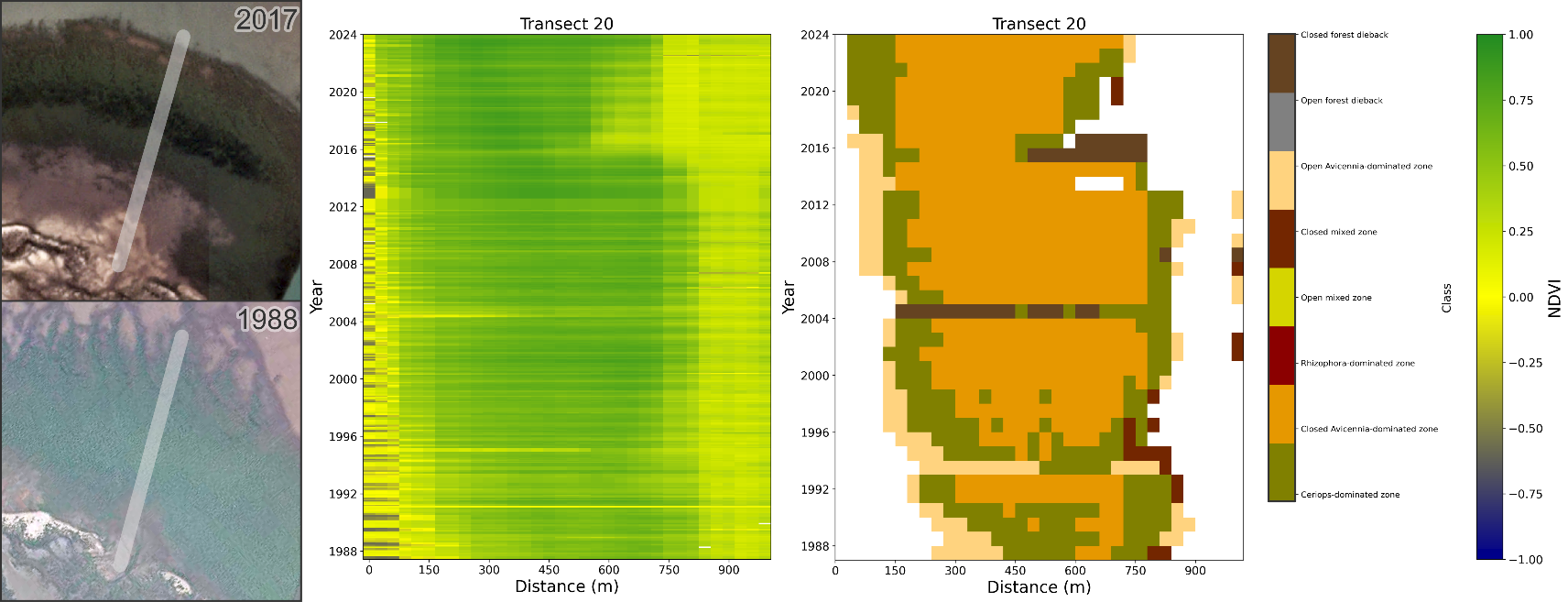

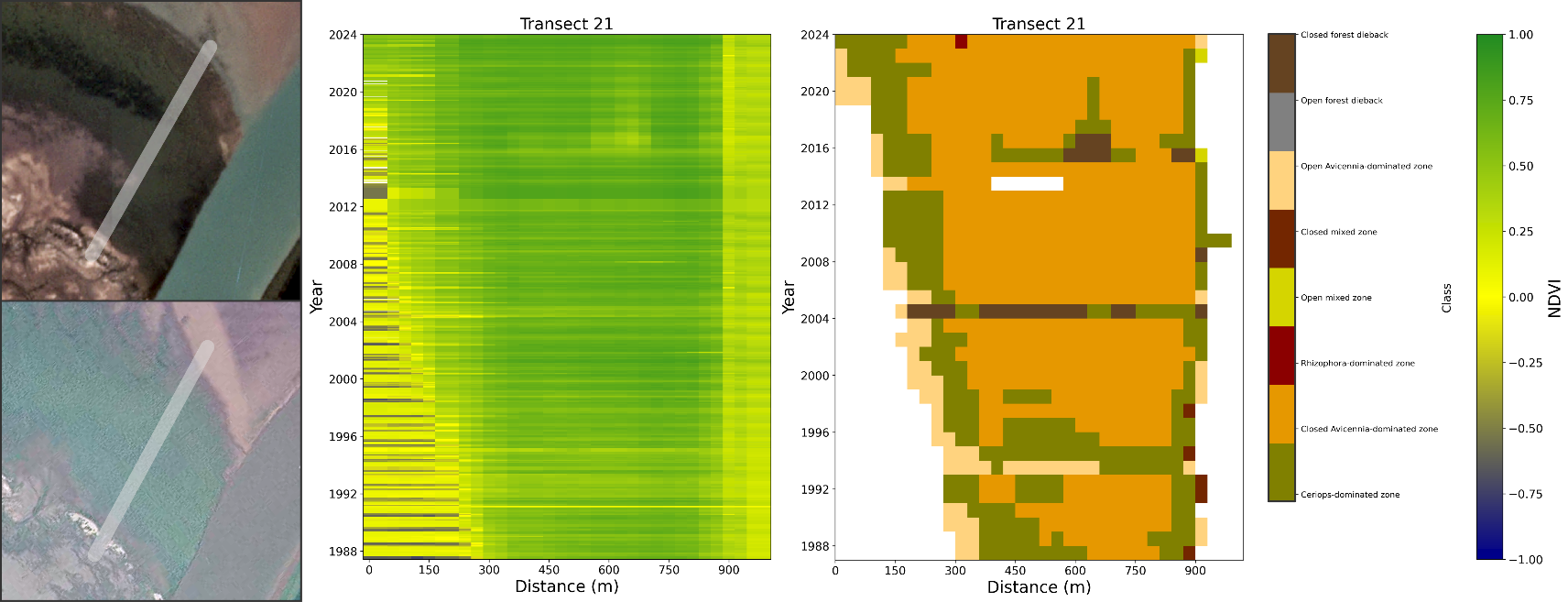

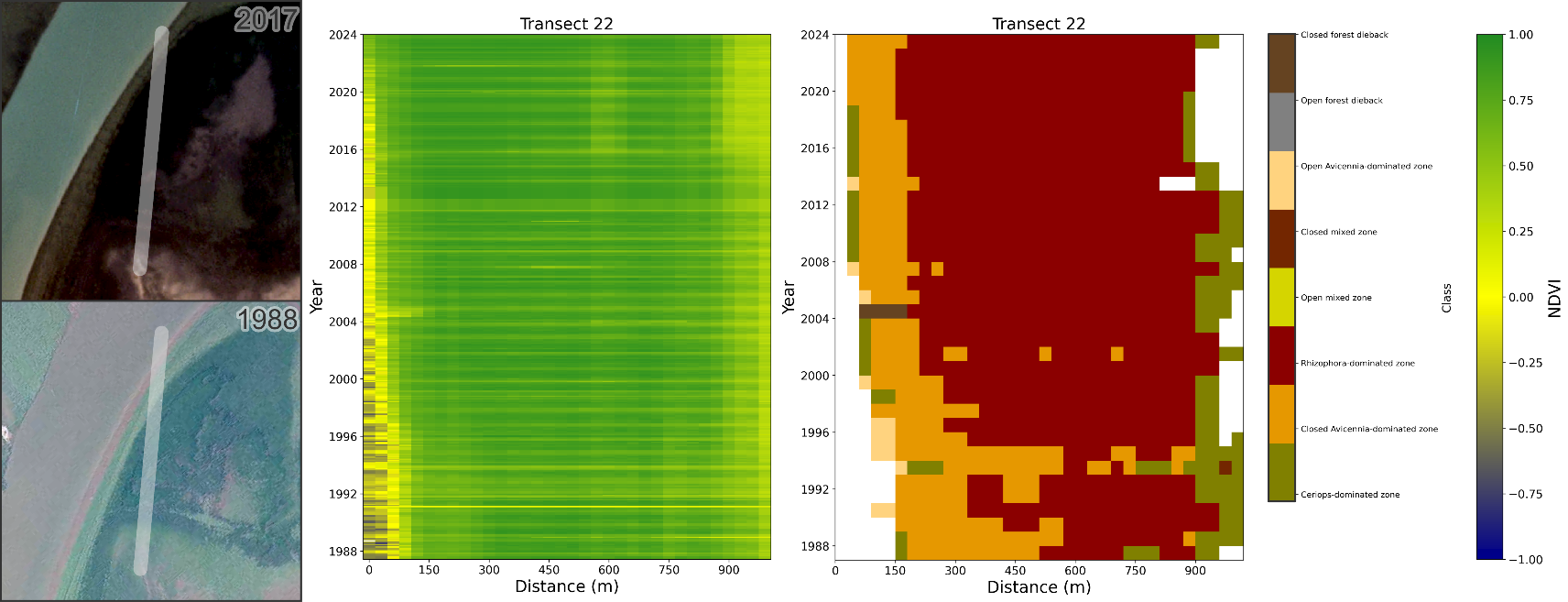

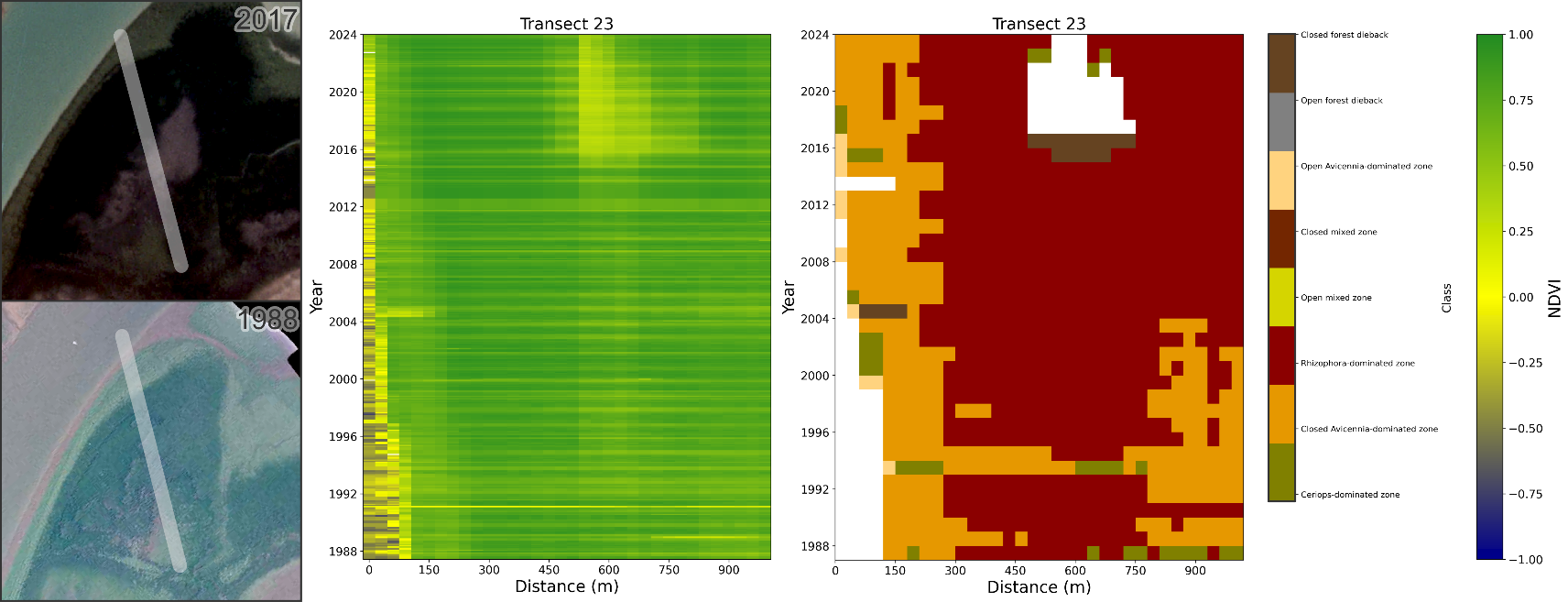

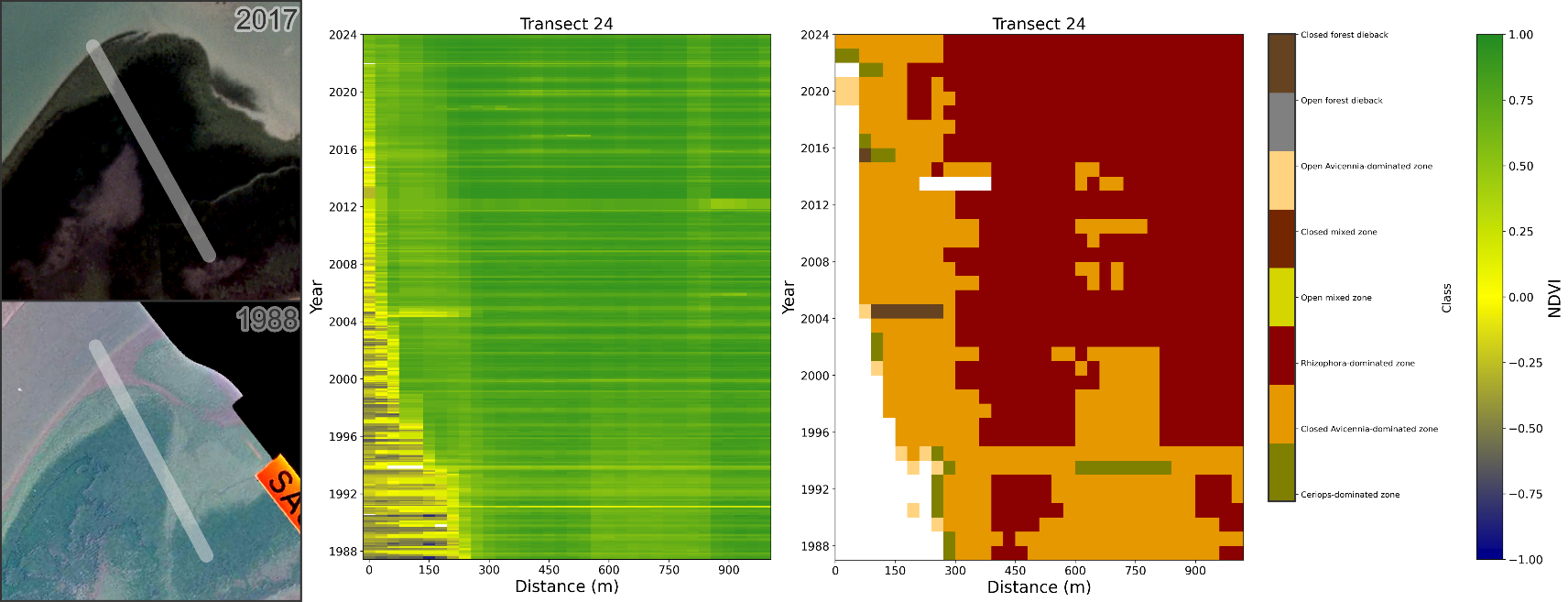

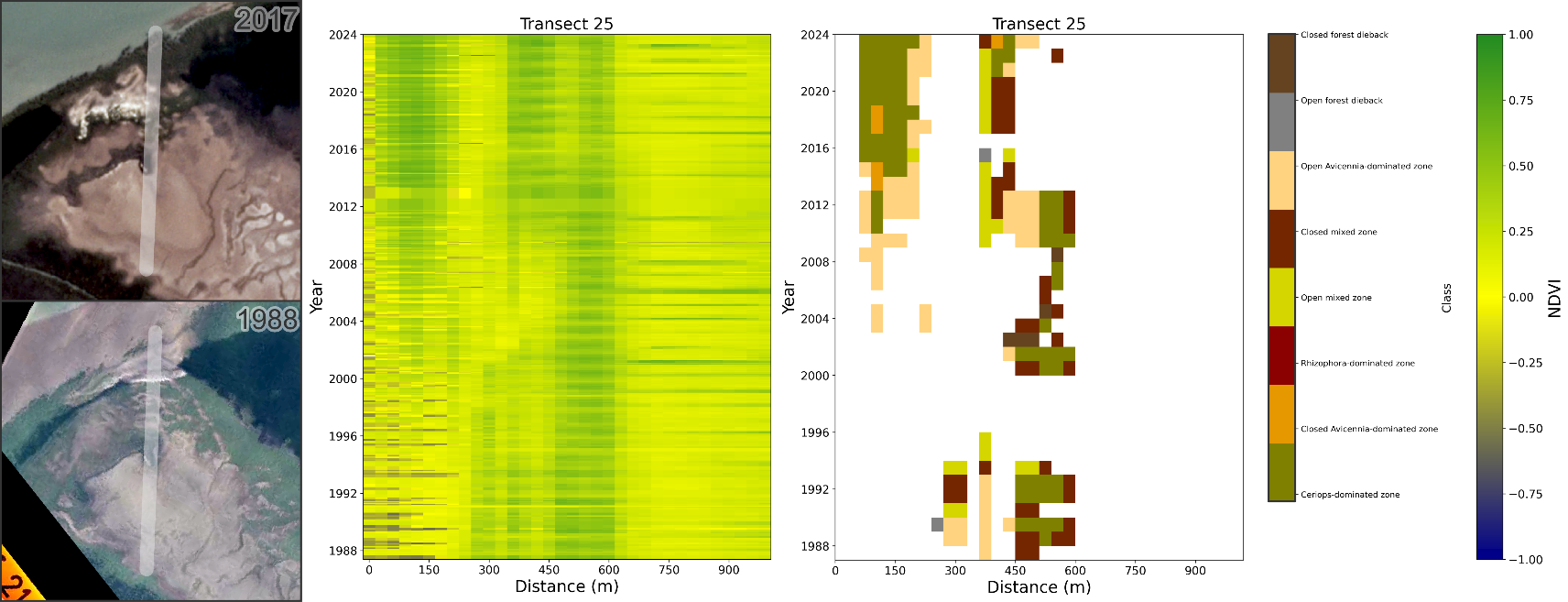

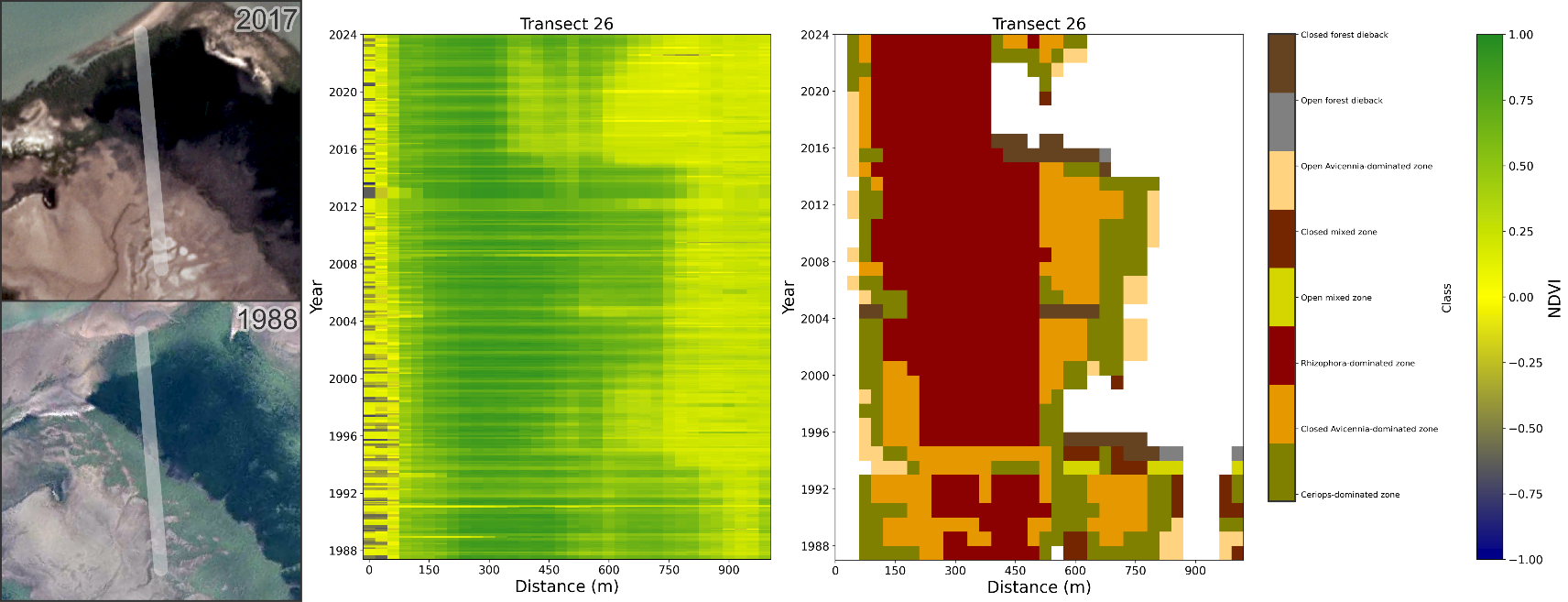

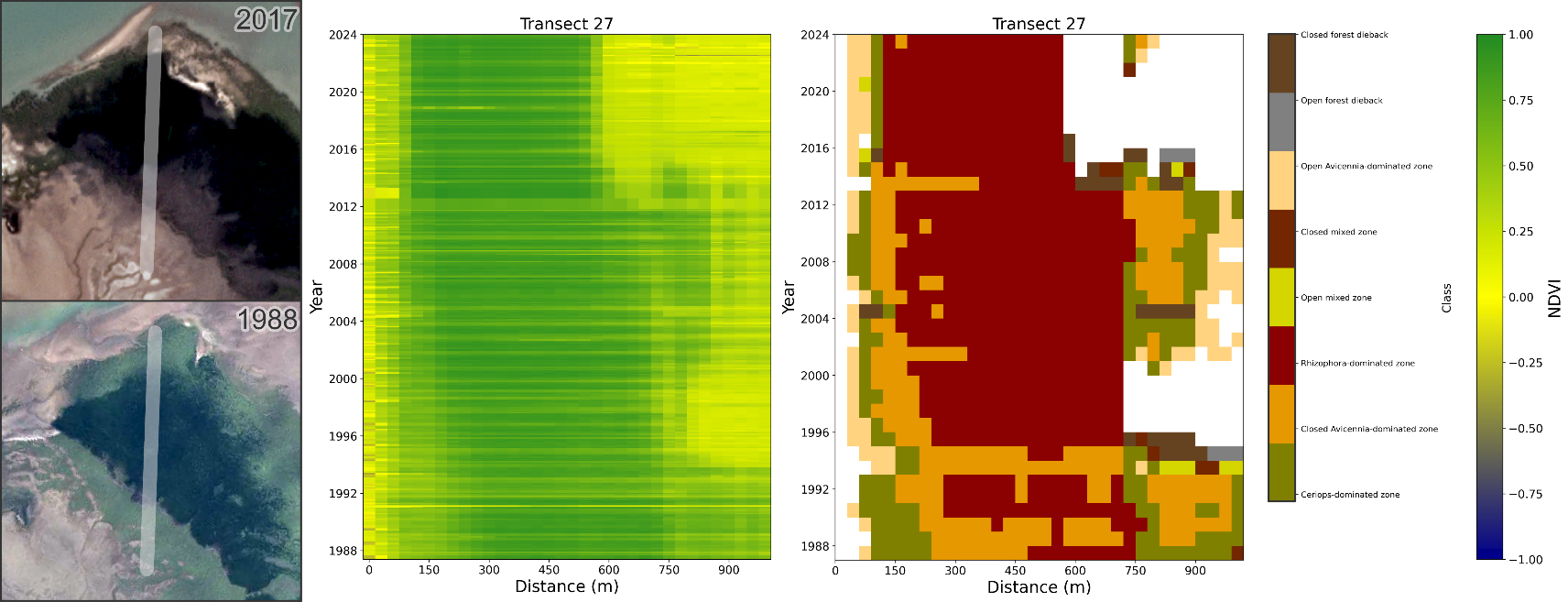

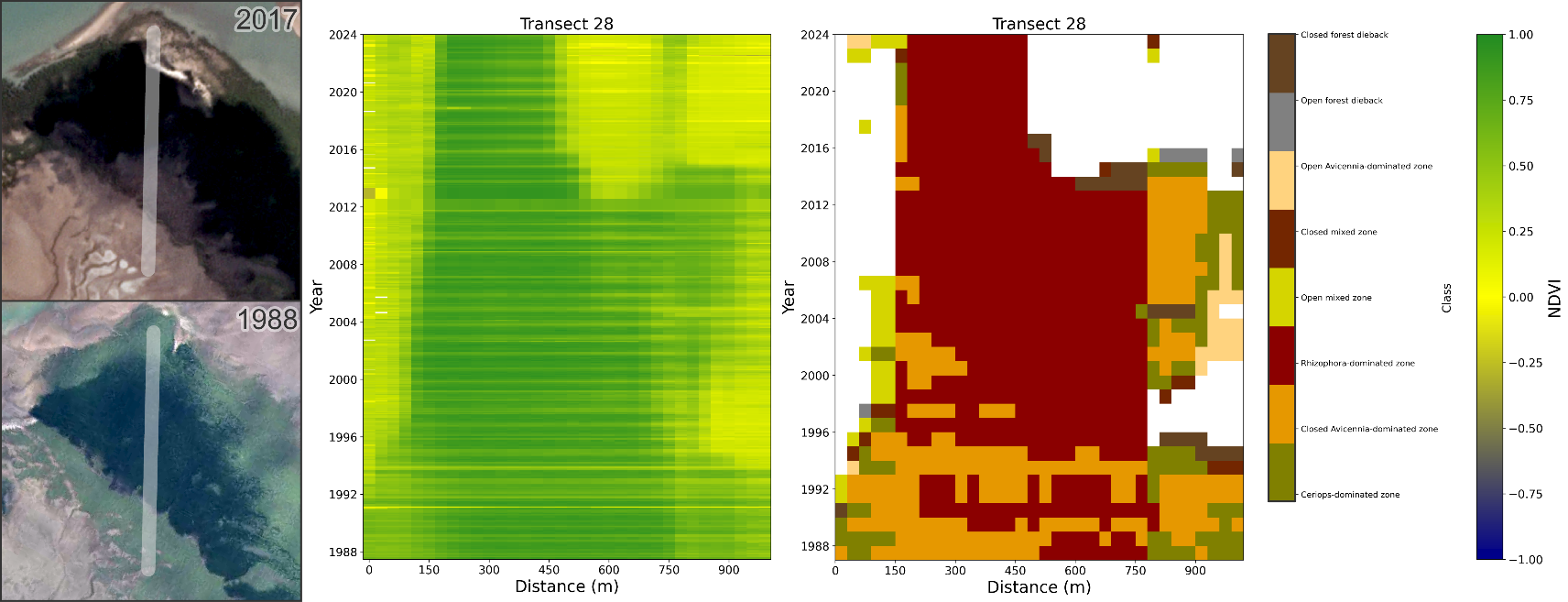

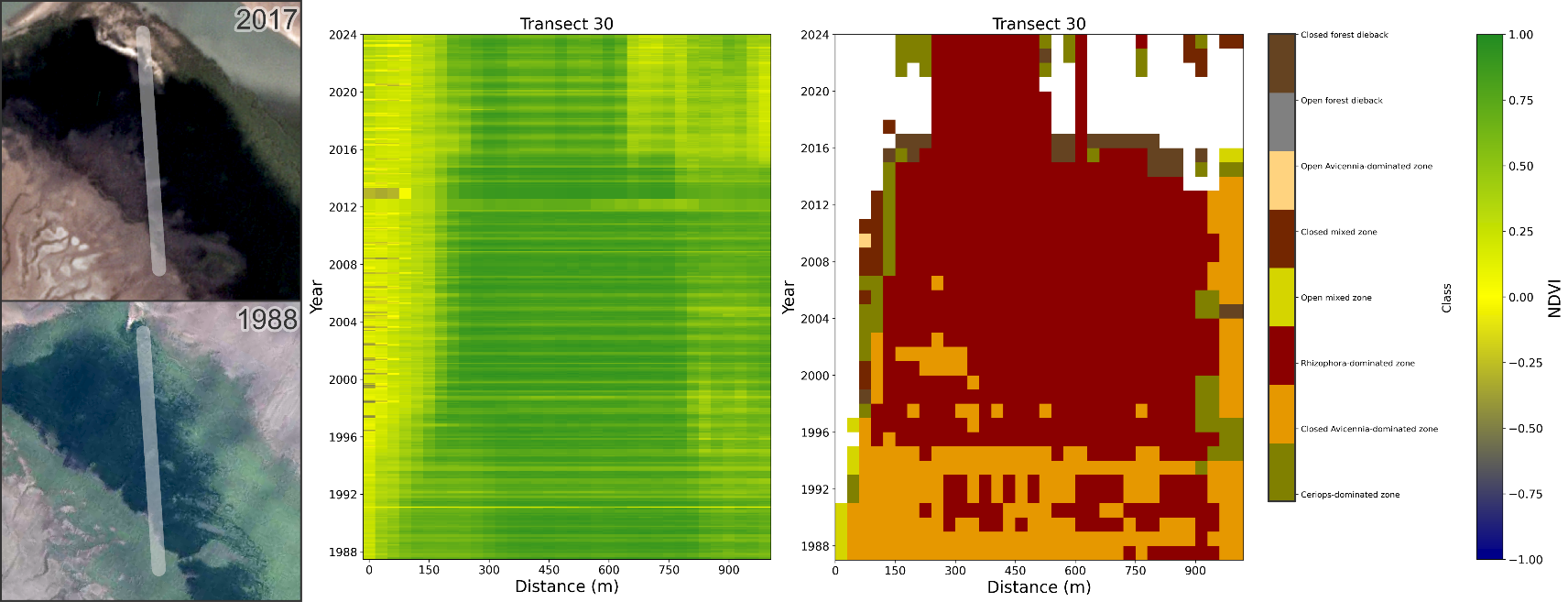

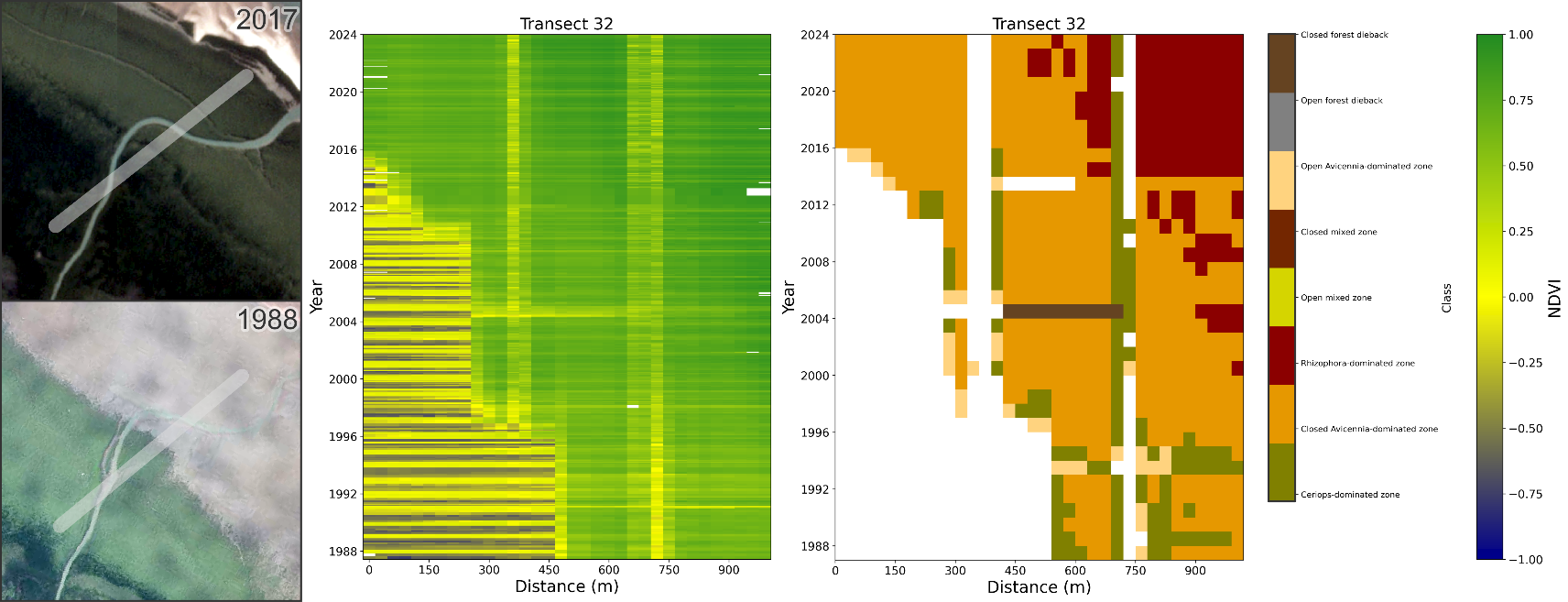

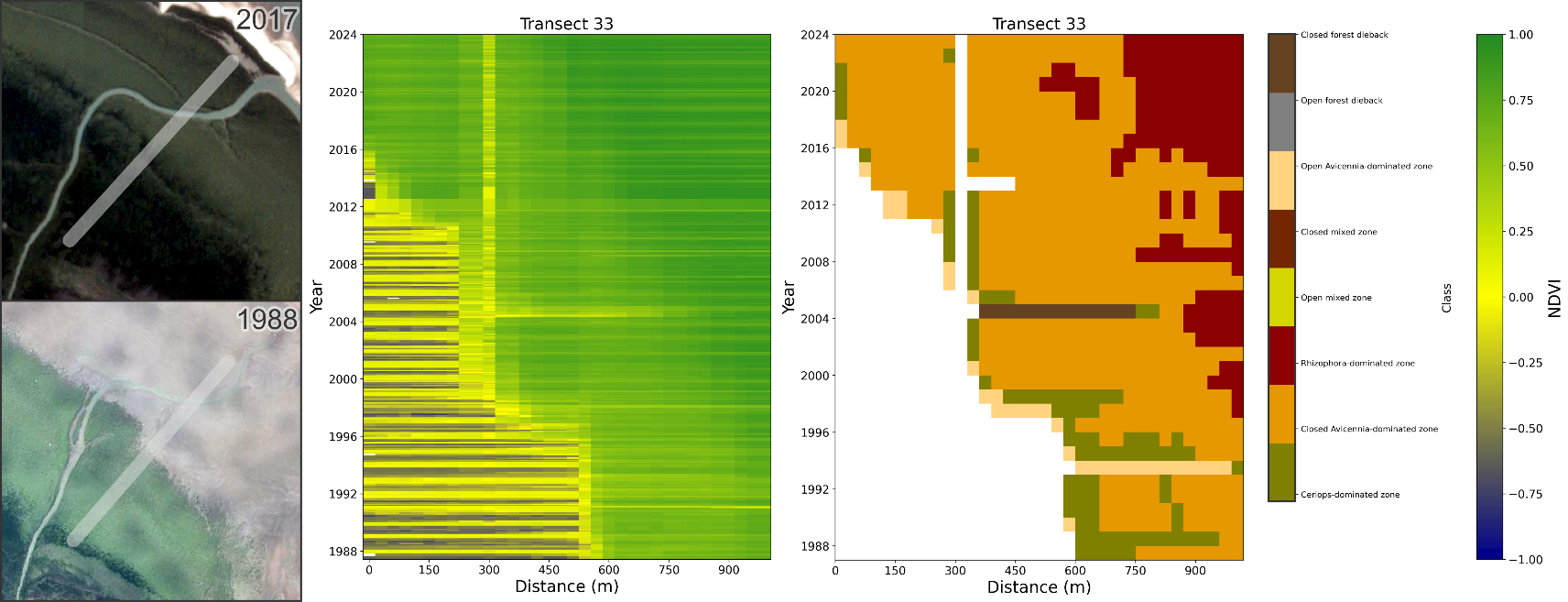

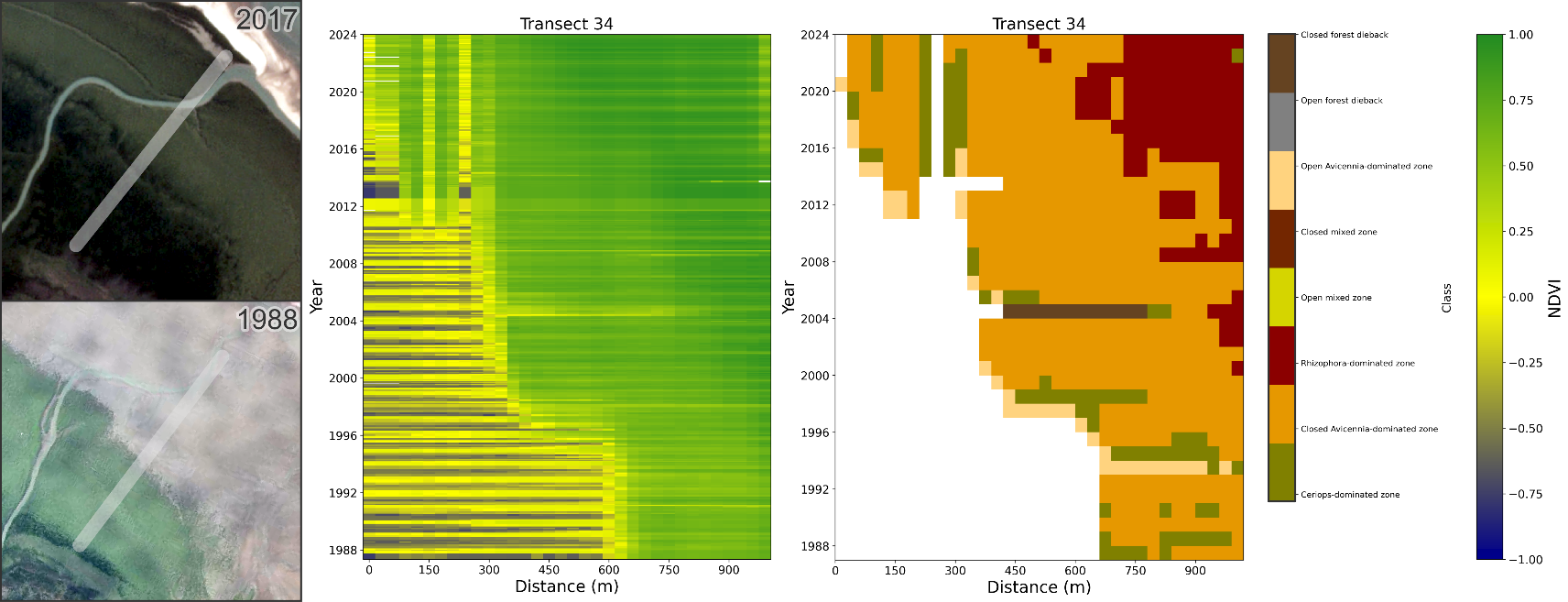

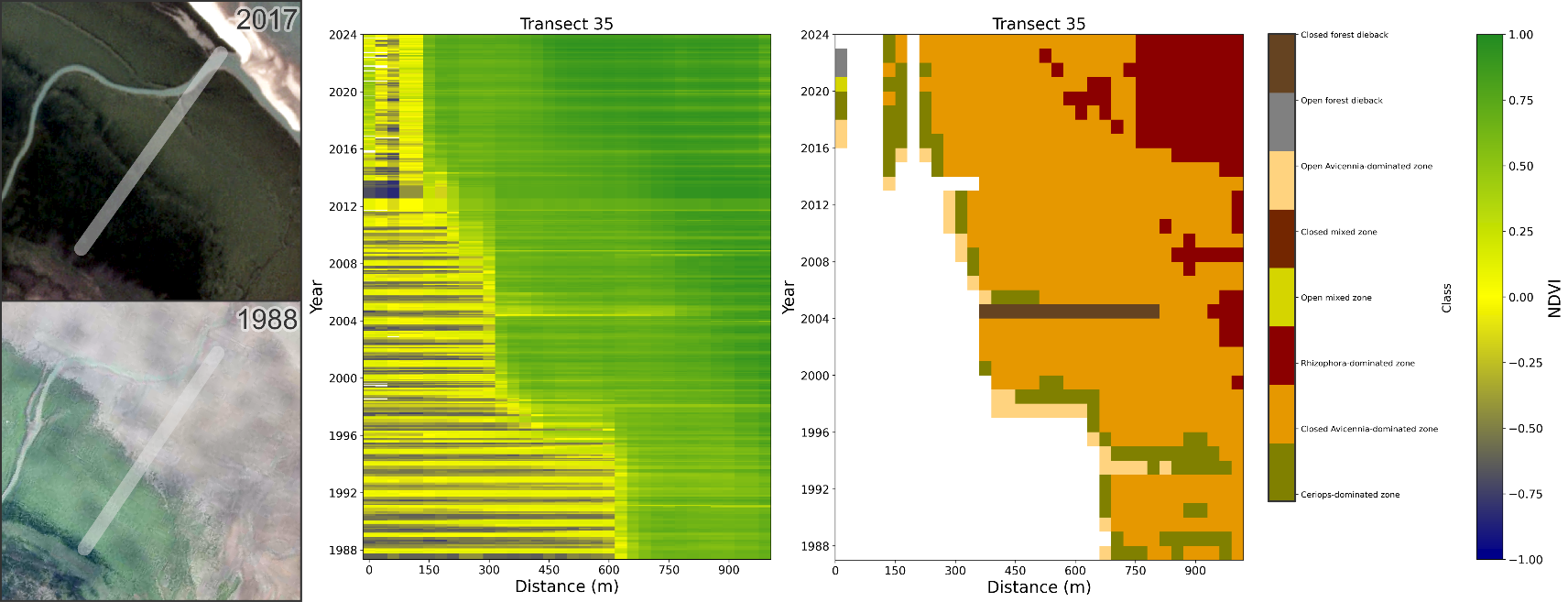

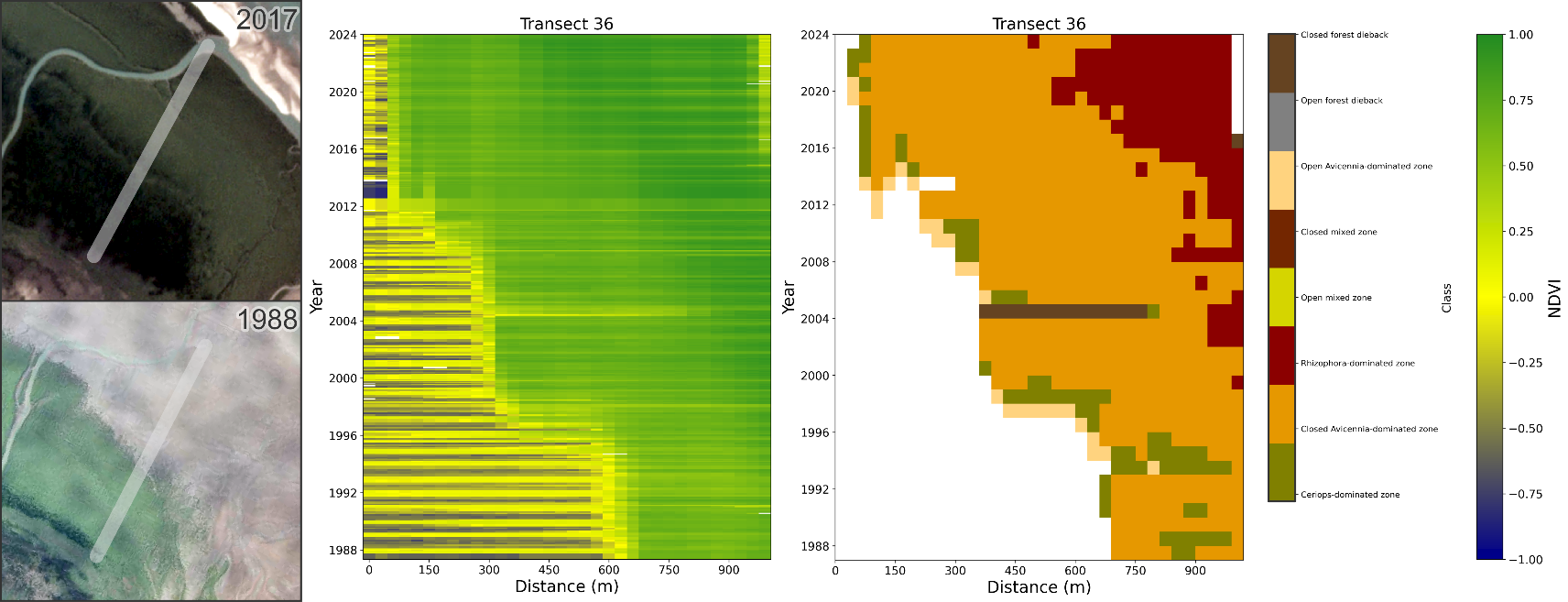

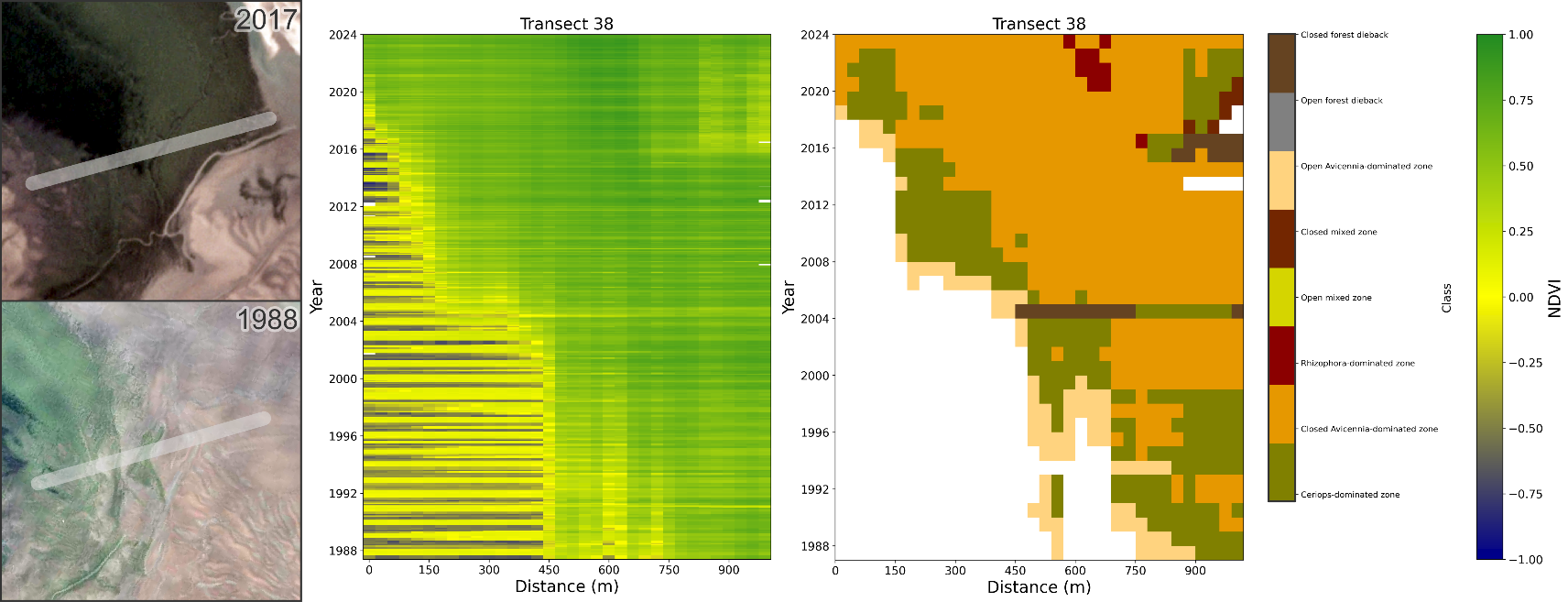

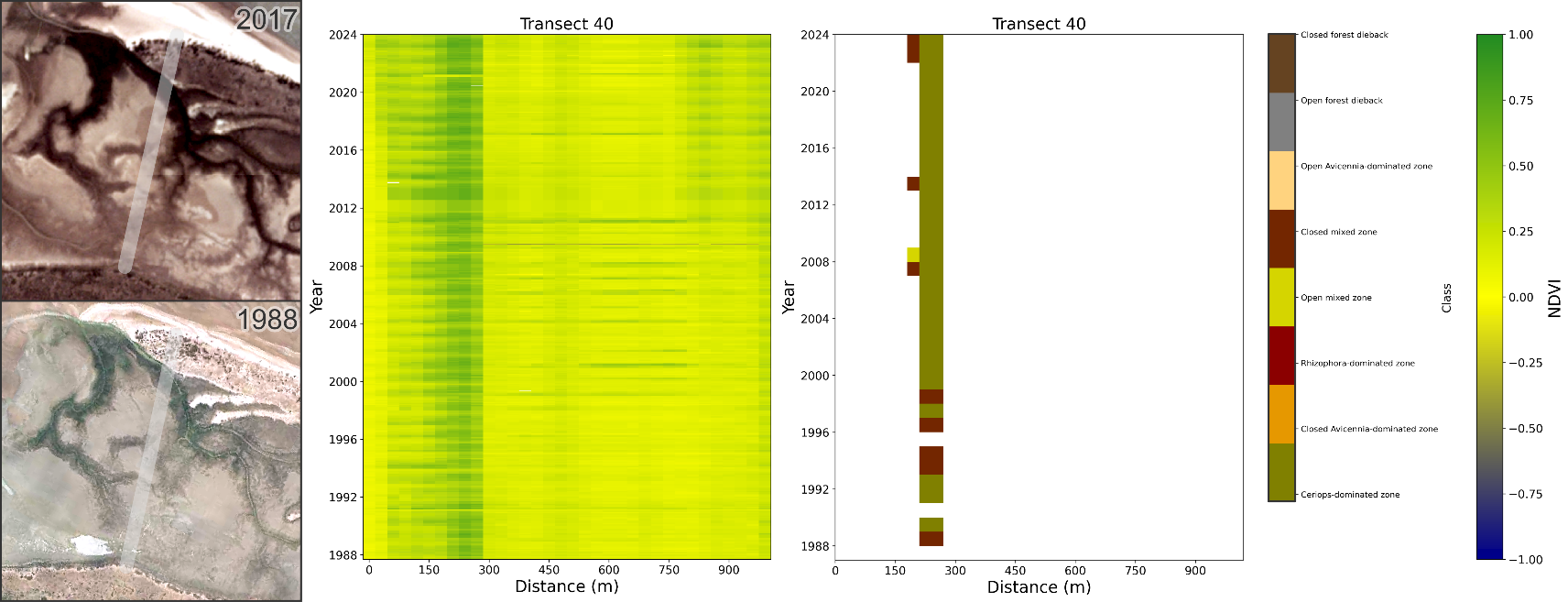

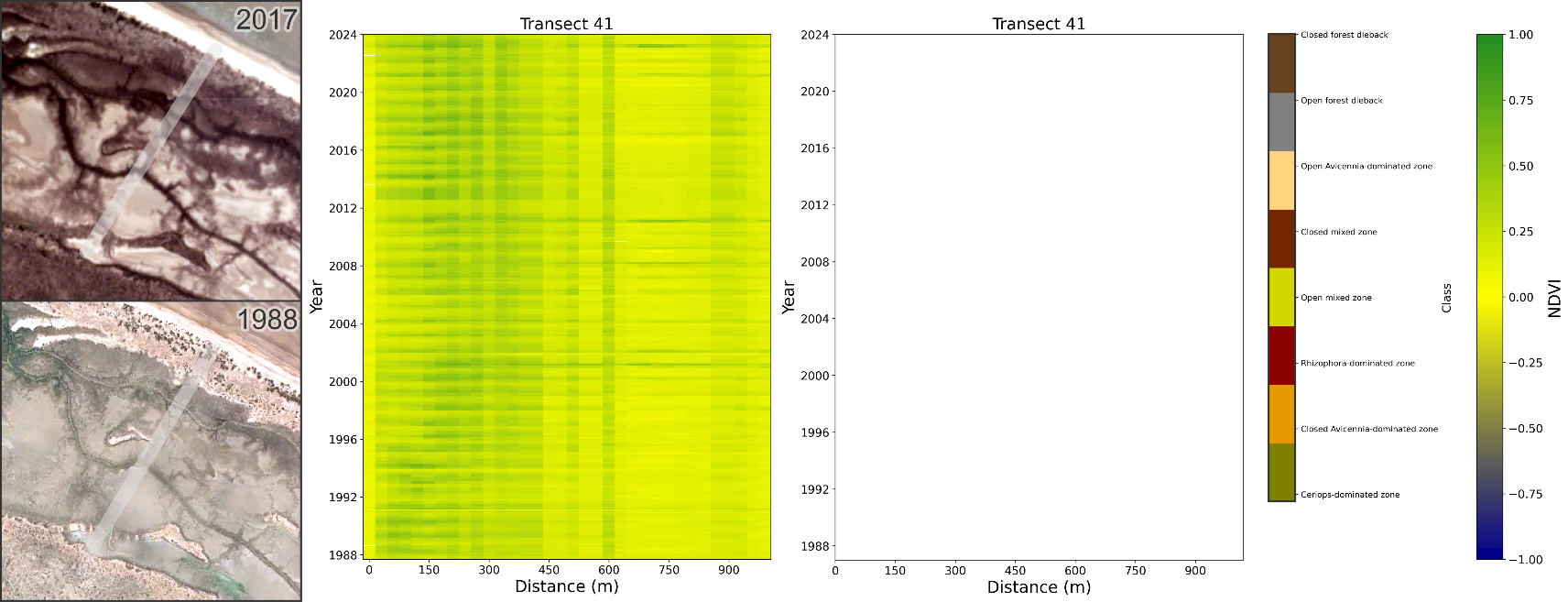

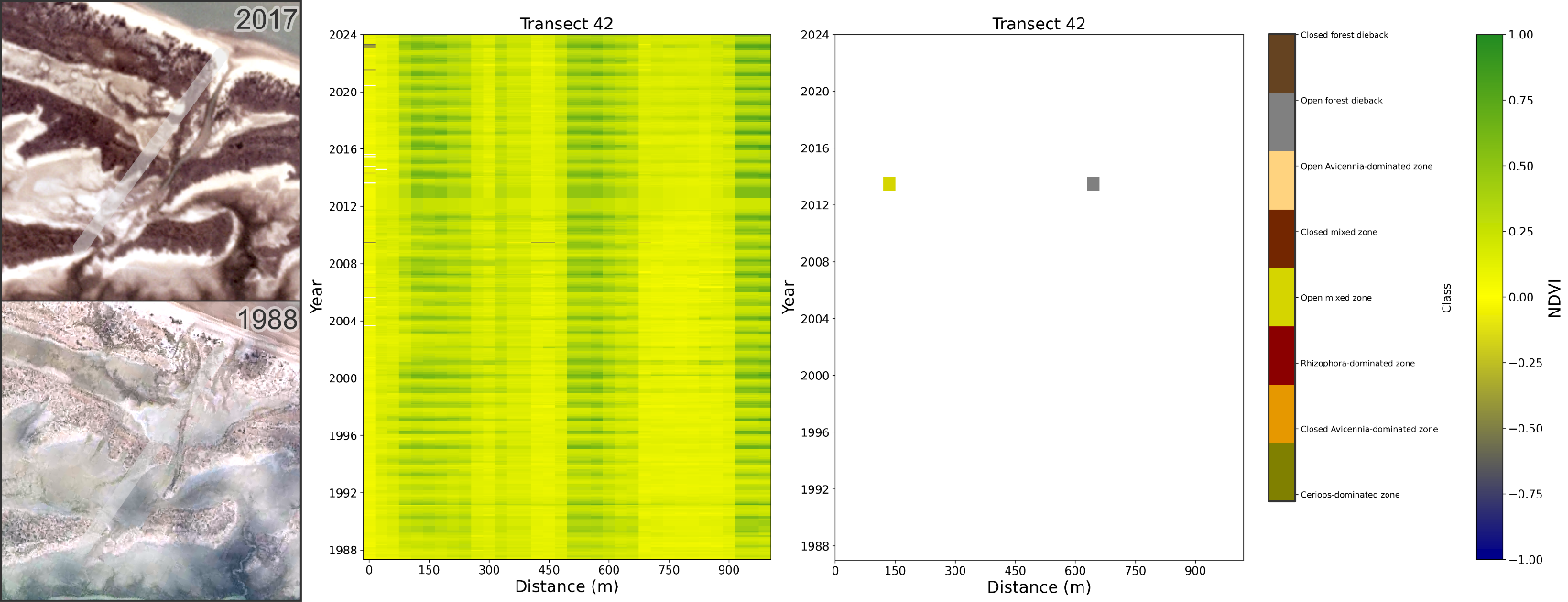

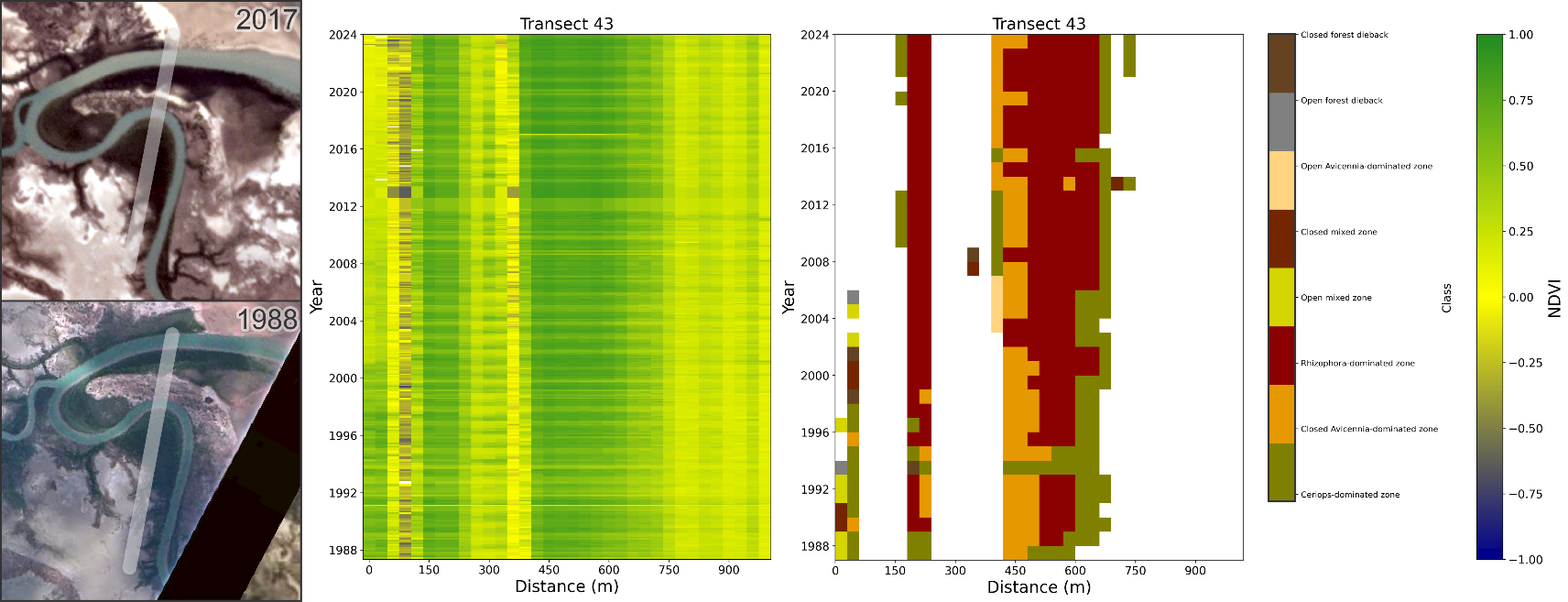

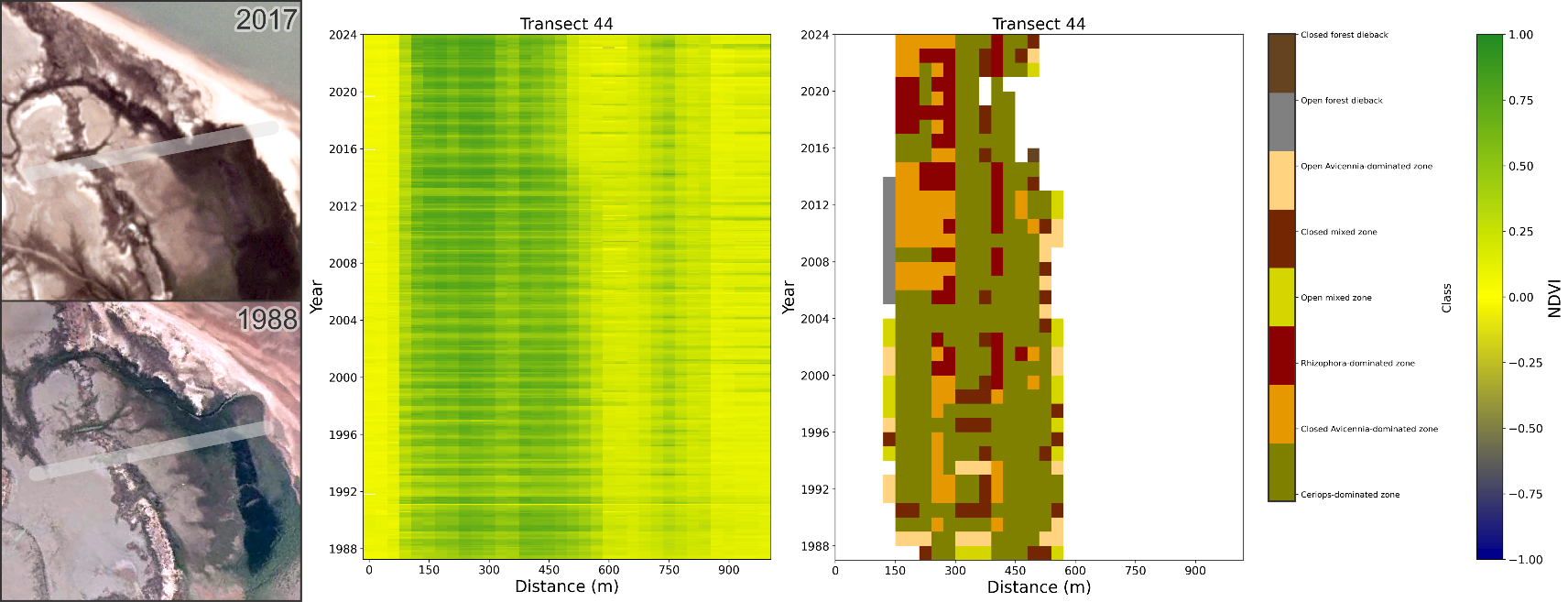

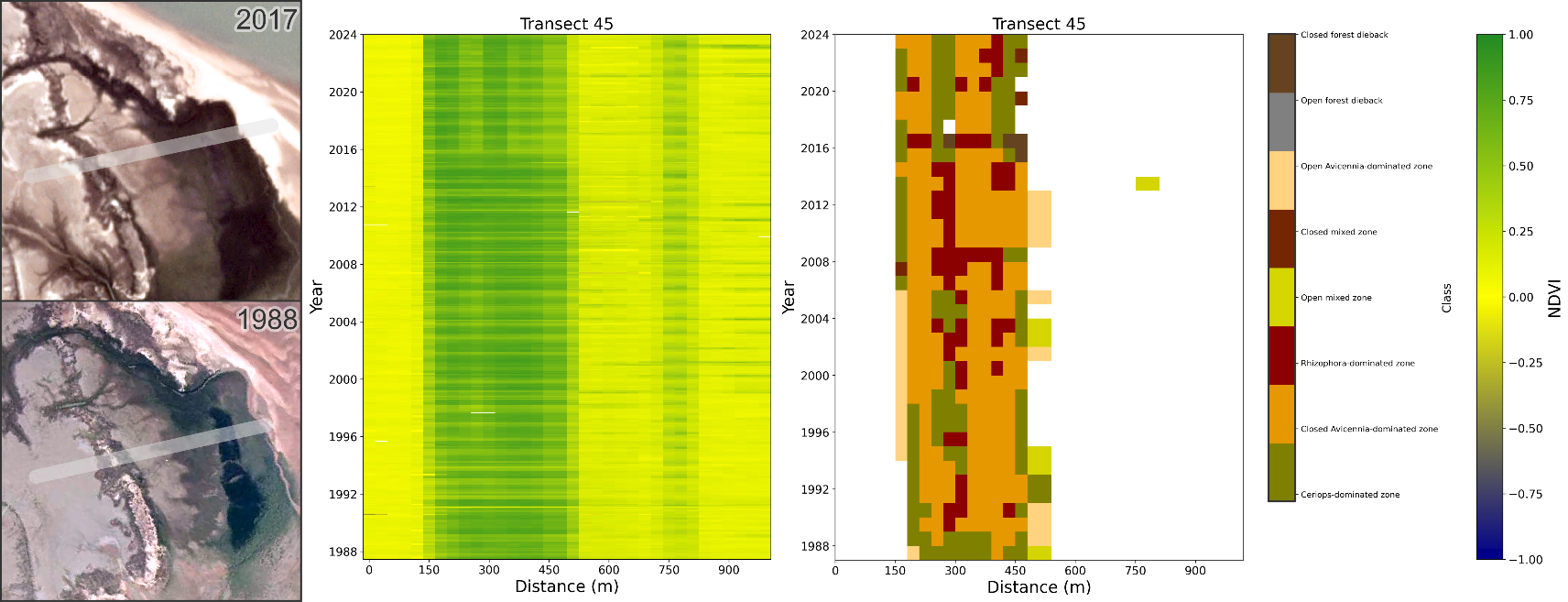

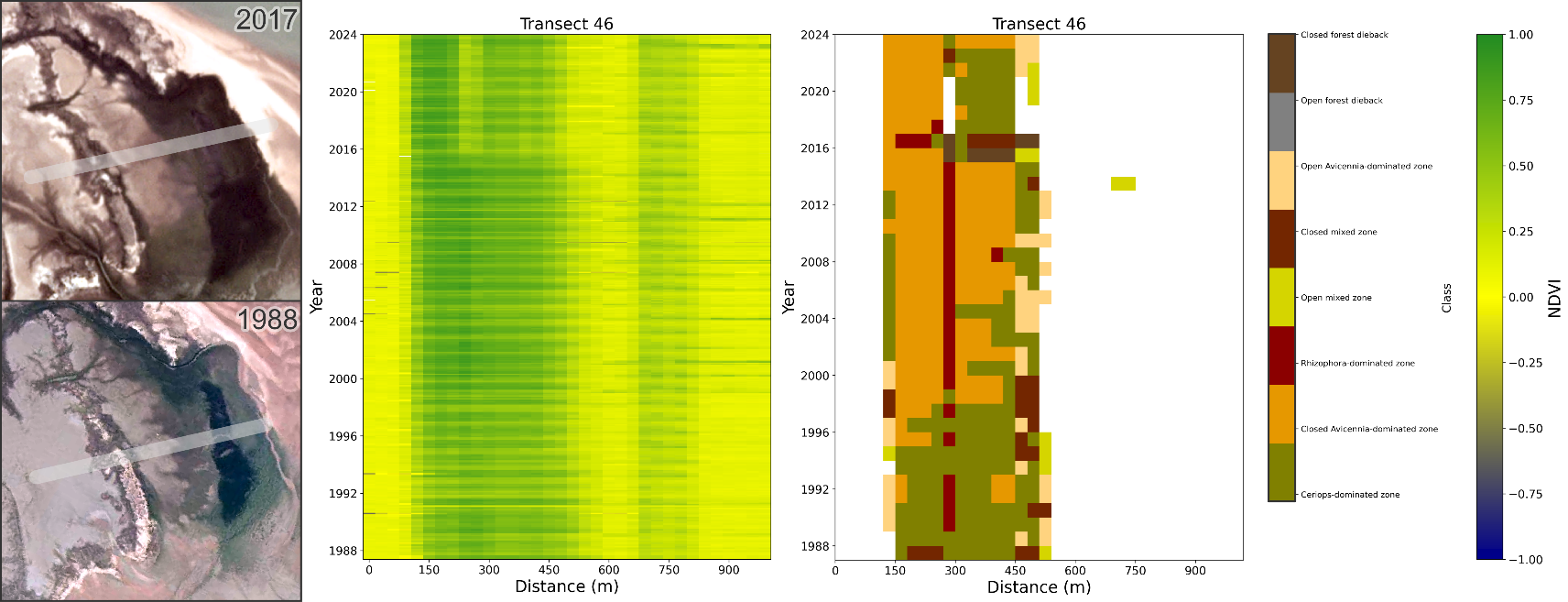

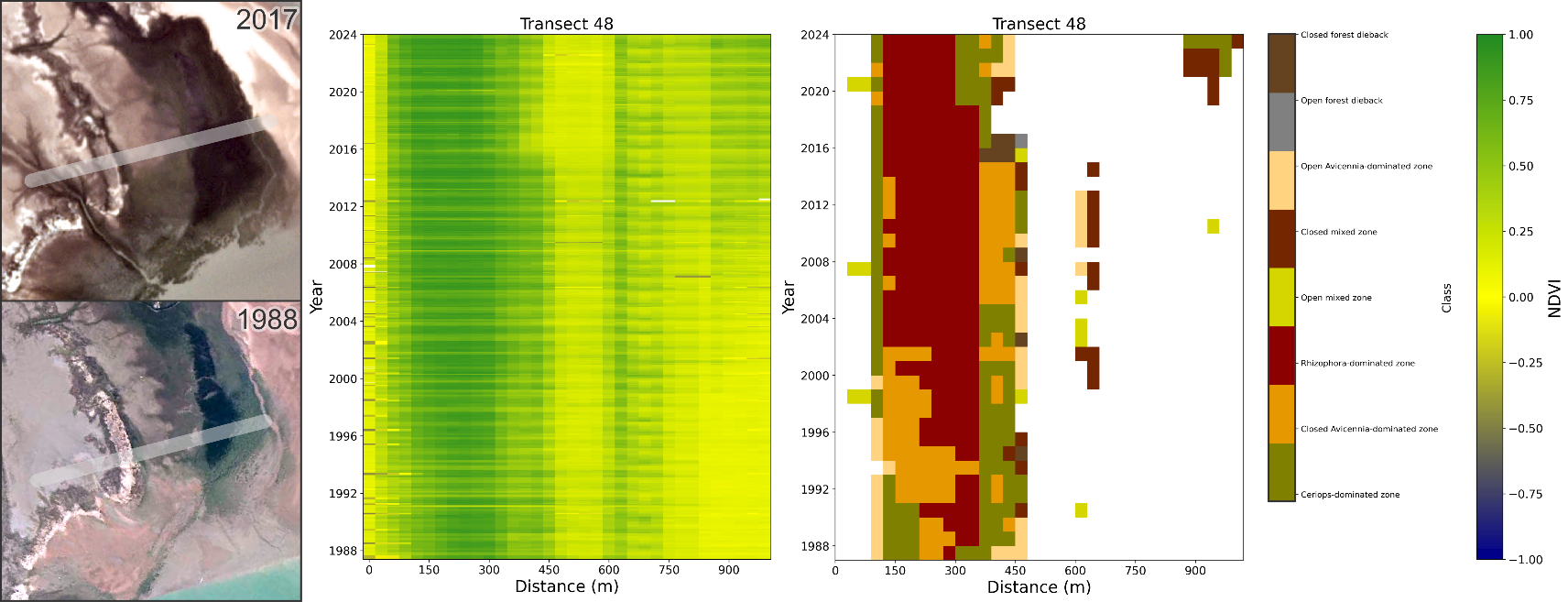

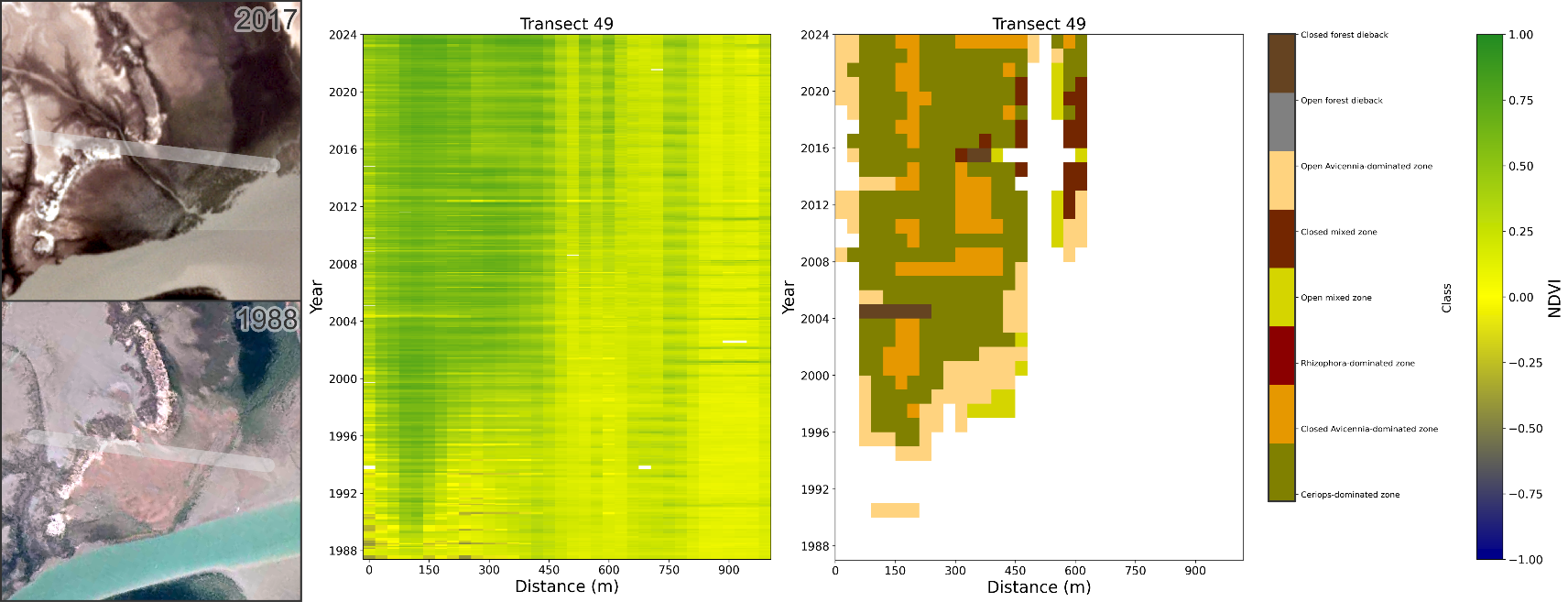

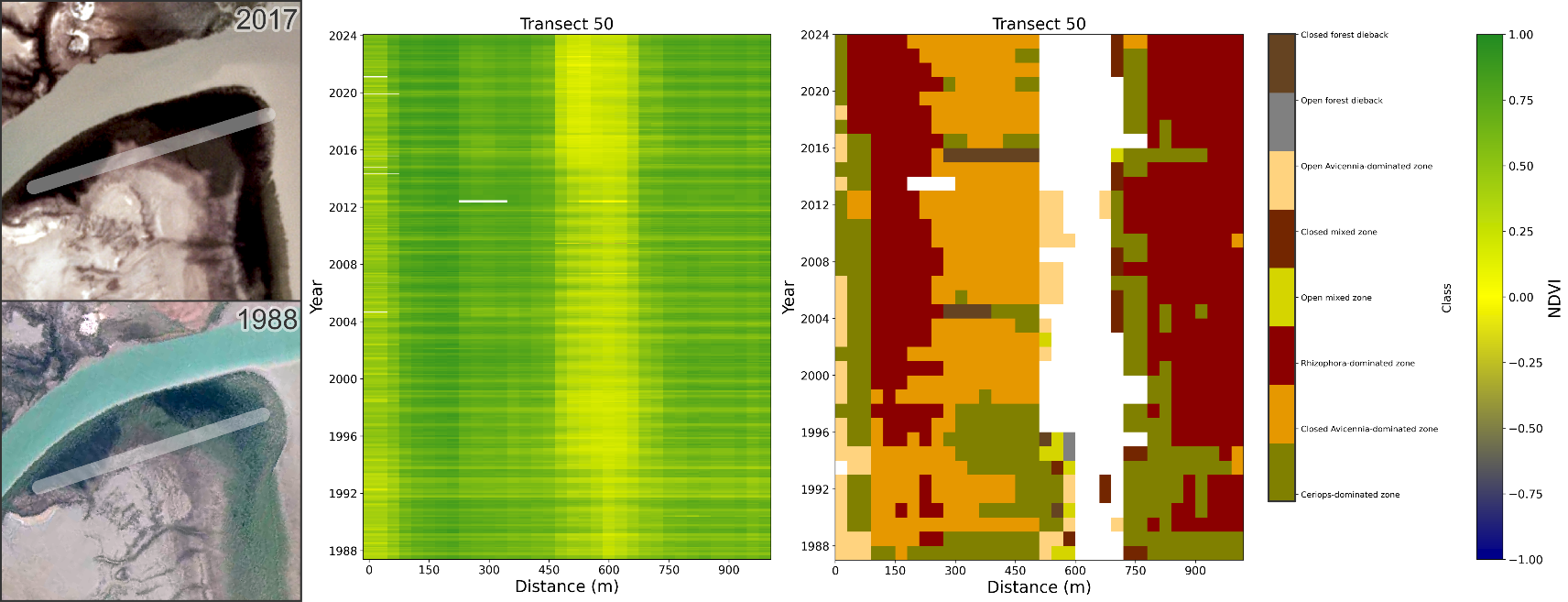

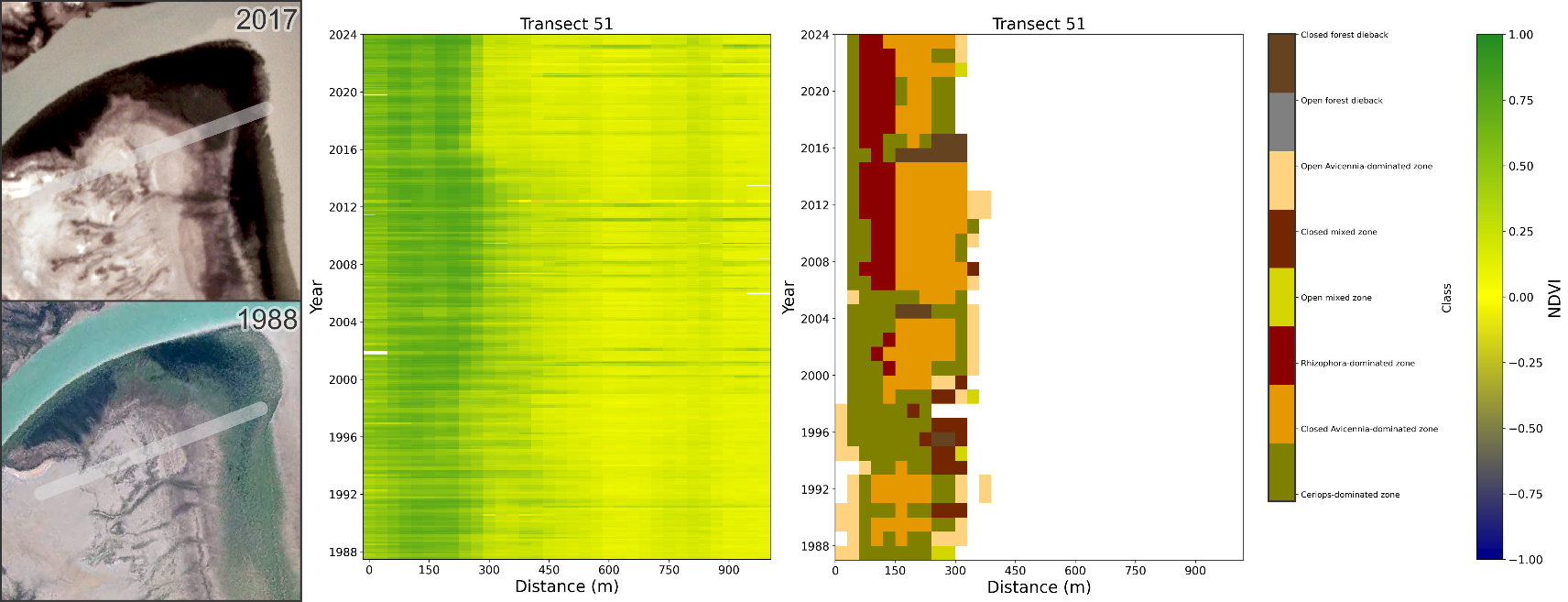

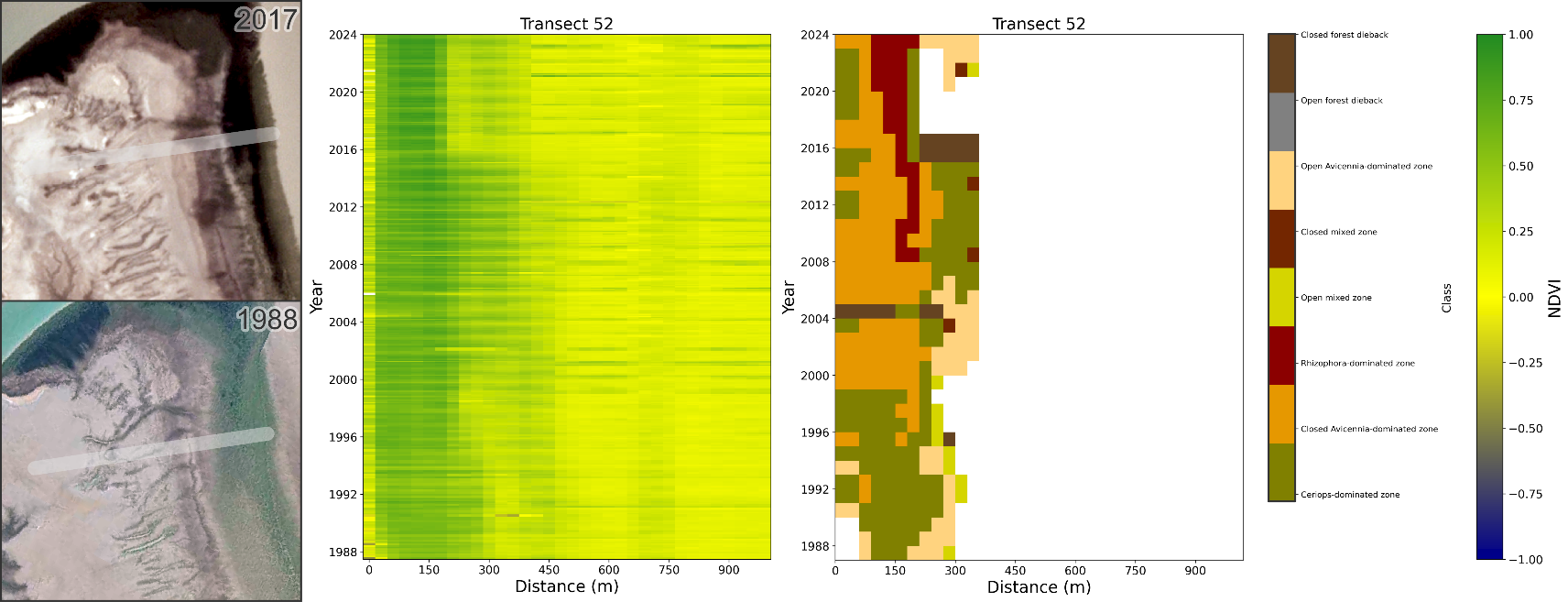

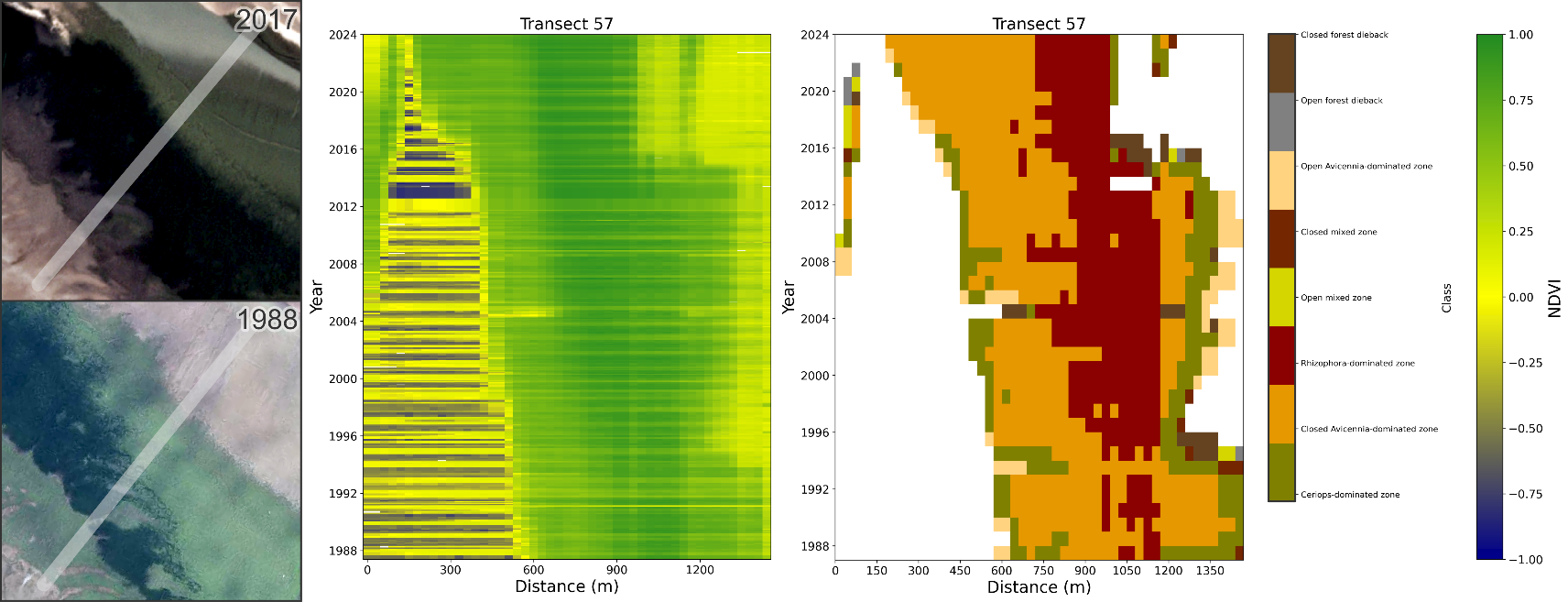
